# The Statistical Building Blocks of Animal Movement Simulations

**DOI:** 10.1101/2023.12.27.573450

**Authors:** Wayne M. Getz, Richard Salter, Varun Sethi, Shlomo Cain, Orr Spiegel, Sivan Toledo

## Abstract

Animal movement plays a key role in many ecological processes and has a direct influence on an individual’s fitness at several scales of analysis (i.e., next-step, subdiel, day-by-day, seasonal). This highlights the need to dissect movement behavior at different spatio-temporal scales and develop hierarchical movement tools for generating realistic tracks to supplement existing single-temporal-scale simulators. In reality, animal movement paths are a concatenation of fundamental movement elements (FuMEs: e.g., a step or wing flap), but these are not generally extractable from a relocation time-series track (e.g., sequential GPS fixes) from which step-length (SL, aka velocity) and turning-angle (TA) time series can be extracted. For short, fixed-length segments of track, we generate their SL and TA statistics (e.g., means, standard deviations, correlations) to obtain segment-specific vectors that can be cluster into different types. We use the centroids of these clusters to obtain a set of statistical movement elements (StaMEs; e.g. directed fast movement versus random slow movement elements) that we use as a basis for analyzing and simulating movement tracks. Our novel concept is that sequences of StaMEs provide a basis for constructing and fitting step-selection kernels at the scale of fixed-length canonical activity modes: short fixed-length sequences of interpretable activity such as dithering, ambling, directed walking, or running. Beyond this, variable length pure or characteristic mixtures of CAMs can be interpreted as behavioral activity modes (BAMs), such as gathering resources (a sequence of dithering and walking StaMEs) or beelining (a sequence of fast directed-walk StaMEs interspersed with vigilance and navigation stops). Here we formulate a multi-modal, step-selection kernel simulation framework, and construct a 2-mode movement simulator (Numerus ANIMOVER_1), using Numerus RAMP technology. We also illustrate methods for extracting StaMEs from both simulated and real data from two barn owls (*Tyto alba*) in the Harod Valley, Israel. Overall, our new bottom-up approach to path segmentation allows us to both dissect real movement tracks and generate realistic synthetic ones, thereby providing a general tool for testing hypothesis in movement ecology and simulating animal movement in diverse contexts such as evaluating an individual’s response to landscape changes, release of an individual into a novel environment, or identifying when individuals are sick or unusually stressed.

## 1 Introduction

One of the major challenges common to several subfields of ecology (e.g., conservation biology, disease ecology, resource ecology) is predicting how the movement of animals changes in response to landscape factors and the state or health of the individual [1, 2, 3]. Examples of change include the spatio-temporal distribution of resources vital to the existence of individuals [4, 3], the movement of animals released into new surroundings for the purposes of conservation [5, 2], or the movement of individuals under stress or with infections [6, 7]. The dynamic resource example has led to the concept of resource tracking, which has become an active field of research in movement ecology [8, 9, 10]. Quantification of this process through models that link animal movement to resources will enable us to better predict the consequences of global change on animal populations [11]. The stressed individual example may be critical to the continued existence of endangered species [12]. The third example may help us identify and selectively remove individuals that are sick, and hence reduce the risk of pandemic outbreaks [13].

Movement, whether simulated or real, generates complex patterns that require various approaches to classify and comprehend. The primary approach to deconstructing this complexity, which has been ongoing for at least 30 years [14, 15, 16, 17], has been to organize the movement track of animals into two or more different movement modes, and to parse the movement tracks of animals into consecutive segments each representing a different mode of movement from the previous segment. The primary quantitative methods that have been developed to carry out this type of path segmentation have been behavioral change point analysis (BCPA) [18, 19, 20, 21, 22, 23] and hidden Markov methods (HMM) [24, 25, 26, 27, 28].

An alternative approach to path segmentation is to view movement tracks as a hierarchical organization of segments with levels that have relevance at different spatiotemporal scales of analysis [29, 30, 31, 32] (Fig 1; also see Appendix A.1). The value of an hierarchical approach is abundantly evident as an epistemological tool for deconstructing complexity (e.g., the construction of texts and genomes), but requires some building-block basis (e.g., letters for texts, codons for genetic coding of proteins), whether real or imagined, for the hierarchical construction [33].

**Figure 1.**
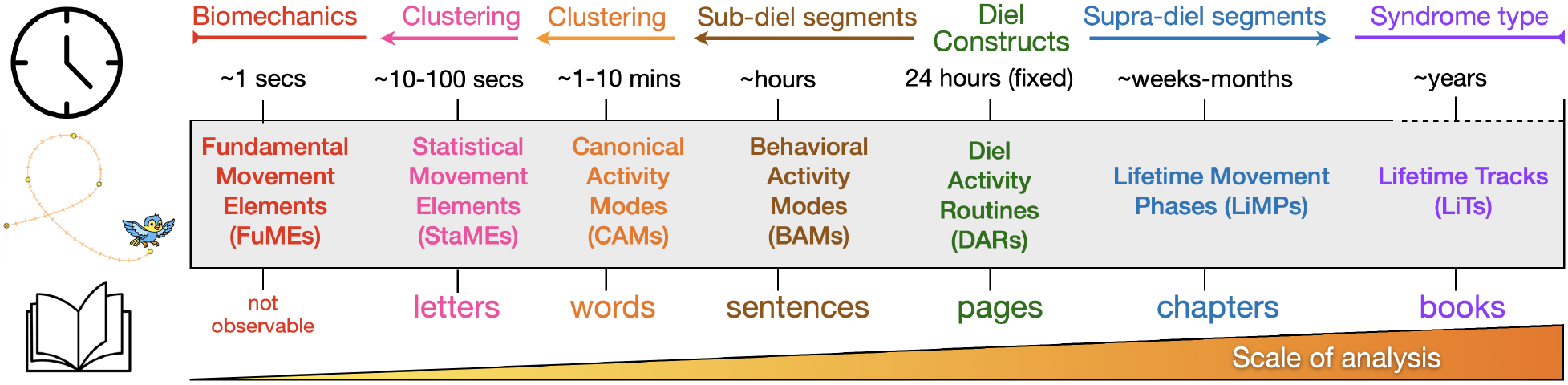
A facile comparison of a hierarchical movement track segmentation scheme (second ribbon from bottom) with hierarchical text elements (bottom ribbon) suggestive of the scheme’s utility to provide a movement track narrative of an animal’s life history. The listed time scale (second ribbon from the top) roughly applies to medium to large vertebrates. The top ribbon indicates that fixed-time DARs provide an anchor for the segmentation scheme, below and above which clustering and change-point methods of analysis can be used to respectively identify sub and supra diel segments (also see Appendix A.2). The StaME approach explored in this paper, provides a basic set of elements that can be used to hierarchically construct higher order elements, such as CAMs, BAMs, and DARs [34].

In the context of movement tracks, the real building blocks are fundamental movement elements (FuMEs; [30]—e.g., for horses this may be walk, trot, cantor and gallop, etc.), but these cannot be extracted from relocation data, even when such data have a resolution on the order of seconds. Identification of the FuME sequence as a animal moves through space is likely to require either analyses of videos of the movements, inferences using accelerometers data collected from particular locations on the animals body [35], or other types of data collected from sufficiently fast sensors to identify the start and end of each type of movement element (e.g. a wing beat of a bird [36]). In the absence of being able to identify the real FuME sequence, we propose the identification of statistical movement elements (StaMEs) as the smallest achievable building block elements for the hierarchical construction of animal movement tracks (Fig 1.

The purpose of this paper is to meet the following three goals:

a. Explore the potential of StaMEs as substitutes for FuMEs in providing a set of basic building block upon which next-level CAM segments of fixed size (number of steps), can be constructed and used to generate a further-level of variable-length behavioral activity modes (BAMs) (Fig 1)
b. Formulate a multi-mode canonical activity movement (CAM) framework, based on the implementation of step-selection kernels, with switching among kernels influenced by landscape structure (cellular arrays of resources and topographic measures), environmental variables (e.g., temperature, precipitation), and internal variables (e.g., surrogates for hunger, thirst, or diel schedules)
c. Provide a highly flexible, user friendly, freely available 2-movement-mode simulator in the form of a computer application package (Numerus Studio platform plus simulator application, both downloadable for free at links provided in Appendix C of the Supplementary Information File) and demonstrate its utility for movement ecologists to generate multi-modal movement tracks using step-selection methods and test hypothesis regarding mechanisms producing emergent patterns of movement.

This paper should be seen as part of a larger body of work that includes the formulation of a general framework for hierarchical track segmentation, as summarized in Fig 1 and discussed more fully in [30, 37]. In parallel, we are also generating measures that can be used to rigorously analyze bottom-up path segmentation methods using information theory measures of coding efficiency [34].

Elaborating on goal c.), our application is called ANIMOVER_1 (**ANI**mal **MOVE**ment **R**AMP; the “1” anticipates future elaborated versions) and the acronym RAMP refers to Numerus’ highly flexible **R**untime **A**lterable **M**odeling **P**latform technology [37]. We stress that this application platform is not a general programming environment and does not require any coding experience. However, it allows the user to set parameter values and, if desired, to overwrite critical lines of default code pertaining to landscape generating algorithms and agent movement rules to meet the user’s specific needs.

Empirically, the movement track of an individual over a landscape is generally represented by a sequence of locations that is recorded using GPS technology [38], ATLAS reverse GPS technology [39], acoustic receivers or other technologies [40]. From such sets of relocation points, also referred to as a “walk,” step-length (SL; also velocities when the sampling frequency is fixed) and turning-angle (TA) time series can be extracted [30] (Appendix A.2). These time series can then be used to compute various derived quantities, such as radial and tangential velocities at each relocation point, and auto correlations of variables along segments of the movement track [18]. The statistics of such variables, computed for fixed short segments of track (e.g., 10-30 points), can then be used to categorize such segments into statistical movement elements (i.e., StaMEs previously called referred to as metaFuMEs in [30]).

These StaMEs can then be classified into a limited number of categories, as demonstrated in this paper (e.g. short elements underpinning direct fast flight, brisk walking, meandering, and so on). A string of same category StaMEs then constitutes a track segment that can be classified as a homogeneous or canonical activity mode (CAM) of a type defined by the underlying category of StaME (e.g., brisk walking might translate into bee-lining and meandering into searching behavior). Characteristic mixtures of CAMs, in turn, can be strung together into identifiable behavioral activity modes (BAMs; e.g., resting, foraging, heading to a known location while being vigilant), with several BAMs coming together each day to form a diel activity routine (DAR) [41, 42] (Fig 1. The DAR itself is a hierarchical segment that can be understood in terms of an invariant 24-hour period for most animals, apart from some deep dwelling marine or cave-dwelling species, because for most species it is a fundamental biological rhythm honed by evolution [43]. The periods of various exogenous environmental cycles around or beyond the diel period (e.g., lunar, seasonal, or tidal), though, can be quite variable in their effects on species, depending on the latitude [44], elevation [45] and the trophic levels (e.g., herbivore, predator, scavenger) at which they function.

If the relocation sampling frequency is relatively high (i.e., ≈ 5 or more relocation points per min), then the statistical properties of a segment of, say, 10-30 consecutive points (e.g., the means of the velocities and others) can serve to construct a set of StaMEs, which may then be classified into a relatively small set of StaME categories and associated canonical activity modes. From our presentation here, it will become clear that StaMEs are dependent, firstly, on the resolution (i.e., frequency) of the relocation data and, secondly, on the number of consecutive points used to derive the statistics of our StaMEs (i.e., its duration). Since some of the measures and features used to characterize movement track segments using relocation data are noticeably frequency dependent [46, 15], these will be influenced by the scale at which we define the underlying set of StaMEs used to reconstruct movement track segments of various lengths.

Movement, of course, does not occur in a vacuum and the statistics of the movement elements are going to be affected by landscape factors (e.g., slope and roughness of the terrain) and various environmental conditions (e.g., resources, temperature, wind, ocean currents, etc.) [47, 48, 49]. The effects of such factors will induce additional variation or noise in the statistics of StaMEs of different types. If all we have is a movement relocation time series without the benefit of covariate variables to provide context, then ignoring such covariates will add some noise to the process of identifying an underlying set of movement track StaMEs. If such data are available, then different sets of StaMEs can be identified for segments occurring for particular ranges of covariate values. Otherwise, modifications to the StaME statistics can be made under the assumptions that for each type of StaME identified for the track as a whole, modifications are needed when particular covariate conditions exist (e.g. terrains that exceed a steepness threshold or when prevailing winds exceed a wind velocity threshold).

Finally, in this paper, we discuss why the construction of our ANIMOVER_1 RAMP facilitates analyses of movement pathways by ecologists compared with the effort needed to code up movement simulations from scratch using current programming platforms, such as Netlogo [50] or R. In addition, we focus on issues relating to parameter selection and running the model, as well as classifying StaMEs derived from both simulated and real data into a limited number of types (in our case 8 categories using hierarchical clustering methods). The real data relate to the movement data of barn owls (*Tyto alba*) in the Harod Valley, Israel, as more fully discussed in other contexts elsewhere [42, 51, 52].

## 2 Model Framework

### 2.1 Arrays and movement kernels

In this paper, we focus on two dimensional models, which of course is only appropriate for some species (e.g., most terrestrial species) but not others (e.g., deep diving marine species or birds use thermals to gain height). For some birds, such as the barn owl data analyzed in Section 5.3 of this paper, accurate tracking in 3 dimension is infeasible and the height component of their flight is much smaller than the surface two dimensional components of their flight. Our model is thus implemented on a landscape represented by an *n*^row^ × *n*^col^ cellular array such that cell(*a, b*) has value *c*_*ab,t*_ at time *t*: i.e.,

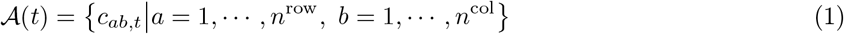

Also, the topology of this array can be selected to be `topology` = `torus` (top-bottom and left-right continuity identification) or `plane` (top, bottom, left and right boundaries).

Each cell is identified both by its location (*a, b*) in the array, and by Euclidean coordinates 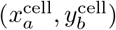 at the lower left corner of cell(*a, b*). Depending on the units used to measure our Euclidean landscape, we define increments Δ*x* and Δ*y* such that *n*^row^Δ*x* and *n*^col^Δ*y* provide the desired dimensions for our landscape. Scaling is important when considering how far individuals are likely to move in one unit of time when in different movement modes and thus the scaling of time is linked to the scaling of space in real applications. In theoretical studies not linked to empirical data, however, it will be convenient to set Δ*x* = Δ*y* = 1 and to set a parameter Ω_scale_ = 10 to scale space with respect to time such that the greatest distance an individual is likely to travel in one time step is given by Ω_scale_Δ*x*.

The model simulates movement of an individual over this landscape, with one step executed at each tick *t* of the simulation clock for *t* = 0, …, *n*^time^. If an individual is at a point 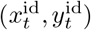 at time *t* and moves to cell(*a, b*), where its new location is now 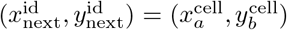, then the distance moved, denoted as 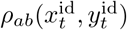 is defined as

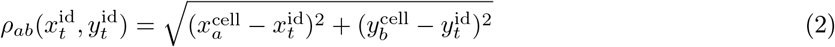

The angle of heading, denoted as 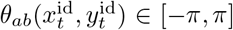 as measured from the positive horizontal (i.e., the axis *x* ≥ 0), is defined in terms of the so-called atan2 function as

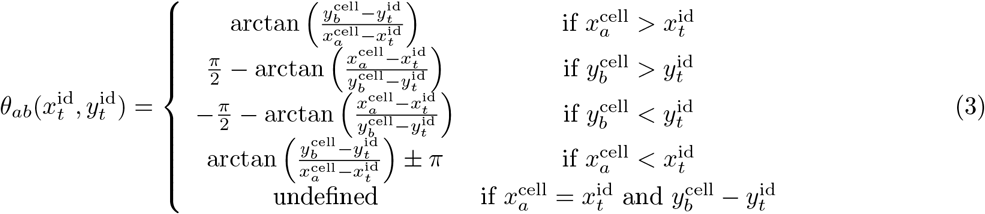

Whenever the angle of heading is reported as ranging on [0, 2*π*], it should be transformed to range over [*π, π*] to ensure that computations of the turning angle and its absolute values are computed correctly (see Eq A.2 in the Appendix A.2).

Movement from current locations 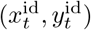 to a set of neighboring cells(*a, b*) is computed in terms of kernels *K*_*α*_, *α* = 1, …, *n*^stame^ belong to a set 𝒦, where each kernel *K*_*α*_ is defined as the rim of the sector of a circle centered on the origin, with rim dimensions 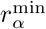 and 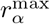 and sector angle 2*ψ*_*α*_ (Fig 2A) and includes an additional parameter 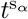 that influences the amount of time spent consecutively using kernel *K*_*α*_:

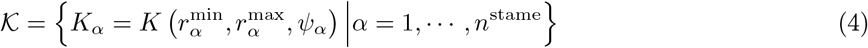

**Figure 2.**
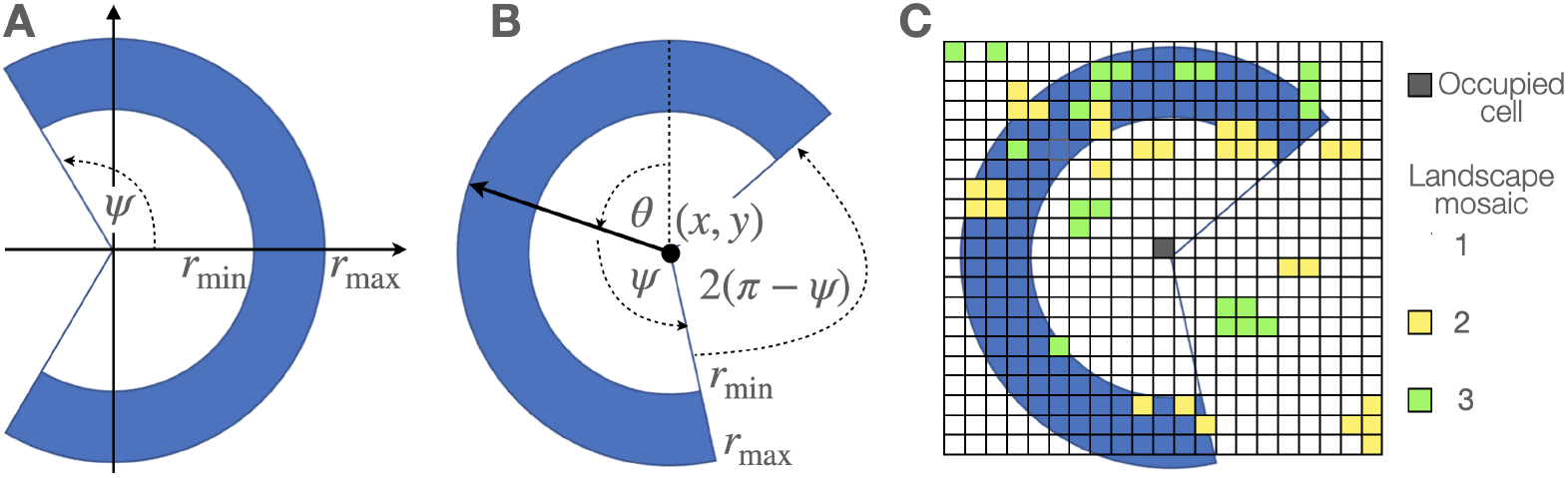
Kernels (**A**. area in blue) are specified in terms of the rims of sectors of circles with subtending angle 2*ψ* and inner and outer rim radii *r*^min^ and *r*^max^ respectively. Kernels become step-selection functions when located at a point (*x, y*), provided an angle of heading *θ* (**B**.), placed on a landscape (**C**.) with cells of different levels of attraction or repulsion (green, yellow, and white squares) and admissibility (all cells that overlap with the kernel), and associated with a step selection rule ℛ.

Of course, if 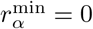, then the rim is actually a sector (slice) of a circle of radius *r*^max^.

The kernels *K*_*α*_ become step-selection functions 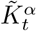 when anchored at time *t* at a point 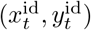 with heading angle *θ*_*t*_ and ‘time spent in current movement mode’ variable 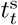 (Fig 2B). They are associated with a set of movement rules 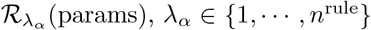 (params), *λ*_*α*_ ∈ {1, …, *n*^rule^}, where “params” are any parameters in the procedure that it is convenient to highlight. In particular, params includes a switching parameter 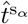 that controls the probability of switching out of the current kernel as a function of 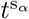 and neighborhood cell value parameter 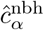: i.e.,

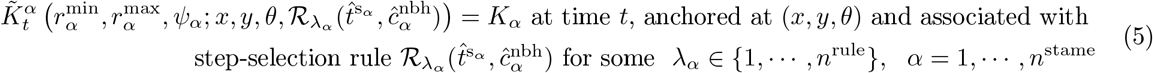

### 2.2 Landscape and individual dynamics

The initial set of landscape values 𝒜 (0) (Eq 1) can either be read in or generated using an algorithm that, for example, either assigns cell values at random or generates patches of high valued cells in a matrix of low-value cells and even barrier cells in more complex versions of the model than presented here. We have implemented an algorithm for constructing either of these two cases using three parameters and default starting values of *c*_*ab*,0_ = 0 or 1: a parameter value *p*^seed^ for laying down the first cells in patches at random, a parameter *p*^cont^ for building patches into neighborhoods, and a parameter *n*^cont^ for controlling the expected sizes of these patches. At the start of a simulation an individual animal has a state *h*_0_, which is a representation of, for example, stored energy.

#### Patchy landscape generation

To initialize the landscape either read in an initial landscape file 𝒜 (0) (Eq 1) or select one of the following two algorithms designated as 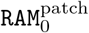 and 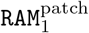, following procedures laid out in the Section 4 (`RAM` is an acronym for Runtime Alternative Module and will be described in more detail later).

#### Randomly located regular patches 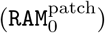

L.1 Default state: Set all cell values *c*_*ab*,0_ = 0 for *a* = 1, …, *n*^rows^, *b* = 1, …, *n*^cols^
L.2 Lay down an initial set of patch seeds by switching the value of cell(*a, b*) from *c*_*ab*,0_ = 0 to *c*_*ab*,0_ = 1 with probability *p*^seed^ for *a* = 1, …, *n*^rows^, *b* = 1, …, *n*^cols^.
L.3 For *a* = 1, …, *n*^rows^, *b* = 1, …, *n*^cols^ if *c*_*ab*,0_ = 1 then switch all cells that lie within a Moore neighborhood of radius *n*^cont^ to cell(*a, b*) to 1. This includes cells that may already have been switched to 1 because of their proximity to some other cell that has already been switched to 1.

#### Randomly located irregular patches 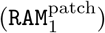

L.4 Default state: Set all cell values *c*_*ab*,0_ = 0 for *a* = 1, …, *n*^rows^, *b* = 1, …, *n*^cols^
L.5 First pass: Lay down an initial set of patch seeds by switching the value of cell(*a, b*) from *c*_*ab*,0_ = 0 to *c*_*ab*,0_ = 1 with probability *p*^seed^ for *a* = 1, …, *n*^rows^, *b* = 1, …, *n*^cols^.
L.6 Second pass: For *a* = 1, …, *n*^rows^, *b* = 1, …, *n*^cols^ if *c*_*ab*,0_ = 0, and the sum of the values of cell(*a, b*)’s 4 neighbors (von Neumann neighborhood) is Sum_4_ (*c*_*ab*_) then switch cell(*a, b*) from *c*_*ab*,0_ = 0 to *c*_*ab*,0_ = 1 with probability 1 − (1 − *p*^cont^)Sum_4_ (*c*_*ab*_)
L.7 Additional passes: Repeat step (c) *n*^cont^ times (with the second pass corresponding to *n*^cont^ = 1).

The the default for the initial value *h*_0_ of an individual is set 10. Other values can be entered as discussed in the simulation parameter setup below.

We note that if *p*^cont^ = 0, then cells in the array are randomly assigned a value 1 with probability *p*^seed^

#### Dynamic updating

The cell array values *c*_*ab,t*_ and individual value *h*_*t*_ for *t* = 0, …, *n*^time^ are updated to account for the possibility that the individual gathers or extracts resources from cells as it moves over the landscape. This extraction may only take place during the implementation of some movement modes but not others.

Here we account for changes in these values as follows. The individual’s value *h*_*t*_ changes over time as it acquires resources when occupying cells of value *c*_*ab,t*_ *>* 0. It also incurs a cost *κ*^sub^ per unit distance moved at each time step. Each time an individual occupies a cell(*a, b*) it removes some resources *f* ^remove^. If we assume that removal is a resource density independent process, with *f* ^remove^ = min{*κ*^add^, *c*_*ab,t*_} then the following updating rules for the value-state of individuals (*h*) and cells (*c*) and parameters *κ*^add^ ∈ [0, 1] and *κ*^sub^ apply:

In moving from 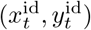 at ti e *t* to cell(*a, b*), we have the resource density independent process

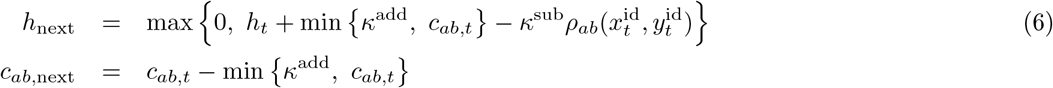

We can make this extraction dependent on the density of resource using the form of 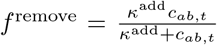 which is familiar to those who model consumer-resource interactions [53]. We can also allow for growth of the resource back to its carrying capacity of 1 at a a rate *κ*^grw^ when completely removed (e.g., for grasses this represents regrowth from an intact root-stock, etc). Finally we can also make the cost of travel size (energy value) dependent on a suitable scaling constant *κ*^scl^ ≥ 1 and multiplying 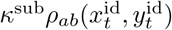 by, for example, 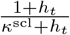. In this case, as 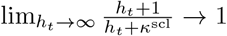 and at *h*_*t*_ = 0 we obtain the factor 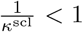. In this case, we obtain the equations:

In moving from 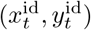 at time *t* to cell(*a, b*), we have the resource density-dependent process

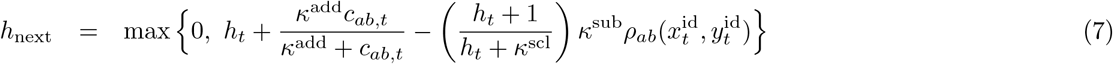

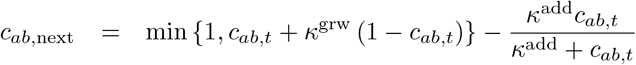 for a currently occupied patch cell(*a, b*)

*𝒸*_*ab*_, next = min {1, *𝒸*_*ab,t*_ + *κ*^*grw*^ (1 − *𝒸*_*ab,t*_) for all unoccupied patch cell(*a, b*)

We note that the simulation will stop either at *t* = *n*^time^ or at *t*^stop^ if *h*(*t*^stop^) = 0.

In addition to the state value *h* of the individual, we will also keep track of the time 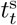 it has spent in its current movement mode. Thus, we will update 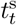 as follows

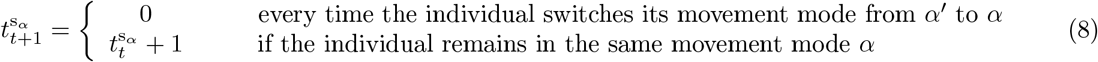

Thus at the start of the simulation, no matter the starting kernel, we initialize *t*^s^(0) = 0.

### 2.3 Movement kernels

In our Numerus ANIMOVER_1 RAMP, we limit the implementation to at most two movement modes and hence two StaME kernels *K*_*α*_, *α* = wp, bp. More specifically, we use *α* = wp to produce within-patch movement tracks and *α* = bp to produce between patch movement tracks. In addition, when *α* = bp we employ a vision kernel 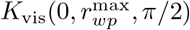 which allows an individual to see patches peripherally and in front of it to the maximum radius of its next *α* = bp movement-mode step.

#### Kernel parameters

In general we expect within patch steps to be smaller than between patch steps, and within patch turning to be larger than between patch turning, although we have the flexibility to set up contrary scenarios. For each of the two movement modes at the start of a simulation, values for the triplets 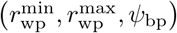 and 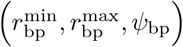 are set up. If our expectation is followed, then we 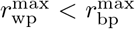 and *ψ*_bp_ < *ψ*_wp_ will hold. We may also expect 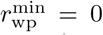 if the individual may choose to remain in the same cell for more than 1 time period and 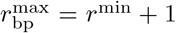 or 2 if within patch movements are considerably less than between patch movement where we suggest setting 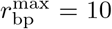. We again stress that the user has the flexibility to set and scale this values arbitrarily.

All that remains in implementing the algorithm depicted in Fig 3, is specification of the parameters needed to implement our step-selection functions (SSFs, [54, 55, 56]) 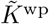 and 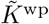 using the rules 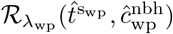 and 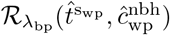 that are specified next (the last of these parameter arguments applies to 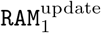, but not 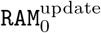). We note that these procedures are coded as the default `RAM`s for our Numerus ANIMOVER_1 RAMP. Other procedures, possibly using integrated step selection analysis (iSSA, [57]) or methods that include direction-biasing external points of attraction or repulsion [58], may be included by the user.

**Figure 3.**
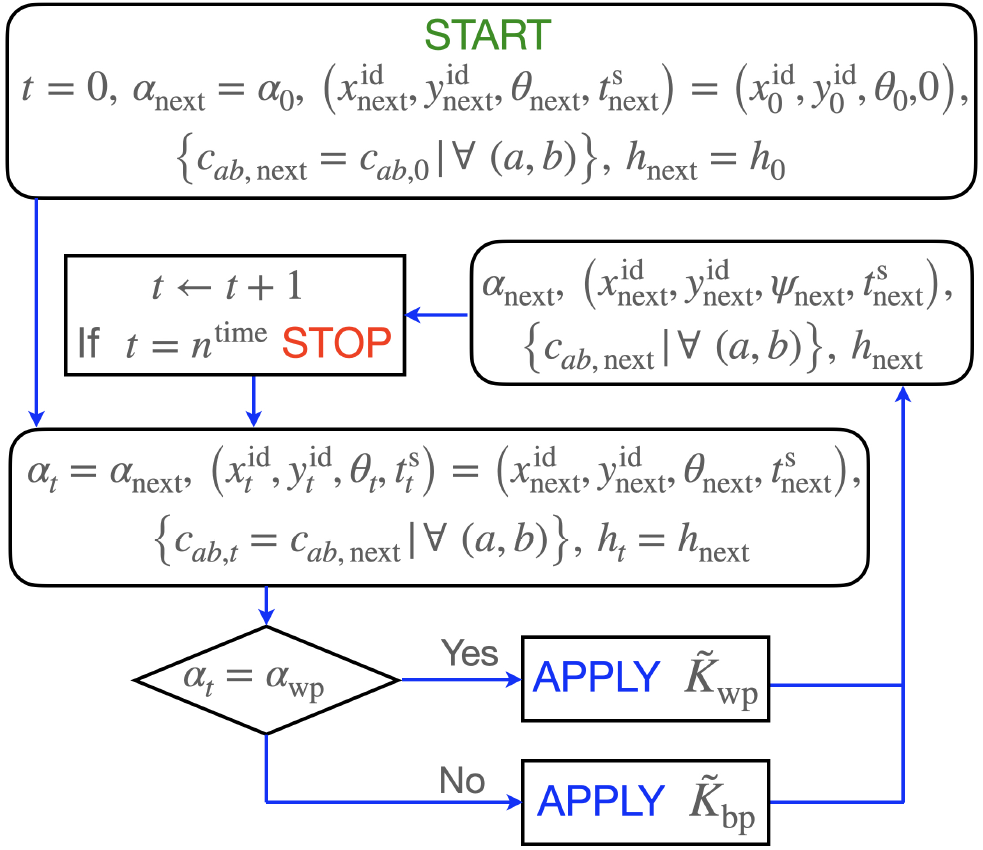
Algorithm used to simulate movement after the structural parameters *n*^time^ (index *t*), *n*^row^ (index *a*) and *n*^col^ (index *b*) been entered, along with the topology of the landscape, the initial landscape values {*c*_*ab*,0_|*∀*(*a, b*)}, the initial state value *h*_0_ of the individual, and the parameter triplets 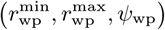 and 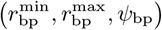 that define the two kernels *K*_wp_ and *K*_bp_ respectively. Implementations 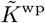 and 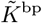 require the specification of step-selection procedures 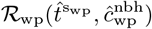 and 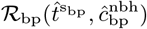 respectively. The latter employs an additional kernel *K*_vis_ that controls how the individual scans the landscape when it is moving between patches. The START, STOP, and APPLY commands, along with the flow arrows are show in different colors for additional clarity. The details of step selection procedures 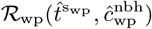 and 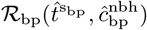 can be found in Appendix A.3.

#### Step-selection cells and probabilities

The step-selection rules 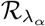 are procedures for updating the next location, angle of heading, and time spent in the current movement mode (*x*_next_, *y*_next_, *θ*_next_, 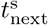), as well as the kernel *α*_next_ to be used next.

This is done in terms of the individuals current location, angle of heading and spent time 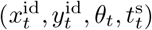, and its current movement mode, as driven by the kernel implementation 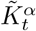. In our case we have two sets of rules, one that specifies within patch movement 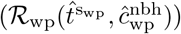 and one that specifies between patch movement 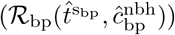. These rules include situations where the topology of the landscapes is a bounded rectangle and normal search fails to find a next location.

In computing our step-selection procedures we will make use of the set 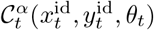 defined as follows:

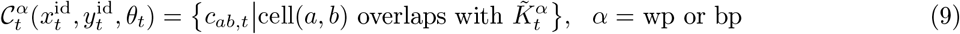

We will also make use of the following sets of probabilities

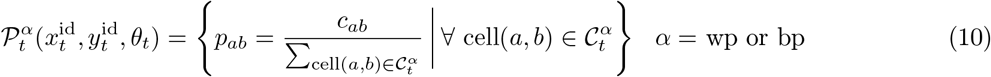

Finally, we will make use of the probabilities *p*_*α*_(*t*^s^) of continuing to use StaME *K*_*α*_ when having used this StaME for the past *t*^s^ time steps

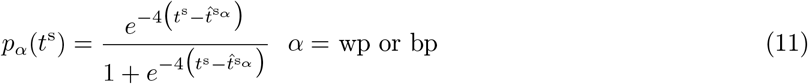

An outline of the implementation of step selection procedures 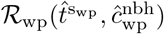 and 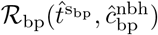 is provided in Fig 3 with details provided in Appendix A.3.

## 3 Movement Paths and StaME Extraction

A movement track, whether simulated or empirical, in the first instance has a representation as a relocation time series of *n*^time^ points—i.e., a walk

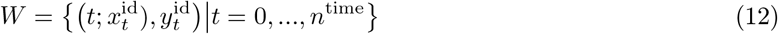

Such tracks can be generated using simulations models, as we discus in some depth in the next section. In the case of empirical data, though, data preprocessing and filtering [59] are needed to get rid of spurious or problematic points or fill in missing points. From walk Eq 12, both velocity and turning-angle time series can be generated, as outlined in Appendix A.2 and elsewhere [30]. Various other statistics (e.g., travel distance, net displacement) can be extracted as well and used as variables to define a set of basic statistical movement elements (StaMEs), following methods described next.

### 3.1 Creating StaMEs

The method described here to create StaMEs uses the velocity (*V*) and turning-angle (ΔΘ) time series derived from walk *W*, as described in Eq A.1-A.4 (Appendix A.2, Supp info). All points in our empirical time series that produced unrealistically large velocities were then removed. Although the putative maximum sustained flying speeds of barn owls have been observed in the range of 6-8 m/s (17.9 mph) [60, 61], to be conservative, we only removed points that represented unrealistically speeds. In our case, this amounted to a handful of points with velocities in excess of 75 m/s (168 mph); and we note that the average speed of the fastest 1 minute segments in our analysis below (Table 2) turns out to be a credible 3-4 m/s.

In the case of our simulated data, we normalized the entries of our cleaned *V* and ΔΘ time series by dividing each of the entries *v*_*t*_ in *V* by 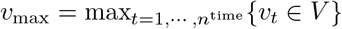 (= 10 in our simulations) and dividing each of the entries in ΔΘ by 2*π* to obtain variables on the ranges [0, 1] and [−1*/*2, 1*/*2] respectively. In the case of our empirical data, we only normalized the turning angles since the velocities had the physical units m/s and we wanted to reflect this in our results.

Next, we parsed the normalized velocity and turning-angle time series into *z* = 1, …, *n*^seg^ segments each of length *μ* (Fig 4) with regard to *t*: in segments with missing points (these points may be filled in using an appropriate interpolation method) we kept the same length of segment and just adjusted for the reduced number of points (the errors from such missing points are likely to be inconsequential when the number missing is a few percent or less). The total number of segments so obtained was 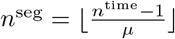, with some points left over when *n*^time^ − 1 was not exactly divisible by *μ*.

**Figure 4.**
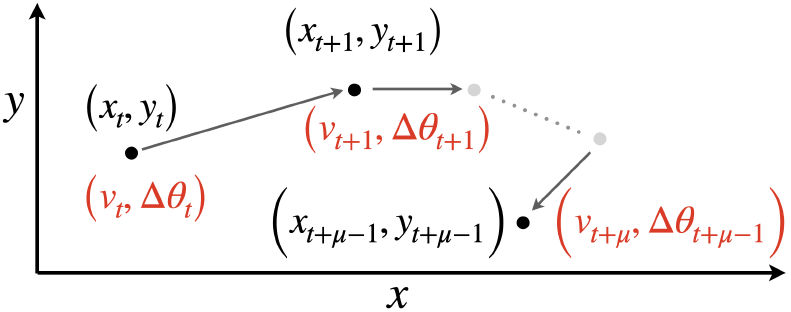
A relocation time series *W* (Eq 12) is plotted in the *xy*-plan with computed velocities *v*_*t*_ (Eq A.2, Supp info) and turning angles Δ*θ*_*t*_ in red (Eq A.4) for a segment of *μ* points at times *τ* = *t, t* + 1, *…, t* + *μ −* 1. These values, or others (e.g., 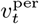 and 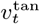; see Eq A.5 and [18]) can then be used to compute statistical elements Seg_*z*_ (Eq 13) from a series of consecutive elements containing *μ* relocation points as, for example, in Eq 13

We then calculated a set of statistics related to the *μ* normalized (in the simulated data only) velocities (equivalent to step lengths) and turning angles for each of the segments *z* in our time series data. Although various sets of statistics can be used (such as persistent and tangential velocities [18]), we settled upon mean velocities *V*_*z*_ and mean absolute turning angles |ΔΘ|_*z*_ for each segment and associated standard deviations 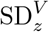 and 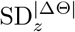. Also to pick up any possible circular motion type biases in movement, we computed a normalized net displacement (Δ^*ρ*^) statistic for each segment (i.e., the distance between the first and last points of each segment divided by quantity equal to the the number of points multiplied by the mean step-length). Specifically, for velocity and turning-angle means and standard deviations (SD) (normalized where appropriate), as well as net displacements, we defined a set of segments 𝒮 _*μ*_ (note below that *v*_max_ = 1 for the empirical data and *μ* is adjusted for segments with missing points)

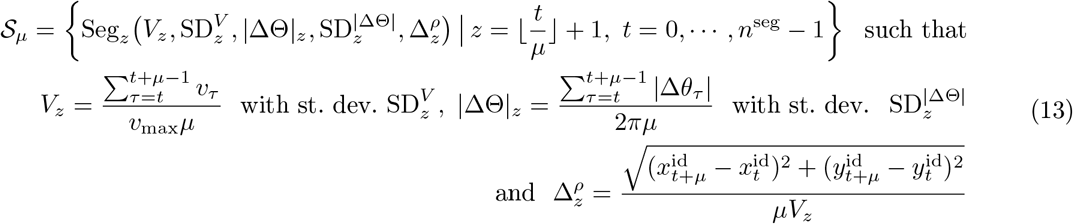

### 3.2 Mapping StaME Centroids to Kernels

We first applied our segmentation procedure Eq 13 to data obtained from simulating the movement of an individual using a single kernel 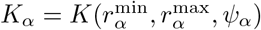 on an unstructured (static, homogeneous) landscape. If one generates a segmentation set 𝒮_*μ*_ (Eq 13) from these simulation data, then using a variety of kernels with different admissible 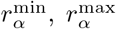 and *ψ*_*α*_ values, one can build up a discrete map represented by the function

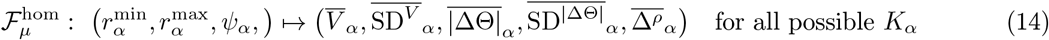

We note that we have not subscripted the quantities in the image of this mapping with the parameter *μ*. The reason is that the difference in values obtained for different values of *μ* are just variances associated with sampling and therefore are not consequential. However, we maintain the subscript *μ* on the mapping itself to remind ourselves that a value needs to be selected before this set can be generated. We also note, in the case of an unbiased walk, the statistic 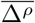 may be ignored, because the realized movement has no prevailing circular bias to its motion. Under these circumstances,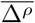 is perfectly correlated with the other statistics defining each segment because clockwise and counterclockwise movements are equally likely. Additionally, values obtained should be relatively insensitive to *μ* when *μ* and the length *n*^time^ of the track itself are sufficiently large to ensure that the law of large numbers is at play.

We then applied this segmentation procedure to data obtained from an ANIMOVER_1 two-kernel patch simulations (*K*_wp_ and *K*_bp_). Once the segmentation set 𝒮_*μ*_ had been generated (Eq 13), we carried out a hierarchical cluster analysis using Ward’s method (Appendix A.4) to obtain a set of *k* centroids represented by the set

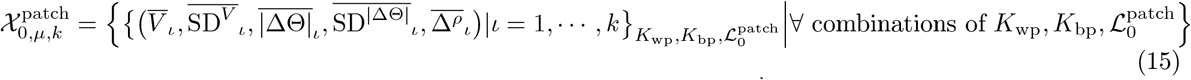

As in the case of quantifying selected points of the discrete map 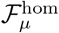, so can we quantify selected points of a discrete map 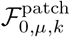 by computing *k* centroids in the set 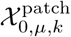 from simulations of ANIMOVER_1 using different combinations of *K*_wp_ and *K*_bp_ kernels (i.e., step-selection procedures ℛ_wp_ and ℛ_bp_) with individuals moving over generated landscape 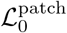. This mapping, in terms of the parameters used to generate it, can be expressed by:

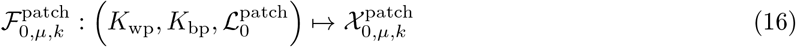

In our computations, we selected our cluster number *k* to be 8 rather than 2, (see Appendix A.4 for details) even though we only had two movement modes. The reason for this is that some segments in our segmentation process will be mixtures of the two modes rather than homogeneous strings of points generated by either one or other of the two modes (see Fig C.1, Supp info Appendix C) while we expected only 2 of the 8 clusters to contain relatively homogeneous movement mode segments. Of course, the extent to which mixed versus pure movement mode segments arises depends both on the length of segments and the frequency at which the two movement modes switch between each another. The shorter the segments, or the less frequent the switching, the more likely any segment represent a series of locations generated by a single movement mode.

We can use our simulation model to numerically construct a mapping 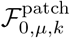 of a set 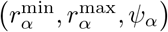 of kernel arguments onto a set of selected cluster centroid statistics 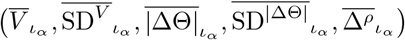 as outlined in Appendix B. By way of illustration, we generate one point of the map 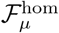 in this paper. The range of values that may be useful to generate in a multi-point construction of this map will depend on the kind of empirical to which this mapping is fitted. Such an exercise is, thus, best left to a detailed study that explores the structure of given set of empirical movement data, such as an extended set of the owl data discussed below—a set containing data collected from at least several tens of individuals (e.g., as in [42, 51]).

## 4 Numerus RAMPs

In the mid-2000’s Grimm et al. [62] proposed an Overview, Design concepts and Details (ODD) protocol for the presentation of agent and individual-based models (ABMs and IBMs). A recent update of this protocol Grimm et al. [63] identifies 7 elements (3 overview, 1 design, and 3 description) needed to provide a coherent presentation of the study that conforms to the central tenant of science that “materials and methods must be specified in sufficient detail to allow replication of results Grimm et al. [63].” In Appendix D, we present an ODD protocol for the development of ANIMOVER_1, following Grimm et al. [63]’s ODD element numbering scheme.

We stress, though, that our study is about more than just building a simulation model to be used to simulate a known process using a currently available coding platform such as `R` or `Python`. In Sections 2-3 we have presented a modeling framework that contains novel features regarding how concepts from step-selection function theory [54], when combined with a set of flexible rules that allow one to compute the probability of switching movement kernels (each of which represents a particular behavioral mode) in terms of local environmental factors, as well as internal clock variables, provides for the construction of a more versatile simulation model than, to the best of our knowledge, currently exists.

In the rest of this section, we present details that emphasize the utility, flexibility, and ease of use of our application platform ANIMOVER_1, based on its implementation of Numerus’ runtime alterable model platform technology. Although, ANIMOVER_1 itself is restricted to switching between two kernels that respectively implement within-patch (wp) and a between-patch (bp) movement modes, from the more general formulation of Sections 2 and 3, it is clear that future versions of ANIMOVER_# can be developed that allow switching among many more modes of movement. It will also become clear in our presentation that although ANIMOVER_1 is not a general programming environment and does not require any coding experience to implement it, it does provide the user with considerable flexibility to implement sections of code pertaining to landscape patch structure initialization and to equations used to update the agent’s and landscape cells’ current states (i.e., the user may substitute the code in the runtime-alterable modules—RAMs—described below that implement Eqs 6 or 7 for their own customized versions of these computations).

Finally, we emphasize once more that ANIMOVER_1 is downloadable for free along with the Numerus Studio Platform needed to play it (links provided in Appendix C).

### 4.1 RAMP Construction

The java-based Numerus Model Builder Designer (**NMB Designer**) Platform was used to code the model and then generate a **RAMP** (runtime alterable model platform) our **Numerus ANIMOVER_1** Application (with future versions 2, 3, etc. planned to include multidimensional landscapes, additional movement modes, and interacting individuals). The application takes the form of a portable file that, as described elsewhere [64, 37], and is played on the free downloadable NMB Studio application.

The flexibility of NMB RAMPs, beyond manipulating parameter values using its sliders (Fig 5E and F) and windows (Fig 5E and F), is facilitated by its runtime alterable modules (`RAM`s, Fig 5G, Fig 6). These `RAM`s provide the user with the ability to chose alternative formulations of component parts of the model just priori to rerunning a current simulation or recode those component parts with alternate expressions. Thus, for example, in updating the value equations *h*_*t*_ and *c*_*ab,t*_, the RAMP uses the default 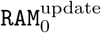, which codes Eq 6, as a default procedure, or the user can select 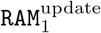, which codes Eq 7, as an alternate procedure (Fig. 6). In addition, the user can create a second alternative `RAM` by opening a new `RAM` window and inserting and saving code for customized equations, although parameters beyond the three already available as sliders at the console will have to be given fixed values (i.e., no new sliders can be created or added outside of upgrading the RAMP using the Numerus Designer Platform).

**Figure 5.**
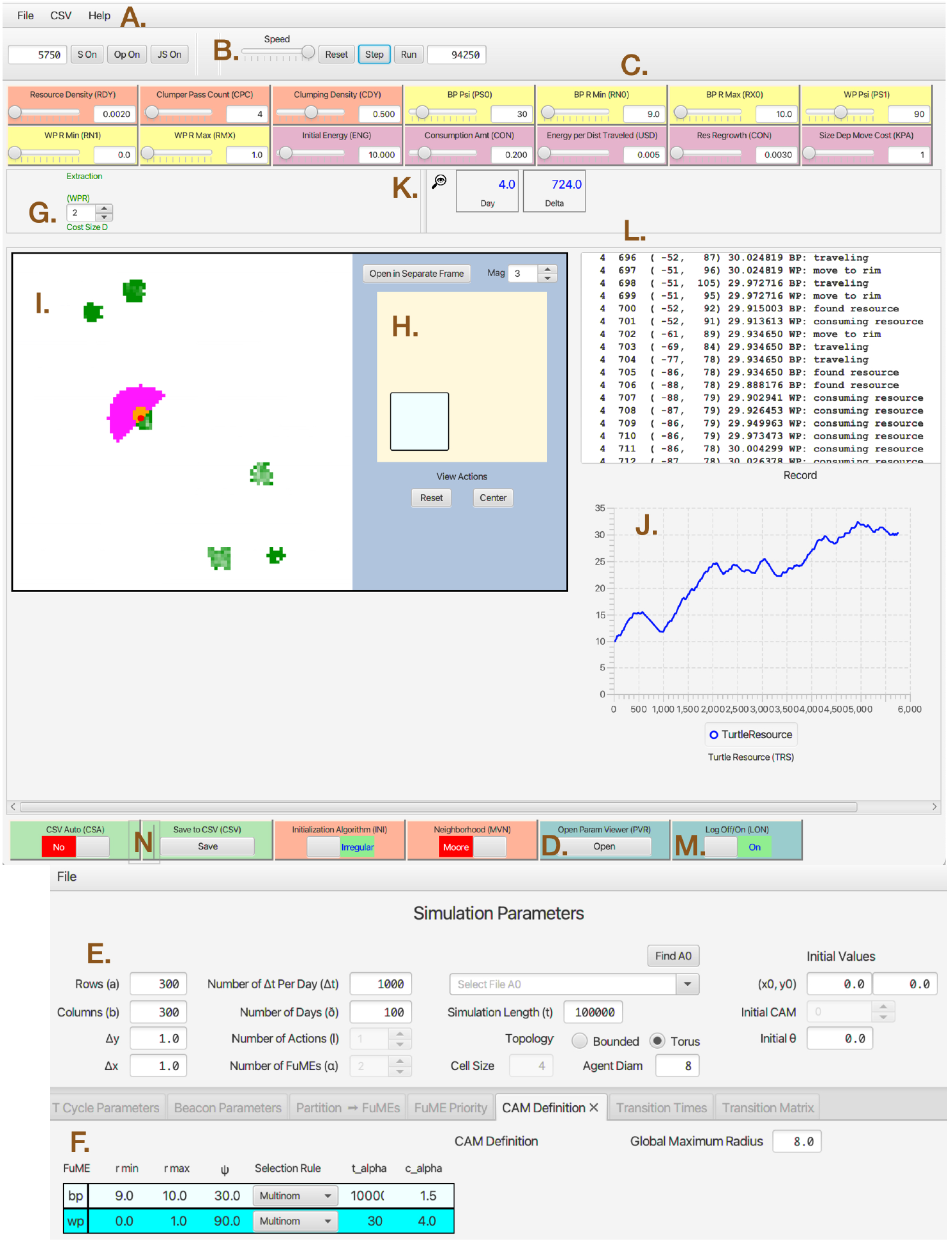
ANIMOVER_1 console as viewed in the Numerus Studio application. Labels A.-N. in brown are added for purposes of exposition. **A:** Pull down menus to load files. **B:** Run time controls allow for a reset in the middle or end of the run, step by step view of the changes (panels I and J), or automated run for full simulation time using parameters in panel E. **C:** Color coded sliders used to change parameter values at the start of or during a run (orange: landscape initialization; yellow: kernel parameters; lilac: consumer resource interactions; green: saving output; turquoise: parameter and monitoring windows. **D:** Button to open the parameter windows E & F. **E:** A form where various parameters that control the size and scaling of the run are set. **F:** A set of forms to enter between (bp) and within (wp) patch kernel parameters (angles in degrees). **G:** The wheel is used to select the `RAM`^update^ to be used. **H:** A viewer of the whole landscape with inset that can be observed as magnified in viewer I. **I:** A mouse-manipulable viewer of the landscape (panel H) that can be moved (using right click and move) and zoomed. Shown in I. are the current location (red dot), five patches (green pixels, lighter one previously exploited), and the current kernel (orange and lilac pixels). **J:** A plot of the value *h*_*t*_ of the agent over time. **K:** Record of current day and within day iteration with additional **L:** activity log and **M:** log switch. **N:** Switches and buttons to save end-of-run output.

**Figure 6.**
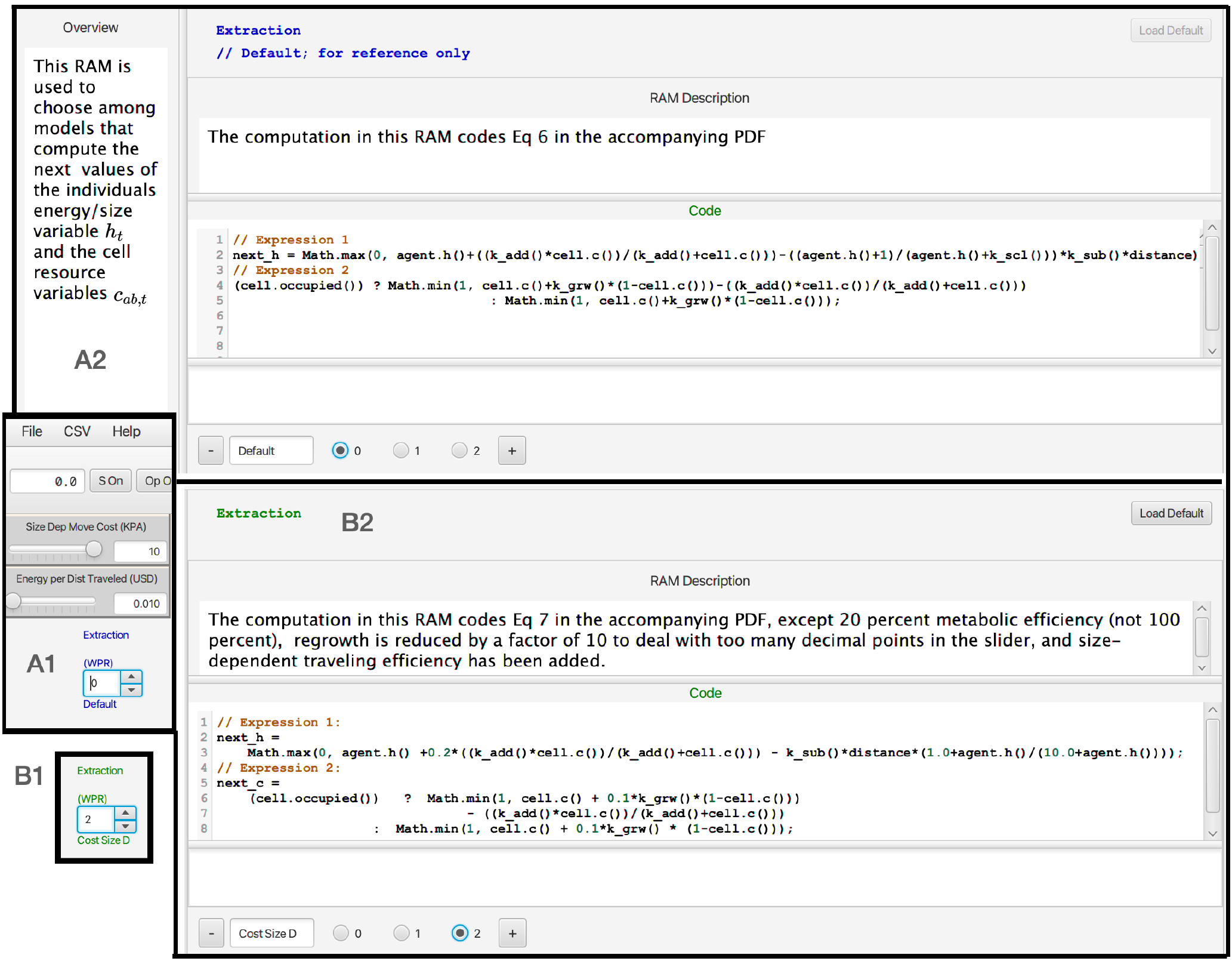
**A1:** The location on the ANIMOVER_1 console (**G**. in Fig 5) where the particular `RAM` to be implement in the current run is selected using the numbered roller. **A2:** The windows of the Default `RAM` showing the code that implements Eq 6. **B1:** This shows the roller selected to position 2 to implement the Extraction `RAM` expressed in Eq 7. Position 1 refers to a form of the equations not discussed in the text. **B2:** The windows of the Extraction `RAM` showing the code that implements Eq 7. The “+” sign at the bottoms of windows A2 and B2 allow the user to open a window for the user to supply their own customized code for updating the state of the individual *h*_*t*_ and cells array values *c*_*ab,t*_.

### 4.2 RAMs

Two or more versions of the `RAMs` listed below exist: first a default version (`RAM`_0_), a selectable alternative version (`RAM`_1_), and perhaps additionally higher numbered versions or extemporaneously created versions, versions that can be added just prior to initiating a run of the model, and saved for future use. Thus, for example, Eq 6 for the value dynamics of individuals and cells are coded up as default 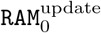, while the more elaborate density-dependent version of these equations (Eq 7) are coded up as 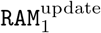.

ANIMOVER_1 includes the following `RAMs`:

Initial landscape `RAM`s

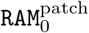 (default) This `RAM` lays down a patchy landscape using rules L.1-L.3 to create irregularly-shaped (stochastic) patches stochastically placed at a specified density. It requires parameter values *p*^seed^ to define the patch density, and *n*^cont^ and *p*^cont^ to generate the patch size and level of irregularity. Patches will be larger and more squarish when Moore is selected over von Neumann neighborhood.

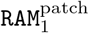 (alt 1) This `RAM` lays down a patchy landscape using rules L.4-L.7 to create regularly shaped patches (cubes using Moore neighborhood, diamonds using von Neumann neighborhood) stochastically placed at a specified density. It requires parameter values *p*^seed^ to define the patch density, and *n*^cont^ to generate patch size.

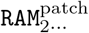 Available to the user for coding customized methods for creating initial landscape structures.

#### Resource Extraction RAMs

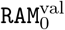 (default): This `RAM` updates the values *h*_*t*_ and *c*_*ab,t*_ using Eg 6 and requires parameter values *κ*^add^ and *κ*^sub^.

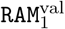 (first alternative): This `RAM` updates the values *h*_*t*_ and *c*_*ab,t*_ using Eg 7 and requires parameter values *κ*^add^, *κ*^sub^ and *κ*^grw^.

### 4.3 Parameter setup

The following set of parameters are needed to simulate the model. We also include some information on how our application console may be used to input some of the parameter values. Once this is done for those parameters that are either read in as a `FILE` (Fig 5A), entered prior to the simulation using a `FORM` (Fig 5E and F), specified using a `PULLDOWN` menu, or ignored—in which case `DEFAULT` values will be used, the Numerus ANIMOVER_1 will create a parameter values data file that can be download, edited and re-uploaded as needed. Other parameters will be entered and flexibly changed using a `SLIDER` (Fig 5C). Some of the these sliders will reflect values that are entered using a `FORM` while others will not be reflected in the form, but saved internally when the current simulation job is saved by the Numerus Studio Application Platform.

#### P.1 Size and scope parameters (FORMS)

These are: *n*^row^ (index *a*), *n*^col^ (index *b*), *n*^time^ (index *t*) and `topology` = `torus` or `plane` (`PULLDOWN`). We note that we have fixed *n*^stame^ = 2 (index *α*), Entry of these numbers and topology type respectively specify the dimensions of the cellular array, the length of the simulation, and whether the simulation takes place on a torus or a bounded plane.

#### P.2 Scaling parameters Δ*x*, Δ*y* (FORMS or DEFAULT)

Entry of these two Cartesian scaling values are used to assign the *x*-*y* coordinate values 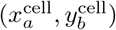 to each of the cells (*a* = 1, …, *n*^row^, *b* = 1, …, *n*^col^) in the simulation space 𝒜. The default values for this are Δ*x* = Δ*y* = 1.

#### P.3 Cellular array 𝒜 (0) (FILE)

or generating parameters *p*^seed^, *p*^cont^ and *n*^cont^ (`SLIDERS`). This is an appropriately configured data file (e.g., csv text) that will up uploaded to the application to provide the initial state of all the cells in the simulation space or the initial landscape will be generated using the specified parameters algorithm L.2.

#### P.4 Kernel definition parameters *K*_*α*_, *α* = wp and bp (FORM and SLIDERS)

Enter the three arguments for each of two kernels: i.e., 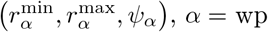 and bp, respectively.

#### P.5 Kernel implementation parameters for 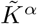 (SLIDER)

Enter the arguments 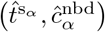 for the step-selection procedures ℛ_*α*_, *α* = wp and bp.

#### P.6 Initial values (various)

The initial time is automatically taken to be *t* = 0. The initial location and angle is computed from the selection of values (*a, b*) entered (`FORM`) or ignored (`DEFAULT`) and an initial value for the heading direction *θ*_0_ is entered (`FORM`) or ignore `DEFAULT`). The default starting cell is *a* = ⌊*n*^row^*/*2⌋, *b* = ⌊*n*^col^*/*2⌋, and default angle of heading is *θ*_0_ = 0. The actual starting location is thus 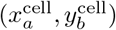. The initial value *h*_0_ for the individual must be entered (`Form`) or will be 10 (`DEFAULT`).

#### P.7 Initial kernel wp or bp (PULLDOWN)

This will set the initial condition *α*_0_

#### P.8 Update parameters *κ*^add^, *κ*^sub^, *κ*^grw^ and *κ*^scl^ (SLIDERS)

The first two of these parameters are used in Eq 6 or all four in Eq 7 for updating *h*_*t*_ and *c*_*ab,t*_.

#### P.9 Step selection rule parameters 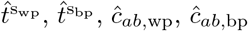 (SLIDERS)

The first two will be integer value sliders between [1, 1000], while the second two will be values to 1 dp between [0,5].

### 4.4 Simulation setups and units

If landscape pixels are Δ*x* = Δ*y* = 10 m and *n*^row^ = *n*^col^ = 500 then our 250,000 pixel landscape is 5^2^ = 25 km_2_. If the units of *t* are 1 m intervals, then in a day an individual will move up to 1440 = 60 × 24 times. If an individual walks at a persistent speed of 5 km per hour, which is 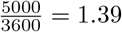 m/s, then an individual can cross either the length or breadth of our landscape in 1 hour, and a single pixel in 14 secs. In this case, the a maximum radius of *r*^max^ = 4 would suffice, although a movement kernel covering short sprints would have a maximum radius of 4 or 6 times this value. For theoretical studies, boundary effects can be avoided during simulations by setting `topology`=`torus` (which makes the left and right columns of cells in the landscape array neighboring columns, and the top and bottom rows of cells neighboring rows).

### 4.5 Model output

During the course of a simulation, one can visual observed the run as it progresses in window I (Fig 5) of the ANIMOVER_1 console. One can also observe the sequence of movements for the current day stage of the simulation in window L if switch M is in the “On” position (Fig 5). At the end of the run a CSV file is automatically saved if switch N is in the “Yes” position or if the “Save to CSV” button N is pressed after the run is complete. The saved CSV file is headed by a list of all the parameter settings for the run. It also has the following columns of data consecutive generated at step of the model simulation: day, within-day step, *x*-location, *y*-location, distance moved angle of head (degrees), agent-state (resources), movement mode (bp or wp) (Fig C.1, Supp info). An example of this output can be found in supplementary online file `Two_Kernel_Movement.csv` with a graphically depicted subset explained in Appendix C.1 (Output Data). In addition, links to downloading ANIMOVER_1 and to a RAMP Users Guide can be found in Appendix C.2.

## 5 Illustrative Examples

### 5.1 One point in the mapping of 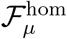

By way of illustration, we used ANIMOVER_1 to construct 1 point in the mapping 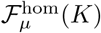 (Eq 14). Specifically, we used the kernel *K*(0, 6, *π/*2) to simulate the movement of an individual for 1500 steps over a homogeneous landscape. Using a *μ* = 15 points segmentation interval we generated the set 𝒮 _15_ (Eq 13) and then calculated the means of the statistical variables of the interest to obtain 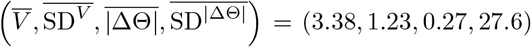. Thus we identified one point in the mapping 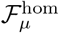 (Eq 14): specifically,

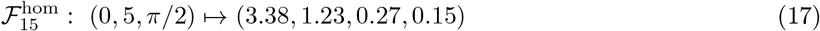

where we reiterate that we did not compute the displacement variable because it is non-informative due to the lack of a circular bias to the motion. Other points in the mapping can of course be constructed, as outlined in M.1-4 above. The range of parameters use to compute the structure of 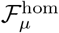 depends on the range of the image space 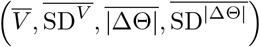 that needs to be covered. Interpolation can also be used, where desired to estimate points that are contained within the nodes of the lattice structure used to compute the mapping at discrete points in the range space (as discussed in Appendix B).

### 5.2 A two-movement mode simulation

We carried out a two-movement mode simulation on a patchy landscape that we manually stopped after 69,525 model steps. The parameters that we used for this simulation were:

#### Two movement mode simulation

For this simulation, we used the following parameters:

*Size and scaling. n*^row^ = 300, *n*^col^ = 300, *n*^time^ = 100, 000 (as an upper limit) and `topology` = `torus`, Δ*x* = Δ*y* = 1

*Landscape generator*. Initialization algorithm = Irregular, Neighborhood = Moore, *p*^seed^ = 0.1 (resource density), *p*^cont^ = 0.7 (clumping density) and *n*^cont^ (clumper pass count)

*Kernel parameters*. Within patch: 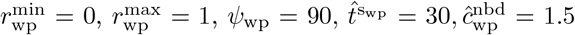; Between patch 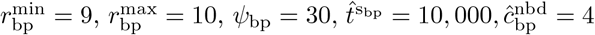

*Update parameters. κ*^add^ = 0.4, *κ*^sub^ = 0, *κ*^grw^ = 0.03 and *κ*^scl^ = 1.0

Under pure between-patch (bp) movement on a homogeneous landscape, given that 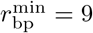 and 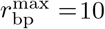, we should expect 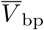 after normalizing by *V*_bp_ ≈ 10 to be in the range [0.9, 1]. Also given that the turning should, on average (since *ψ*_wp_ = 30 degrees) be around (0.5 × 30)*/*180 ≈ 0.083. We carried out such a simulation over 100,000 steps and obtained 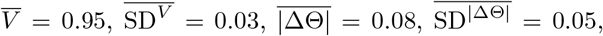, and 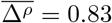. These values are almost identical to the values associated with the 10-step segmentation of the data that has the largest 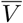 value or fastest speed (i.e., cluster C = 1), as reported in Table 1. Thus the segments in this cluster from our two movement mode simulation are very close to what we obtain when we simulate pure one-mode between-patch movement. The cluster with the largest 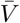 in our 30-step segmentation (cluster C = 8) does not fit nearly as well, presumably because many fewer pure bp movement segments arise in this case compared with the 10-step segmentation case.

**Table 1:** The centroid arguments (Eq 15) obtained from a hierarchical cluster analysis (with *k* = 8 clusters labeled C = 1, *…*, 8, see legends in Fig 7) analysis of 10- and 30-step segmentation of our two movement mode simulation output. The column “Prop. wp” lists the mean and standard deviation of the proportion of with-in to between patch kernels (i.e. 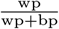; see Fig C.1, Supp info Appendix C) averaged across all segments in each cluster. The results are listed in descending size of average speed 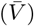 for the centroid of each cluster. Depictions of 130 randomly selected segments from each cluster for the 10-step segmentation case can be seen in Figs A.2-A.9 (Supp info Appendix A).

| C | # Segs. (%) | $\bar{V} \pm SD^{\dagger}$ | $\vert \Delta\Theta\vert \pm SD^{\vert \Delta\Theta\vert \dagger}$ | $\bar{\Delta\rho} \pm SD^{\Delta\rho \ddagger}$ | Prop. wp |
| --- | --- | --- | --- | --- | --- |
| **10-step segs\*** |  |  |  |  |  |
| 1 | 811 (11.7) | 0.95±0.04 | 0.08±0.05 | 0.86±0.14 | 0.01±0.04 |
| 7 | 694 (10.0) | 0.84±0.21 | 0.28±0.25 | 0.14±0.11 | 0.51±0.10 |
| 2 | 776 (11.2) | 0.75±0.33 | 0.18±0.18 | 0.79±0.13 | 0.27±0.15 |
| 8 | 546 (7.9) | 0.38±0.40 | 0.27±0.23 | 0.81±0.08 | 0.66±0.16 |
| 6 | 838 (12.1) | 0.38±0.38 | 0.33±0.25 | 0.11±0.06 | 0.75±0.10 |
| 5 | 1057 (15.2) | 0.37±0.36 | 0.31±0.23 | 0.46±0.14 | 0.71±0.17 |
| 3 | 987 (14.2) | 0.12±0.10 | 0.35±0.23 | 0.18±0.07 | 0.90±0.10 |
| 4 | 1243 (17.9) | 0.10±0.07 | 0.34±0.23 | 0.44±0.12 | 0.93±0.08 |
| **30-step segs\*** |  |  |  |  |  |
| 8 | 138 (6.0) | 0.91±0.15 | 0.14±0.14 | 0.27±0.14 | 0.18±0.17 |
| 6 | 51 (2.2) | 0.86±0.20 | 0.12±0.12 | 0.82±0.09 | 0.12±0.10 |
| 7 | 306 (13.2) | 0.71±0.36 | 0.25±0.24 | 0.17±0.12 | 0.48±0.16 |
| 5 | 416 (18.0) | 0.53±0.41 | 0.25±0.23 | 0.49±0.09 | 0.52±0.13 |
| 1 | 463 (20.0) | 0.40±0.39 | 0.27±0.23 | 0.70±0.10 | 0.63±0.15 |
| 3 | 361 (15.6) | 0.39±0.39 | 0.31±0.25 | 0.14±0.10 | 0.74±0.07 |
| 2 | 246 (10.6) | 0.20±0.26 | 0.33±0.23 | 0.29±0.11 | 0.85±0.06 |
| 4 | 336 (14.5) | 0.15±0.18 | 0.35±0.23 | 0.08±0.05 | 0.89±0.06 |

\*All numbers are dimensionless due to normalizations (see text for details)
<sup>†</sup>This is the average of the standard deviations reported for each segment in the cluster
<sup>‡</sup>This is the standard deviation of the normalized net displacement across all segments in the cluster

This last conclusion is reinforced by looking at the mean proportion of wp kernel steps associated with the segments of the different clusters reported in right-most column of Table 1. Thus only 1% of the 811 cluster 1 (largest 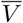) segments of length 10 steps were generated by a wp kernel, while the mean proportion of wp kernel steps associated with the two largest 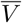 30-step segments were clusters 8 and 6 with reported proportions 0.12 and 0.18 in clusters 6 and 8 respectively. At the other end of the velocity spectrum, the two smallest velocity 10-step segment clusters 3 and 4 reported a mean of 0.90 and 0.93 wp segments, while in the 30-step segment case the two smallest velocity clusters 2 and 4 reported a mean of 0.85 and 0.89 wp segments respectively. This result reinforces our earlier comment and obvious result (see Fig C.1, Supp info Appendix C) that segments with fewer steps provide a greater proportion of pure step-type StaMEs than those based on more steps.

The remaining seven 10-step segment clusters and six 30-step segment clusters contained segments that included mixtures of wp and bp, ranging from averages of a quarter to three quarters of each type in the segments of each cluster. This calls into question what the optimal number of base segment clusters and, hence number of StaME types should be. For example, to what extent should we merge clusters reporting similar segment statistics? The results reported in Table 1 suggest that for the 10-step segmentation perhaps clusters 3 and 4 could be combined, although they are well-separated by their mean relative net-displacement value (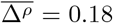 versus 0.44 respectively). This is true of many clusters in the 30-step case. For example, ignoring the value of 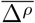 suggests that clusters 6+8, 5+1+3, and 2+4 might be combined to yield four distinct clusters, but in each case the variable 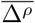 provides some cause for separation. In general, however, the optimal number of StaMEs will be data dependent. Additionally, finding the optimal number requires that we develop suitable measures so that such questions can be answered with some rigor. Such measures are currently being developed in the context of an information theory formulation of CAM construction using StaMEs as a set of building block segments [34].

A random selection of 130 segments from each of the 8 clusters in the case of the 10-step segmentation are plotted in Figs A.2-A.9 (Supp info, Appendix A, where we note that the actual size of segments across panels is not comparable because the axes in each panel have been automatically set by our plotting routines. From these illustrations it is clear that the fastest cluster (C = 1; Fig A.2, Supp info Appendix A) consist primarily of unidirectional lines, as is characteristic of between-patch movement, with some small deviations and occasional changes in direction. This is in contrast with the smallest cluster (C = 4; Fig A.5) that has a number of segments switching directions by *π/*2 every couple of steps, as is characteristic of in-patch foraging. The difference between segments C = 6 (Fig A.7) and C = 8 (Fig A.9), which have virtually the same average speeds (Table 1) and similar average turning angles look quite different: the start and end points are relatively close with spiky profiles (C = 6) or far apart with much more open profiles (C = 8).

### 5.3 Empirical data example

We analyzed relocation data obtained from two barn owls (*Tyto alba*), using an ATLAS reverse GPS technology system that was set up in the Harod valley in northeast Israel [51]. The relocation data for both individuals, an adult female and a juvenile male (GG41259 and GG41269 in the original data set and individuals with IDs 29 and 31 in [42]), were collected a frequency of 0.25 Hz (i.e., one point every 4 seconds) during a several week period in the late summer of 2021. We used a 15-step segmentation, which corresponds to segmenting the movement track into one minute sequences. We then performed a hierarchical cluster analysis with 8 clusters, described in Appendix A.4, and obtained the results illustrated in Fig 8. The centroid statistics obtained for each cluster are provided in Table 2. Unlike the simulated results were we normalized the velocities to lie between 0 and 1, the velocities in Table 2 have the units of meters per second. Also, recall that the absolute values of average turning angles 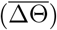 are proportions of *π* while the net displacements 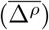 are the distances between the start and end points of each segment as a proportion of the sum of the lengths of the 15 consecutive steps that make up each segment.

**Figure 7.**
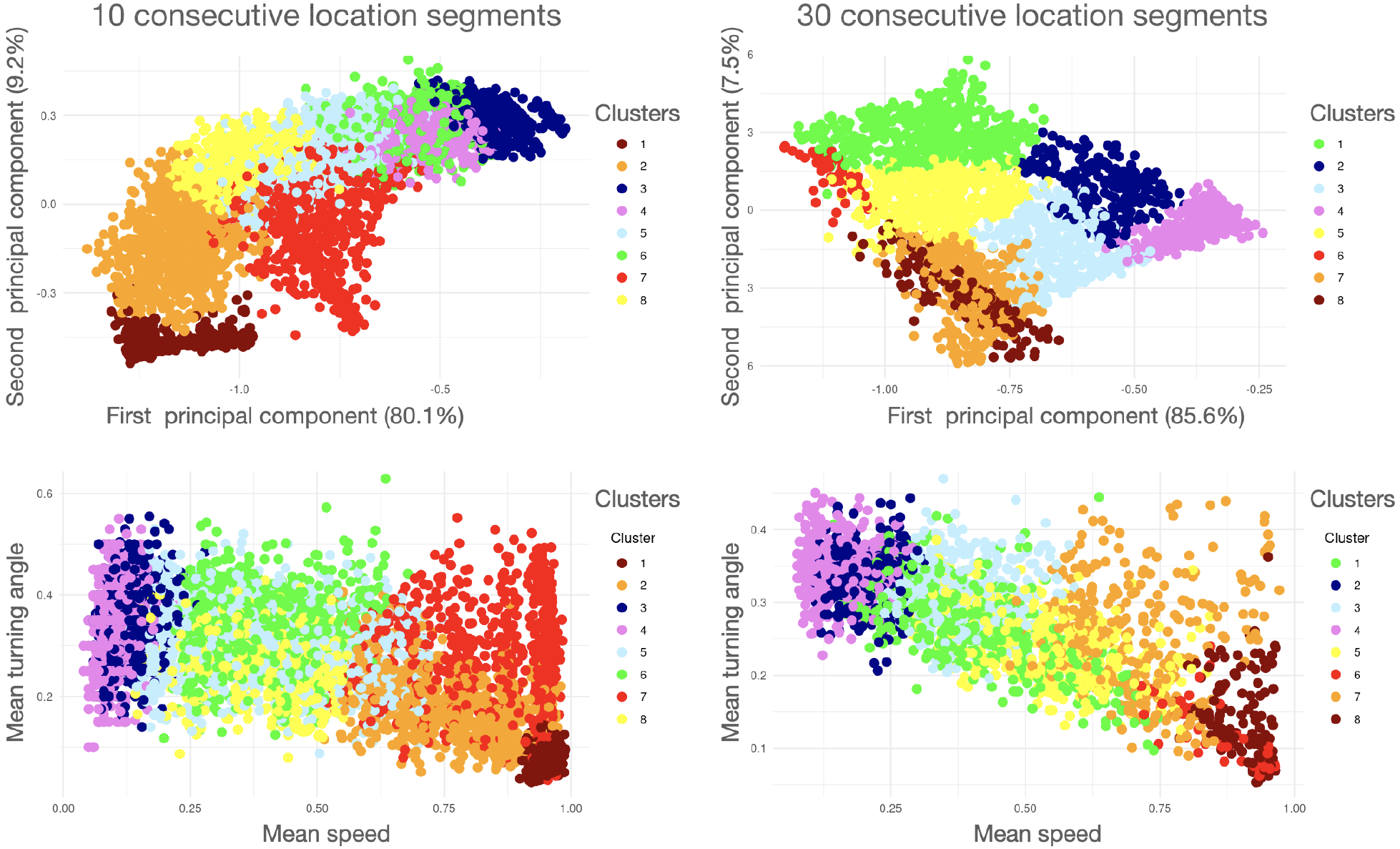
Results of a hierarchical cluster analysis (with number of clusters *k* = 8) performed on both 10-point (upper and lower left panels, *ν* = 10; see Fig 4) and 30-point (upper and lower right panels, *ν* = 30) segmentation of the simulation data generated with parameters specified for our two movement mode simulation (also see Fig C.1, Supp info Appendix C). The two top panels are plots of the clusters in PC1/PC2 space and the two lower panels are plots in Mean-speed/Mean-turning-angle space. A color spectrum is used to depict the smallest (blue end) to largest (red end) of segments within clusters. The centroid arguments of each cluster (Eq 15) are listed in Table 1. The colors have been selected to reflect a spectral scale of the largest to smallest mean speed for each cluster’s centroid, though the cluster numbers (Column C in Table 1) are set by the clustering algorithm. Depictions of 130 randomly selected segments from each cluster for the 10-step segmentation case can be seen in Figs A.2-A.9 (Appendix A, Supp info).

**Figure 8.**
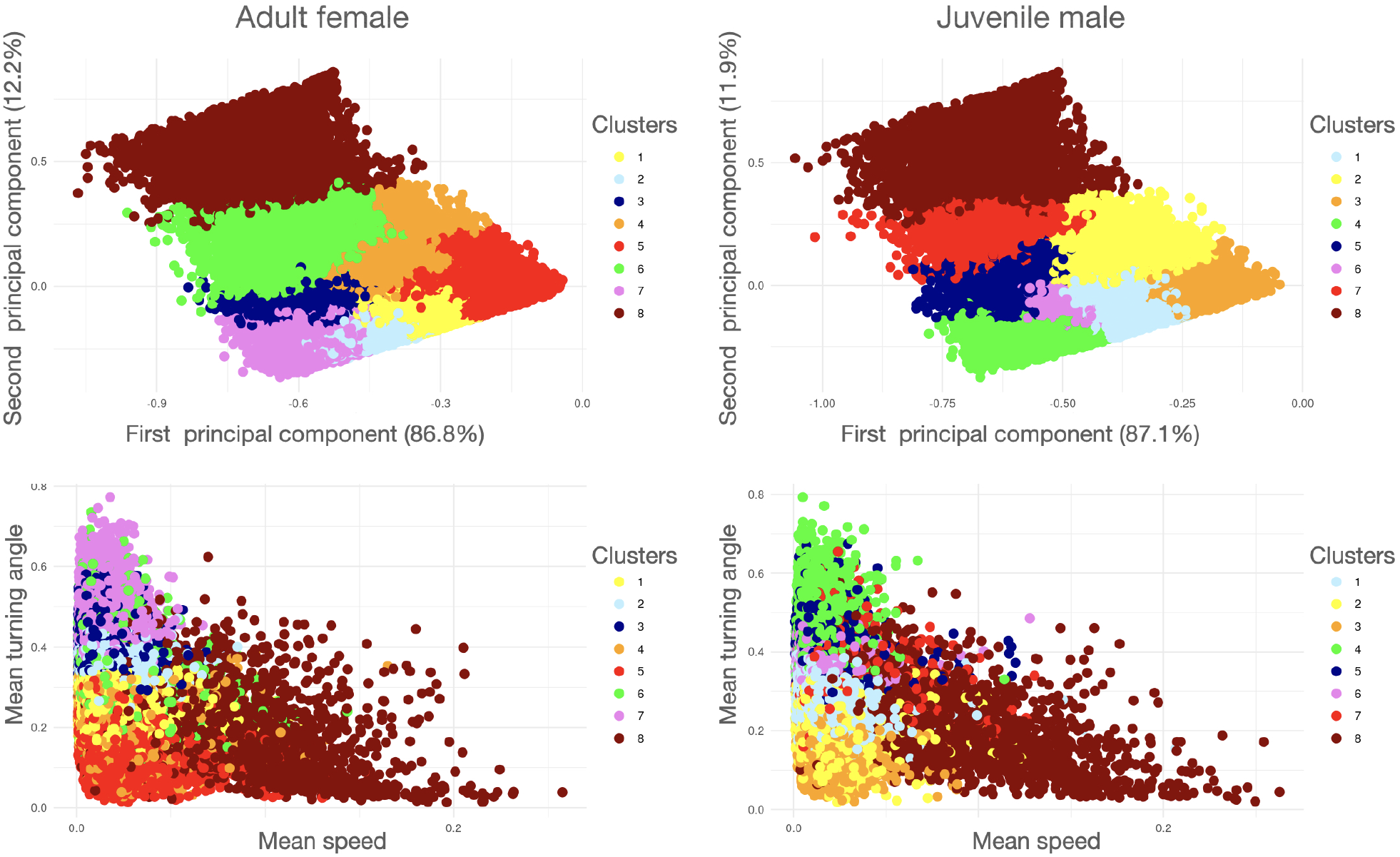
Results of cluster analysis (*k* = 8) performed on segmentation of the tracks of two different barn owls. As in Fig 8 The two top panels are plots of the clusters in PC1/PC2 space and the two lower panels are plots in Mean-speed/Mean-turning-angle space. A color spectrum is used to depict the smallest (blue end) to largest (red end) of segments within clusters. The centroid arguments of each cluster (Eq 15) are listed in Table 2. The colors have been selected to reflect a spectral scale of the largest to smallest mean speed for each cluster’s centroid, though the cluster numbers (Column C in Table 2) are set by the clustering algorithm. Note that mean speed, as plotted here is 10 times smaller than the mean speeds 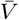 recorded in Table 2: this resizing was made to avoid extra zeros after the decimal points the in the Table. Also note that the triangular distribution of points in bottom two graphs arises because smaller turning angles are associated with the occurrence of segments with faster average speeds. Depictions of 130 randomly selected segments from each cluster for the Adult female and Juvenile male segmentation cases can be seen in Appendix A (Supp info) in Figs A.10-A.17 and Figs A.18-A.25 respectively.

**Table 2:** The centroid arguments (Eq 15) obtained from a hierarchical cluster analysis (with *k* = 8 clusters labeled C = 1, *…*, 8, see legends in Fig 7) analysis of a segmentation of tracks from two different barn owls. The results are listed in descending size of average speed (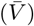) (m/s) for the centroid of each cluster. The turning angles are proportions of *π*, and the displacement are proportions as described in the methods. Depictions of 130 randomly selected segments from each cluster for the Adult female and Juvenile male segmentation cases can be seen in Appendix A (Supp info) Figs A.10-A.17 and Figs A.18-A.25 respectively.

| **C** | **Segments**<br>**# (%)** | $\bar{V} \pm \overline{SD}^{\dagger}$<br>(m/s) | $\vert \Delta\Theta\vert \pm \overline{SD}^{\vert \Delta\Theta\vert \dagger}$<br>(units $\pi$) | $\bar{\Delta}^{\rho} \pm \overline{SD}^{\Delta^{\rho} \dagger}$<br>(proportion) |
| --- | --- | --- | --- | --- |
| **Adult female** |  |  |  |  |
| 8 | 4530 (5.2) | 2.80±1.81 | 0.24±0.23 | 0.79±0.12 |
| 5 | 11655 (13.3) | 1.07 ± 0.68 | 0.18±0.15 | 0.13±0.06 |
| 4 | 12670 (14.4) | 0.79 ± 0.52 | 0.25 ± 0.22 | 0.27±0.06 |
| 1 | 19641 (22.4) | 0.76±0.47 | 0.27±0.25 | 0.10±0.04 |
| 6 | 9278 (10.6) | 0.71±0.54 | 0.33±0.28 | 0.42±0.08 |
| 2 | 11639 (13.3) | 0.69±0.42 | 0.36±0.31 | 0.73±0.03 |
| 3 | 10450 (11.9) | 0.61±0.38 | 0.39±0.30 | 0.22±0.04 |
| 7 | 7895 (9.0) | 0.61±0.36 | 0.46±0.31 | 0.12±0.05 |
| **Juvenile male** |  |  |  |  |
| 8 | 3622 (5.8) | 2.55±1.75 | 0.24±0.23 | 0.77±0.12 |
| 7 | 2679 (4.3) | 0.76±0.64 | 0.35±0.28 | 0.49±0.06 |
| 3 | 3553 (5.7) | 0.72±0.45 | 0.16 ± 0.15 | 0.09±0.04 |
| 2 | 7730 (12.4) | 0.59±0.40 | 0.24±0.21 | 0.28±0.07 |
| 4 | 9446 (15.1) | 0.51±0.31 | 0.44±0.32 | 0.08±0.04 |
| 1 | 21029 (33.6) | 0.50±0.30 | 0.28±0.24 | 0.12±0.06 |
| 5 | 5956 (9.5) | 0.50±0.33 | 0.41±0.30 | 0.29±0.06 |
| 6 | 8484 (13.6) | 0.46±0.27 | 0.38±0.29 | 0.16±0.05 |

<sup>†</sup>This is the average of the standard deviations reported for each segment in the cluster
<sup>‡</sup>This is the standard deviation of the normalized net displacement across all segments in the cluster

We see from Table 2 that the average speeds of the cluster of fastest (i.e., largest) segments for the adult female and juvenile male (C=8 in both cases) are 2.80 and 2.55 m/s respectively. For both individuals, we see in Table 2 that these “fastest clusters” (as represented by 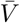) are around 3 times faster than the next fastest clusters (compare first and second rows of values in Table 2), with the fastest clusters containing only 5-6% of all segments. For both individuals the bulk of the segments have speeds that lie around the geometric means of the fastest and slowest clusters of segments (i.e., between 0.35-0.8 m/s), are very similar in size but these intermediate speed clusters highly variable in their shape: some are relatively open (say, net displacement 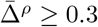) and some are relatively closed (say, net displacement 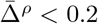). The slowest clusters of the adult female and juvenile male have the relatively low speeds of 0.10 and 0.15 m/s respectively, but these slowest adult female segments are much more open on average (C = 7, 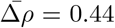) than the juvenile male segments (C = 6, 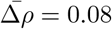). The reason for this requires a much closer look at the locations of these segments on the actual landscape where they occurred.

Depictions of 130 randomly selected segments from each cluster for the Adult female and Juvenile male segmentation cases can be seen in Figs A.10-A.17 and Figs 18-A.25 (Appendix A, Supp info) respectively. These depictions cannot be compared across segments for size because the axes in each panel of each figure have been automatically generated by our plotting routines. They also do not provide any landscape details.

Such considerations remain a subject for future studies once analyses of the movement tracks of multiple individuals of each type have been undertaken.

As somewhat expected, a distinct correlation is evident between the mean speed and mean turning angle of a segment. This expresses itself through the lower triangular distribution of points in the mean-speed/mean-turning-angle plots in Fig 8. This correlation also appears in the simulated data as well, but it is much less obvious: it is indicated by the negative slope of a band of points (Fig 7) rather than by a triangular distribution of points. The band implies that segments with lower speed in the simulation data have larger turning angles, while in the empirical data, lower speed still allows for a range of turning angles.

## 6 Discussion

The information that can be extracted from the movement path of an individual is akin to decoding a string of highly degraded symbols without an accompanying Rosetta stone to interpret the meaning of these symbols. In fact, interesting albeit superficial commonalities and differences can be drawn between reading a book and “reading” the life-time track of an animal’s story from its birth to its death. In a book, pages are spatially well-defined objects, as are the temporally well-defined diel activity routines of the lifetime track of an animal (Fig 1). Canonical activity mode (CAM) segments within a diel activity routine (DAR) are like the sentences on a page and “activity types” represented by strings of CAM segments are like paragraphs. Unlike a book, however, where every letter is clearly visible, the letters (FuMEs) of an animal track are not at all visible, only hints of letters at regularly “spaced” (in time) intervals. Thus, instead of being able to see the words as we can in any book, we can only conjure up poor images of ersatz words in the form of StaMEs (Statistical Movement Elements, Fig 1). The book metaphor for lifetime tracks, however, is too simple in one very important way: books start with blank pages upon which letters are then printed. Tracks are generally imposed upon highly structured rather than blank landscape and the underlying landscape structure plays a decisive role in determining the spatiotemporal characteristics of the tracks that are imprinted on it. [65, 66, 67].

In some ways, our Rosetta stone is our simulation model. It allows us through simulations to see how different kernels produce segments with particular sets of statistics. We illustrated this in Section 5.1 where we showed, for example, that a kernel with minimum- and maximum-step-length and range-of-turning-angle parameters values *r*^min^ = 0, *r*^max^ = 5, and *ψ* = *π/*2 creates a cluster of segments whose centroid has mean step-length and turning-angles (standard deviations in parenthesis) of 3.38 (±1.23) and 0.27*π* (±0.15*π*) (Eq 17).

More ambitiously, we could numerically construct 2-patch mappings, as defined in Eq 16, where Table 2 represents the set of centroids 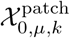 (Eq 15) obtained under a hierarchical 8-cluster, *μ* = 15 point-segmentation of the simulated data (using 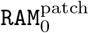) when the arguments of this mapping are (see Eq 17) the simulation parameters listed in Section 5.2. In the case of trying to match movement modes to empirical relocation data, we can use the actual landscape associated with those data, plus set selection functions fitted to different parts of the landscape (e.g., movement within and between areas where particular types of activities take place). This appears to be a challenging process where it remains to be seen in future studies how the methods discussed in this paper can be applied to identifying putative movement modes that appear to be used by individuals when moving across real landscapes and carrying out a range of activities.

When comparing the diversity of the clusters obtained from the adult female (AF) and juvenile male (JM) barn owl tracks, as reported in Table 2, with that of the simulated data, as reported in Table 1, its clear that our StaME approach has the potential to match real with simulated data. For example, when using 10-step segments the range of values is nearly 10-fold for velocity, 4 to 5-fold for absolute turning-angle, and 8-fold for relative net-displacement. For the 30-step segments these fall to 6-fold for velocity, 3-fold for absolute turning-angle, and 6-fold for relative net-displacement. Comparable ranges for the barn owl are around 5-fold for velocity (somewhat less for the AF and more for the JM), around 3-fold for absolute turning-angle, and 8-10 fold for net displacement (lower end for AF, upper end of JM). Thus our simulations show a greater velocity range and a greater absolute turning-angle range, but a smaller relative net-displacement range than our barn owl data. The larger barn owl net-displacement range arises because the owls typically return home each day, but displacement to another resting site occurs on some days [42]. To capture this behavior in our ANIMOVER_1 simulator, we would need to add a movement kernel that is biased to move in the direction of a homing beacon at certain times during the diel cycle.

The challenge should not be underestimated when it comes to demonstrating how a bottom-up StaME approach to constructing and analyzing movement track structures, when combined with fitting auto-regressive models (AR(p) where p is the depth of the time delay dependence), can be used to fit models to movement patterns at the subdiel and diel scales. SDE methods have typically been used to estimate macro level quantities emerging from the movement behavior of individuals and populations, such as home range, speed, and distance travelled [68, 69]. To keep things simple, however, these methods avoid the complexities of dynamic background environments, which are known to greatly influence movement behavior [70]. Bottom up hierarchical modeling approaches, however, by linking movement to environment using step-selection functions [54], provide a direct way of incorporating environmental covariates into movement behavior. This behavior is complicated by the fact that an individual’s internal state variables (motivations linked to time-of-day and time-of-year factors, hunger, thirst, fear, etc.) are also critically important [71] Ultimately these factors will need to be included in simulation models used to predict the movement patterns of individuals at the level of diel and subdiel movement patterns. Incorporation of these, which remains a challenge for future studies, will require fitting auto regressive models to the sequencing of CAMs with movement mode switching probabilities that are functions of all critical covariates that individuals use to decide on where to move next as their movement track relocation time series unfolds under various environmental conditions.

Of course, other approaches to simulating movement have been developed, several of which are mathematically more sophisticated than the approach we take. Some of these treat movement as continuous-time stochastic differential equation (SDE) process for which Brownian motion (purely random movement) and Orstein-Uhlenbeck processes (directionally correlated random walks) and modifications thereto (e.g., velocity correlated walks, central attractors, etc.) [72, 73, 74, 75, 76] are examples. Others represent movement in terms of partial differential equations some random components that switch between gradient following search (advection plus random noise) and random search [77] or discrete approaches using integrated step-selection functions [78]. In addition, machine learning methods have been shown to outperform SDE models for predicting the next location of individuals in some systems [79].

It is worth noting that spatio-temporally continuous models are from a computational point of view also discrete in requiring discretization schemes to generate numerical solutions. In such cases, however, the aim is to keep the discretization small enough and solution methods robust enough so the associated discretization errors do not affect interpretations of outcomes. Formulating movement models discretely, as we have, with regards to space and time avoids the issue of what is an appropriate solution method, but introduces both philosophical and practical issues regarding differences in discrete-versus continuous-time representations of ecological processes [80], particularly since data always discretized in one way or another [81]. Additionally, discrete time, space, and variable trait formulations of ecological processes facilitate the incorporation of idiosyncratic details that elude the more rigorous formulation of continuous variable formulations. This is particularly true when it comes to turning model formulations into computational code for analysis through numerical simulation. Such formulations are naturally facilitated when movement is embedded into network structures [82, 83] or is modeled as taking place on cellular arrays [84, 85, 86, 87].

Given the complex shapes of the segments plotted in Figs A.2-A.25 (Appendix A, Supp info), the hierarchical clustering approach that we have taken to parsing the movement track segments into categories may not be the best approach. Supervised and unsupervised deep-learning approaches provide less forced and more powerful ways to extract clusters, particularly those targeted at time series data [88, 89, 90, 91, 92, 93]. In terms of supervised, deep-learning approaches, convolutional neural nets (CNNs) of appropriate types may be trained using segments simulated by ANIMOVER_1 over real landscapes, obtained through the application of various sets of movement rules, as a way of generating the training sets [94]. Once trained these CNNs can then be used to classify empirical data segments thereby providing some insights into the types of movement rules that may have been used by actual individuals when moving over the landscapes in question. Further, we note that rectangular looking elements illustrated in Figs A.4 and A.5 are a function of the scale of the discretized landscapes over which trajectories are simulated or, for that matter analyzed, in terms of the smallest step size invoked or time interval used to record consecutive locations. However, no matter what scale is selected, the smallest StaMEs—i.e., those associated with resting behavior—will be limited by both the level of discretization used in simulating data or the errors associated with measuring locations using GPS or reverse GPS technologies [95]. In addition, the scale of discretization is ultimately limited by the consistency/stability trade-off of numerical computations [96]. It should also be added that various measures can be taken to correct errors associated with the measurement of locations using techniques to smooth out data or apply Kalman filtering, but this may lead to some loss of temporal resolution in the underlying data (e.g. see [21]).

## 7 Conclusion

In the absence of suitable high frequency data, insights into the ecological underpinnings of the movement tracks are currently being obtained by studying them the hierarchical level of behavioral activity modes (BAMs), [21], diel activity routines (DARs) [42, 41, 97] and above [98, 99]. The reason for this is its fixed temporal length of 24 hours, which makes it possible to segment in a relatively unambiguous way. The only ambiguity is what point in time during the diel period should be fixed as the start/end point for each DAR. This problem has been discussed elsewhere [100, 101]. Additionally, DARs can be strung together to obtain supra-diel constructs (Fig 1) such as the extent of seasonal range of individuals [102, 32], and beyond this to classifying the syndromic movement behavior from whole or abbreviated lifetime tracks [99].

Next steps to follow on from the work presented here likely include studies to evaluate the most appropriate clustering methods to be used to classify segmentation types at different levels of segmentation (e.g., StaMEs to LiTs, Fig 1). It should also include studies of how a bottom-up StaME approach to constructing and analyzing movement track structures, when combined with fitting auto-regressive models (AR(*p*) where *p* is the depth of the time delay dependence), can be used to fit models to higher order segments, particularly variable length BAMs and how this compares with the more direct approach of identifying BAMs using behavioral change point analysis (BCPA) and hidden Markov models (HMM).

Over the next decade we can expect exponential increases over time in the availability of high-frequency movement relocation time series for many different species, reflecting both growing interest in the topic, and technological improvements in tracking methods. This increase, together with the increasing computational power of server clusters, the use of parallel processing, and improvements and extensions to our ANIMOVER_1 simulator will make the methods discussed in this paper for deconstructing diel activity routine tracks into StaMEs more reliable and easier to implement. Additionally, deep learning methods and possibly some other machine learning methods [94, 90, 91, 92, 93, 103] that are able to account for complexities in the shapes of the segments arising from our analysis (e.g., see Figs A.2-A.25 in Appendix A, Supp info) may be more useful for categorizing segments than the hierarchical clustering approach used here. It will also facilitate using StaMEs to bring movement canonical activity modes (CAMs) and higher level behavioral activity modes (BAMs) or movement syndromes [104, 32] into sharper focus using various time series forecasting techniques [105, 106] and more sophisticated movement simulation algorithms.

## Glossary

For the convenience of the reader we provide a glossary of indices and symbols in Table 3.

**Table 3:**
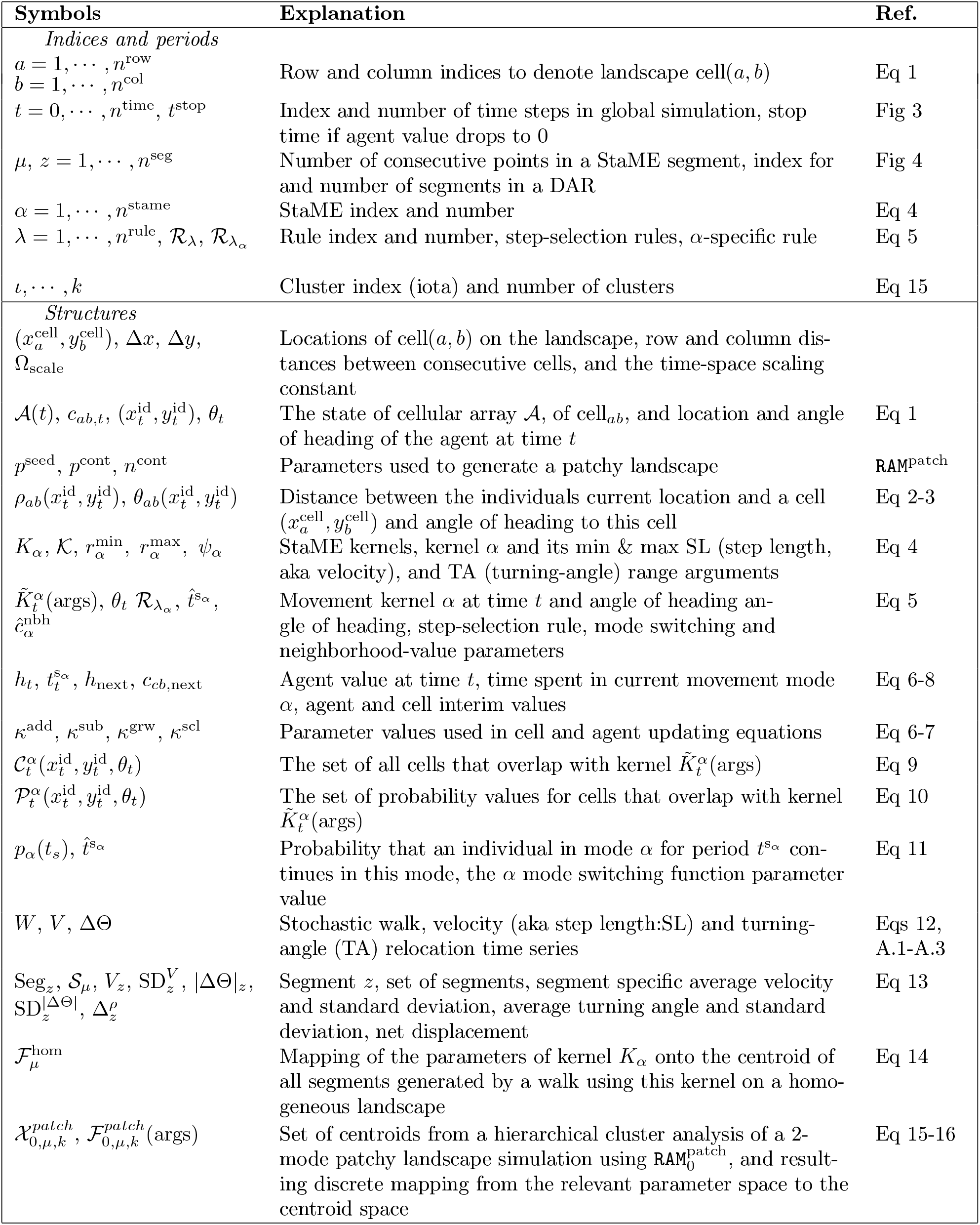
Indices and symbols used to formulate the model’s structure.

## Acknowledgments

Assaf Uzan, Michal Handel, Nadav Amir, Shani Cain and Yohay Wasserlauf helped with fieldwork and other components of data collection. We are also grateful to many people and institutes who permit us to use their towers and power supply for our base stations. These include Eli Chupa and Pablo Uner (Harod Valley Water Supply Co-op LTD), Shmuel Wirzberger (Kibbutz Beit Hashita), Eytan Ofer (Kibbutz Sde Eliyahu), Yitzhak Rivlin, Yado Shalev and Elon Shaked (Kibbutz Ein HaNetziv); Shaldag Refrigeration Storeroom LTD, Avihu Levanon (Partner PHI LTD), Michael Zelmanov (Pelephone Communications Ltd), Asher Nahmani and Rafi Levi (Mekorot), and Eli Cohen and Eli Mimran (Bezeq Inc.), and The Israeli Telecommunication Corp Ltd.

## Authors contribution

WMG designed the study, formulated the model, and drafted the manuscript

RS built the Numerus coding platform and RAMP simulators

VS carried out the StaME extraction and clustering procedures

OS contributed to model design

SC, RN, OS, and ST were involved in the setting up and running the ATLAS system and collecting and organizing the barn owl relocation data

All authors read and edited various versions of the manuscript including the final version

## Declarations

### Ethical Approval

Trapping and tagging procedures were authorized by permits 2019/42155 and 2020/42502 from Israel Nature and Parks Authority.

### Consent to participate

Not applicable.

### Consent to publish

Not applicable.

## Funding

This work was funded in part by the A. Starker Leopold Chair of Wildlife Ecology at UC Berkeley, by the Minerva Foundation, the Minerva Center for Movement Ecology, and grants ISF-965/15, 1919/19 and 396/20, from the Israel Science Foundation. Further, support was given by a grant from the Data Science Center at Tel Aviv University (TAD), and by the Koret-UC Berkeley-Tel Aviv University Initiative in Computational Biology and Bioinformatics (WMG, OS, ST). SC was also supported by the Hoopoe Foundation, Society for the Protection of Nature in Israel, Ministry of Agriculture and Rural Development, Ministry of Regional Cooperation, Larry Kornhauser, and Peter and Naomi Neustadter.

### Availability of data and materials

The simulation data can be found in the supplementary online file `Two_Kernel_Movement.csv`. The data used for the barn owl analysis is available at the Github repository https://github.com/LudovicaLV/DAR_project, as provided in [42]

## Appendices

These appendices contain references to equations and tables appearing in the main text of *Getz et al*., *The Statistical Building Blocks of Animal Movement Simulations* and the bibliographic citations herein are included in the *References* section of that text.

### A Hierarchical Segmentation and Empirical Data

#### A.3 Issues of scale

We have various temporal scales in our model as they relate to the following structures and processes from likely fastest to slowest:

##### 1 FuME scale (fundamental movement element)

The scale of the actual “hidden” movement elements underlying the StaMEs (previously called metaFuMEs). This is the average time it takes to perform a typical FuME, such as one repeatable sequence when striding, wing flapping, wiggling or undulating one’s body.

##### 2 Landscape traversal scale

This converts the landscape pixel scale to the typical time it takes for individual to move across one pixel. This scale is typically finer than the relocation data scale (see next), though if the pixels are large (e.g., 10 meters or more) and the relocation data scale fine then this scale may be coarser than the relocation data scale.

##### 3 Relocation data scale

This is set by the frequency of the relocation data. Fine scale implies that data points are less than 10 secs apart. Course scale implies points more than 1 minute apart. Intermediate scale is between these two.

##### 4 StaMe scale (statistical movement elements, formerly metaFuMEs)

This is the scale of our StaME elements—i.e., it relates to the number of relocation data points (including missing points) used to compute our StaMEs, which my vary from typically 10 to 30 points, depending on the fineness of the relocation data scale.

##### 5 CAM scale (Canonical activity mode) (formerly short or homogeneous CAMs)

This scale relates to the typical length of a homogeneous sequence of some underlying metaFuMEs. This scale will vary somewhat, depending on the type of behavior. Directed walking, for example, may last several hours, though it will likely be interspersed with pauses when the individual rests or scans their surrounding for information relating to navigation or safety. Grazing, which is a mix of several different FuMEs that each last a short time (e.g., stop-start motions involving forward, sideways and backward movements coupled with various head motions) may also last on the order of large fractions of an hour or multiple hours.

##### 6 Activity type scale (formerly long CAMs)

When relocation data is relatively fine (consecutive points are seconds apart) then “activity types” may be identified in terms of a characteristic mix of CAMs. Thus a migration movement CAM that lasts for a substantial part of a diel may be made up of a characteristic mix of directed movement, grazing, and resting CAMs. On the other hand, if the relocation data scale is relatively course (i.e., relocation points are minutes apart) then StaMEs reflect average movement behavior during “migration movement” may already have statistics that reflect a mix of directed movement, grazing, and resting behaviors.

#### A.2 Movement path data and segmentation

Here is a refinement of the hierarchical definitions provided in [30] to help us understand how to parse out the information contained in the planar points relocation time series or walk (recall Eq 12)

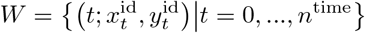

From this time series, the following step length *S*, or equivalently velocity *V* and turning-angle ΔΘ time series, can be derived using the relocation point frequency *F*

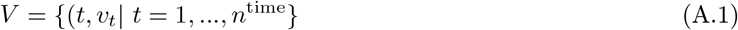

using the derived values

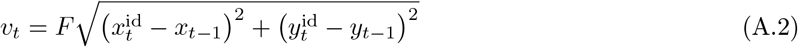

and

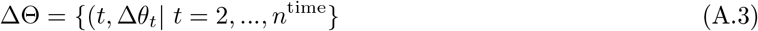

using the derived values

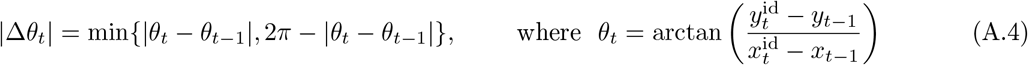

and the sign of |Δ*θ*_*t*_| is easily determined by considering which quadrants *θ*_*t−*1_ and *θ*_*t*_ are in and whether moving clockwise or anticlockwise subscribes the smaller of the two angles between them: anticlockwise implies positive and clockwise negative. As in [18], we may also study movement in terms of the persistent (*v*^per^) and tangential (*v*^tan^) velocities rather than speed (*v* or stepsize) and turning angle (*δθ*), using the transformations

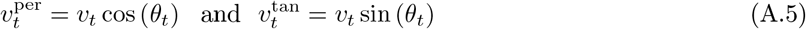

**FuME** A FuME is a fundamental movement element that for some types may be hard to observe its start and end points, or even define with any precision

**Step** A step is one point *t* in the time series *W* with which we associate a step length *s*(*t*) and turning angle Δ*θ*(*t*)

**StaME** (previously called metaFuMEs) These are statistically entities computed from consecutive segments of fixed length of a walk *W* (Eq 12) (i.e., each segment contains a specified number of consecutive relocation points—typically 10-30 to obtain reasonable statistics). These segments can be classified into a few different StaME categories or types based on their statistics—typically the mean and variances of the persistent and turning velocities *v*(*p*) and *h*_*t*_ obtained for each segment.

**CAM** A canonical activity mode, formerly referred as a “short” or “homogeneous” CAM that is a consecutive sequence of two or more StaMEs of the same type.

**Activity type** Formerly referred to as a long CAM, and is a characteristic mix of CAMs defining a complex movement activity such as grazing (a mix of stop-start movements)

**DAR** A diel activity routine with start and stop points in its 24 hour cycle selected to correspond, for example, to its major resting period.

**LiMP** A lifetime movement phase is a sequence of DARs that occurs during the individuals life history. It may be a migration event that occurs over several days or weeks, or it may be a seasonal movement pattern (e.g., a nomadic phase, or a home ranging phase) that repeats annually.

**LiT** The lifetime track of an individual from the moment it starts moving after birth to the moment it ceases to move at death.

#### A.3 Step-selection procedures

ANIMOVER_1 includes two movement modes, each involving its own step selection procedure. One specifies the movement of individuals within patches (wp) and a second between patches (bp). Each of these two modes involves a kernel with mode-specific parameters 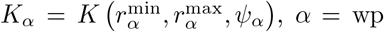, bp (Eq 4), as well as mode specific time-in-mode parameter 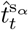 (i.e. the value of 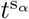 at time *t*) and neighborhood value parameter 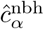. The procedures also involve updating individual and cellular array states using Eqs 6 or 7, depending on which of the `RAM`s (see Section 4.2) have been selected.

##### Within patches step-selection procedure, 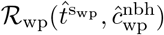

ℛ_wp_.1 *SET α*_*t*_ = *α*_wp,next_, 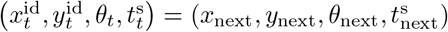, and (*c*_*ab,t*_, *h*) = (*c*_*ab*,next_, *h*_next_) (Fig 3).
ℛ_wp_.2 *COMPUTE* the values of the admissible set of cells 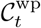 (Eq 9) and *COMPUTE* 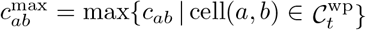. Note, for the case `topology`=`plane`, there may be no admissible cells.
ℛ_wp_.3 *COMPUTE*

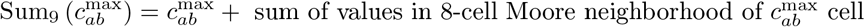
ℛ_wp_.4 *IF* Sum_9_ 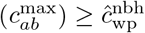 *THEN*
  a. *COMPUTE* the set of probabilities 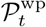 (Eq 10) associated with selecting one of the admissible cells
  b. *SELECT* the next cell, cell(*a, b*) using the probabilities in 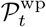 in a multinomial drawing (i.e., the cell most likely to be selected is the one with the largest *p*_*ab*_ and so on) *ELSE*
  c. *MODIFY* 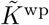 by replacing 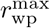 with 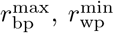 with 0 (typically we already have 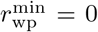), and *ψ*_wp_ with *π* and repeat steps ℛ_*wp*_.2 & 3.
  d. *APPLY* rule ℛ_wp_ 4a & b to the selected cell.
  e. *IF* the updated Sum_9_ 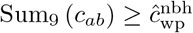 *THEN* move to this cell
  f. *ELSE* move at random to one of the cells on the rim of a circle at radius 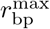 from the current location (i.e., all points whose distance from the current location is within the band 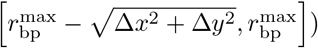. We note that this step will also prevent an individual from getting stuck at a boundary when `topology` = `plane`.
ℛ_wp_.5 Now that we have identified the next cell(*a, b*) to be occupied:
  a. *SET* 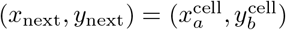.
  b. *COMPUTE* the next angle of heading *θ*_next_ (Eq 3) and distance moved *ρ*_*ab*_(*x*_next_, *y*_next_) (Eq 2).
  c. *UPDATE c*_*ab* next_ and *h*_next_ using 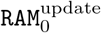 (Eq 6) or 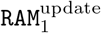 (Eq 7) or a user specified 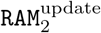
ℛ_wp_.6 *UPDATE* the StaME index *α*_next_ using probability *p*_wp_(*t*^s^) (Eq 11) as follows (where Sum_9_ (*c*_*ab*_) is sum of values in the 8-cell Moore neighborhood of Cell(*a, b*) plus *c*_*ab*_ itself) *IF Z* ∼`BINOMIAL`[*p*_wp_(*t*^s^)] = 1 *AND* 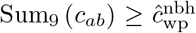 (if updated under ℛ.4c use latest value) *THEN α*_next_ = wp *ELSE α*_next_ = bp

##### Between patches step-selection procedure, 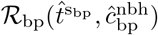

ℛ_bp_.1 *SET* 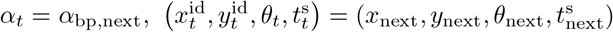, and (*c*_*ab,t*_, *h*) = (*c*_*ab*,next_, *h*_next_) (Fig 3).
ℛ_bp_.2 *COMPUTE* the values of the admissible set of cells 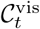 (i.e., using 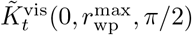 rather than 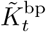 in Eq 9) and *COMPUTE* 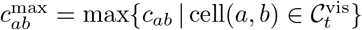. Note, for the case `topology`=`plane`, there may be no admissible cells when the individual is on the boundary near a corner. In this case, when no visible cells are available, then use 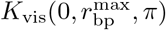, i.e., the individual now looks behind itself as well.
ℛ_bp_.3 *COMPUTE*

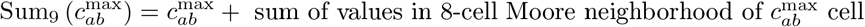
ℛ_bp_.4 *IF* Sum_9_ ℛ *THEN*
  a. *COMPUTE* the set of probabilities 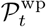 (Eq 10) associated with selecting one of the admissible cells
  b. *SELECT* the next cell, cell(*a, b*) using the probabilities in in a multinomial drawing *ELSE* move to any cell at random in the set 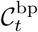 (i.e., overlapping with 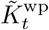).
ℛ_bp_.5 Now that we have identified the next cell(*a, b*) to be occupied:
  a. *SET* 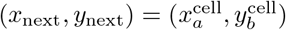.
  b. *COMPUTE* the next angle of heading *θ*_next_ (Eq 3) and distance moved *ρ*_*ab*_(*x*_next_, *y*_next_) (Eq 2).
  c. *UPDATE c*_*ab* next_ and *h*_next_, but for this movement mode setting *κ*^add^ = 0 in the same `RAM`^update^ used in ℛ_wp_.6
ℛ_bp_.6 *UPDATE* the StaME index *α*_next_ using probability *p*_bp_(*t*^s^) (Eq 11) as follows (where Sum_9_ (*c*_*ab*_) is sum of values in the 8-cell Moore neighborhood of Cell(*a, b*) plus *c*_*ab*_ itself) *IF Z* ∼`BINOMIAL`[*p*_bp_(*t*^s^)] = 1 *AND* Sum_9_ 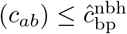*THEN α*_next_ = bp *ELSE α*_next_ = wp

#### A.4 StaMEs from empirical data

Various methods can be applied to extract StaMEs from empirical data. We have outlined one method in Section 3 of our text. Here we present more comprehensive details on the method we use. In order to separate different movement modes from each other, we perform a cluster analysis. Clustering is an unsupervised machine learning procedure which finds an optimum partition of a set of points with two or more variables into subsets (or clusters) of similar points.

Here we use hierarchical clustering approach, which results in a tree structure called dendrogram having a single cluster at the root, with leaf nodes representing data points in the set. Recall from Eq 13 that the variables 𝒮_*μ*_ had been constructed using the relocation time series *W* in Eq 12. which we repeat here for convenience:

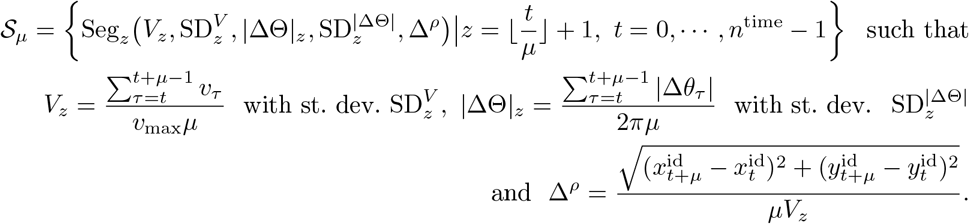

To begin with, it is important to discard any points containing missing or non-numeric values (since the variables to be used in clustering algorithm are all numeric). Further, each of these variables ranges in [0, 1], and hence no further normalisation has been carried out. Cluster analysis is performed on the segment level data (i.e., with 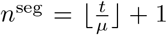 points) using the variables in 𝒮_*μ*_. Specifically, we want to find an optimal partition

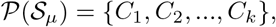

where 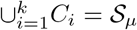, and *C*_*i*_ ∩ *C*_*j*_ =∅ (hard clustering).

Hierarchical agglomerative algorithms implement bottom-up clustering methodology, which starts with each point of 𝒮 in its own singleton cluster, followed by successive fusions of two clusters at a time, depending on similarity between them, leading to a specified number of clusters (or, alternatively, leading to one cluster followed by cutting the dendrogram at the desired number of clusters). These are deterministic, yet greedy, in the sense that the clusters are merged based entirely on the similarity measure, thereby yielding a local solution. Most of the similarity schemes are specified not in terms of an objective function to be optimized, but procedurally. We use Ward’s minimum sum of square scheme, which performs a fusion of clusters while minimizing the intra-cluster variance. Our distance metric to quantify dissimilarity is the Euclidean measure.

To perform clustering, we make use of `hclust` function from `fastcluster R` package. It replaces the `stats`:: `hclust` function, which offers the most common implementation of Ward’s hierarchical clustering in R. The conventional algorithm from `stats` package takes as input the set of points 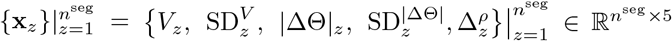 (represented by 𝒮 _*μ*_) and a pair-wise dissimilarity matrix 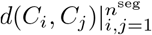 (to be computed in advance). Starting with *n*^seg^ clusters, it fuses multiple pairs of them in each of 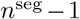steps while satisfying Ward’s criterion and updates the dissimilarity matrix with the distance of each of the clusters still available to be merged with the newly created cluster. Accordingly, a series of partitions are produced, with the first one containing singleton clusters and the last one containing all *n*^seg^ points in one cluster. At each step, a set of clusters available to be merged (active clusters) is maintained, and each merger is tracked leading to a dendrogram. Concisely, the algorithm can be represented as shown in Algorithm 1. The dendrogram thus produced can also be understood as a weighted graph, with leaf nodes representing data points, and each internal node representing the cluster of its descendent leaves. The dissimilarity between clusters is represented by edge weights. Built-in `R` function `cutree` can then be used to cut the dendrogram at the required number of clusters, and `colMeans` can be used to compute a cluster’s center.

##### Algorithm 1

Naive hierarchical agglomerative clustering algorithm

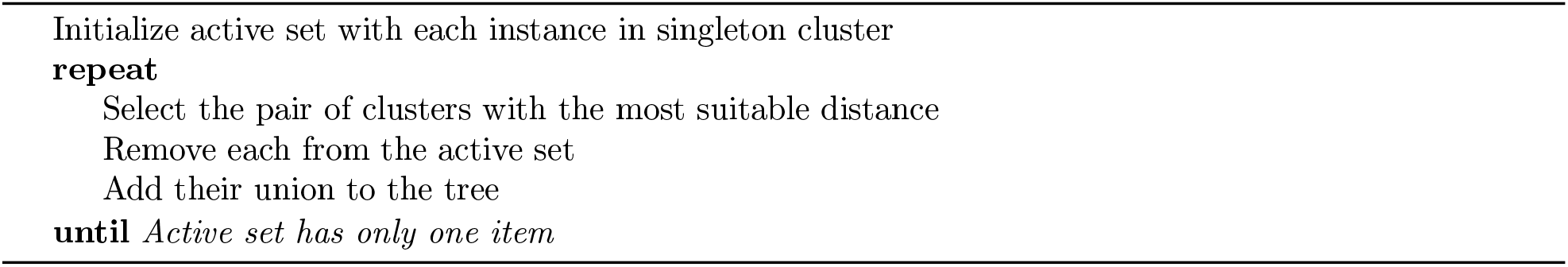

This algorithm has a time complexity of 𝒪 (*n*^seg3^ and requires Ω(*n*^seg2^) memory (on account of distance matrix computation and storage), making it unsuitable for our segment-level data set with ∼ 10^5^ points. Further, it is also difficult to distribute over multiple threads because the complete dissimilarity matrix along with active clusters and current state of dendrogram is required by all the processes. To get around these difficulties, we make two modifications to this workflow. First, we make use of `parDist` function of `parallelDist` package, which permits parallel computations of pair-wise dissimilarities. It offers the same interface as that of `stats::dist R` function. Second, we use an improved algorithm for hierarchical agglomerative clustering from `fastcluster` package, as mentioned above. The algorithm performs hierarchical clustering with Ward’s scheme faster by accomplishing the search for the best cluster to merge with any cluster in the most efficient way [107]. For the clustering procedure, we choose to not perform dimensional reduction and use all 5 variables in 𝒮 _*μ*_. This ensures enhanced interpretation of results.

As noted above, the number of clusters *k* needs to be specified in order to cut the dendrogram. As is evident from Figures 7 and 8, we selected *k* = 8 to be used in our analyses. We initially choose to err on the side of having more than fewer clusters than may turn out to be optimal because we wanted to try to isolate clusters at the extremes of the cluster groups that were as homogeneous as possible in terms of not mixing many different movement modes in one movement segment. It may also be useful in some particular cases to use a gap statistic to justify one’s choice of *k*. For example, in the two-movement mode simulation with 10-step segmentation we used a Gap statistic that compares the total intra-cluster variance for different values of *k* with the corresponding no clustering case. In this case, we see in Fig A.1 that the rate of increase of our Gap statistic somewhat flattens at *k* = 8. Accordingly, this would be the appropriate choice for our 10-step segmentation two-mode movement simulation. A relatively large value large value of *k* allows us to consider how close the centroids of different clusters would be and what it may mean to combine one or more clusters. Also, every data set may have a different preferential value for *k* when considering a gap statistic. Further, to facilitate comparisons across our analyses of different data sets, we kept to our selection of *k* = 8. As mentioned in the discussion section, hierarchical clustering is but one of many methods that may be used to cluster the data and other machine learning and even deep learning methods [108, 109, 110], can be explored, particularly when segment shapes vary widely.

Our computations have been performed on a node of Berkeley’s HPC Cluster Savio. The system is Lenovo NeXtScale nx360m5 equipped with two Intel Xeon 12-core Haswell processors (core frequency 2.3 Ghz) and 128 GB of 2133 Mhz DDR4 memory. The `R` function `parDist` distributes distance matrix calculation over all of 24 available processor cores, and returns a “`dist`” class object - the lower triangle of the distance matrix (since it is symmetric). `hclust` returns a list with several components describing the dendrogram. `cutree` method, which is used to cut the tree, returns a vector of integers indicating which cluster each point has been assigned to.

The segment level data in 𝒮 has 5 variables. Accordingly, principal components analysis is required to visualize the clustering results. This is a coordinate transformation procedure to a new set of variables such that maximum variance is preserved (which amounts to minimum loss of information). The two principal components accounting for maximum variance are used for 2-d cluster plots.

Our speed and turning-angle measurements are scaled with maximum speed and *π*, respectively, ensuring that the segment level data has all variables ranging in [0, 1]. This makes it reasonable to calculate principal components without further normalization. For better interpretation, we report the variable 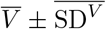 in Table 2 in units of *m/s* after rescaling with the maximum speed. PCA has been performed using built-in R method `prcomp`. To ensure no further normalization, the binary arguments `center` and `scale` in `prcomp` (which govern whether the variables are zero centered, and whether the variables are scaled to have unit variance, respectively) are set to FALSE. The function returns a list with various components which can be used to read eigenvectors and eigenvalues (and hence standard deviations of principal components) of the covariance matrix.

#### A.5 Plots of 10-step simulation data segments

Each plot in this subsection, particularly across different clusters, has had its axes for each panel automatically set by the plotting routine. Thus the size of segments is not comparable, only their relative shapes is informative.

**Figure A.1:**
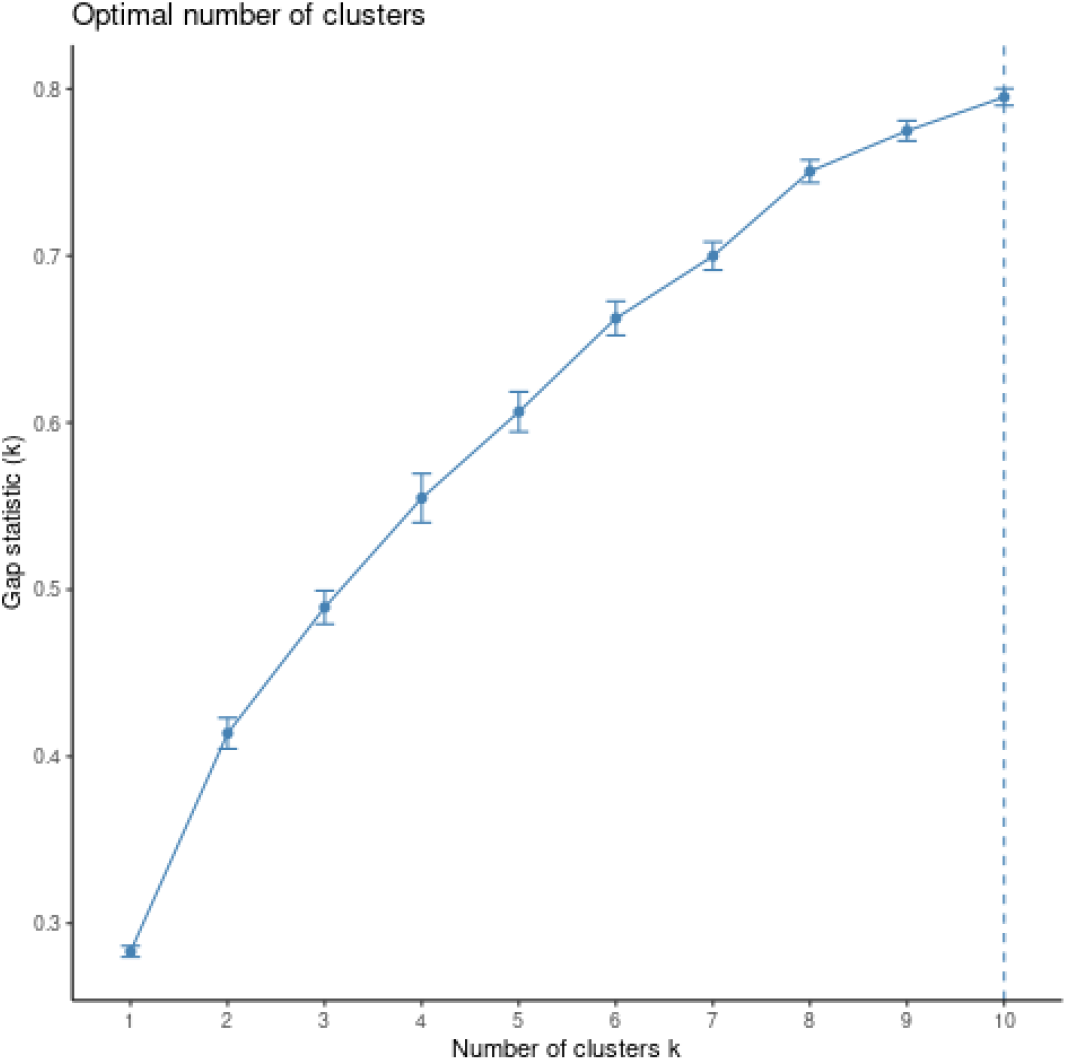
Gap statistic for the two-movement mode simulation data in section 5.2.

**Figure A.2:**
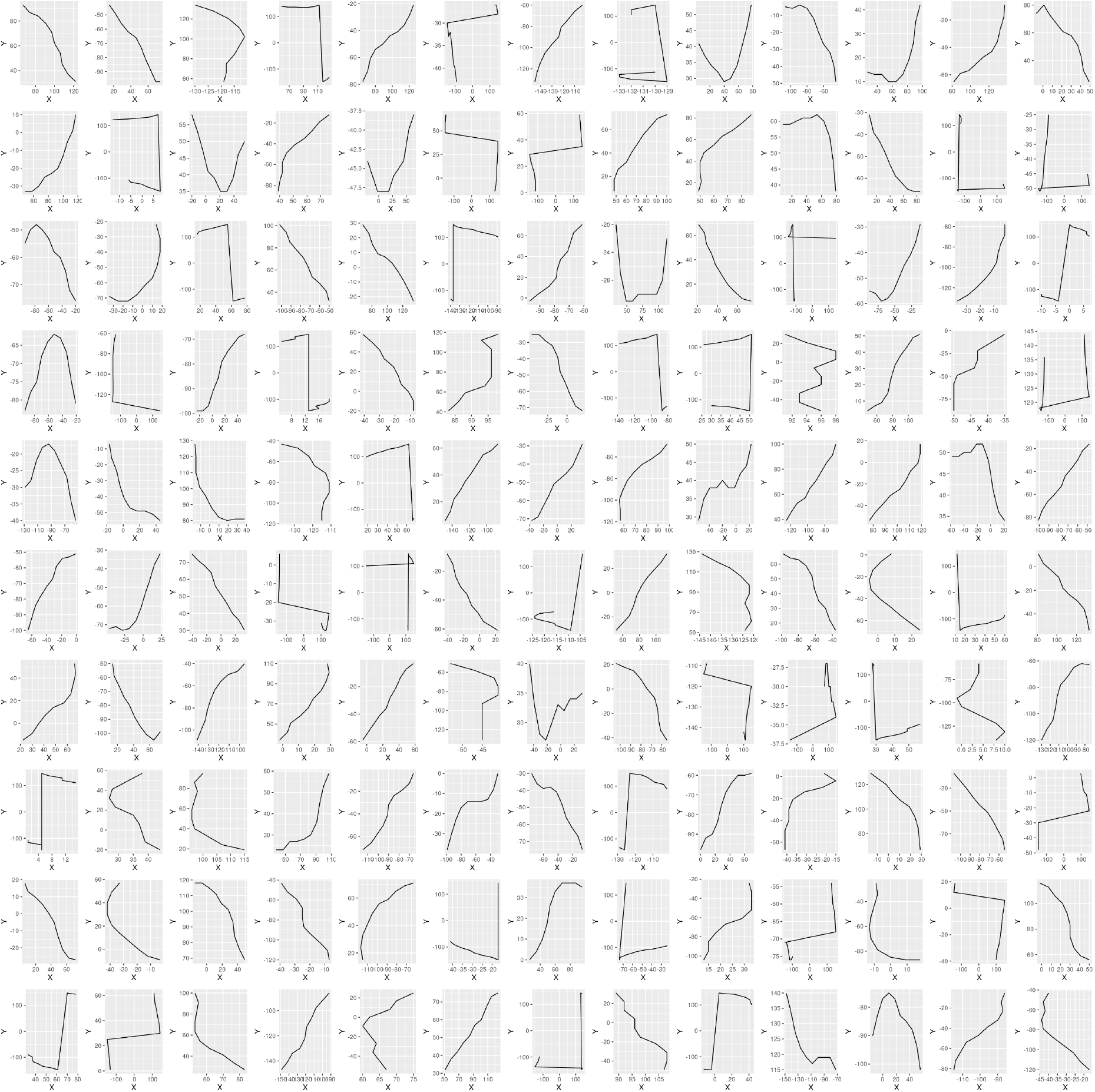
130 random 10-step segments from cluster C=1 (Table 1)

**Figure A.3:**
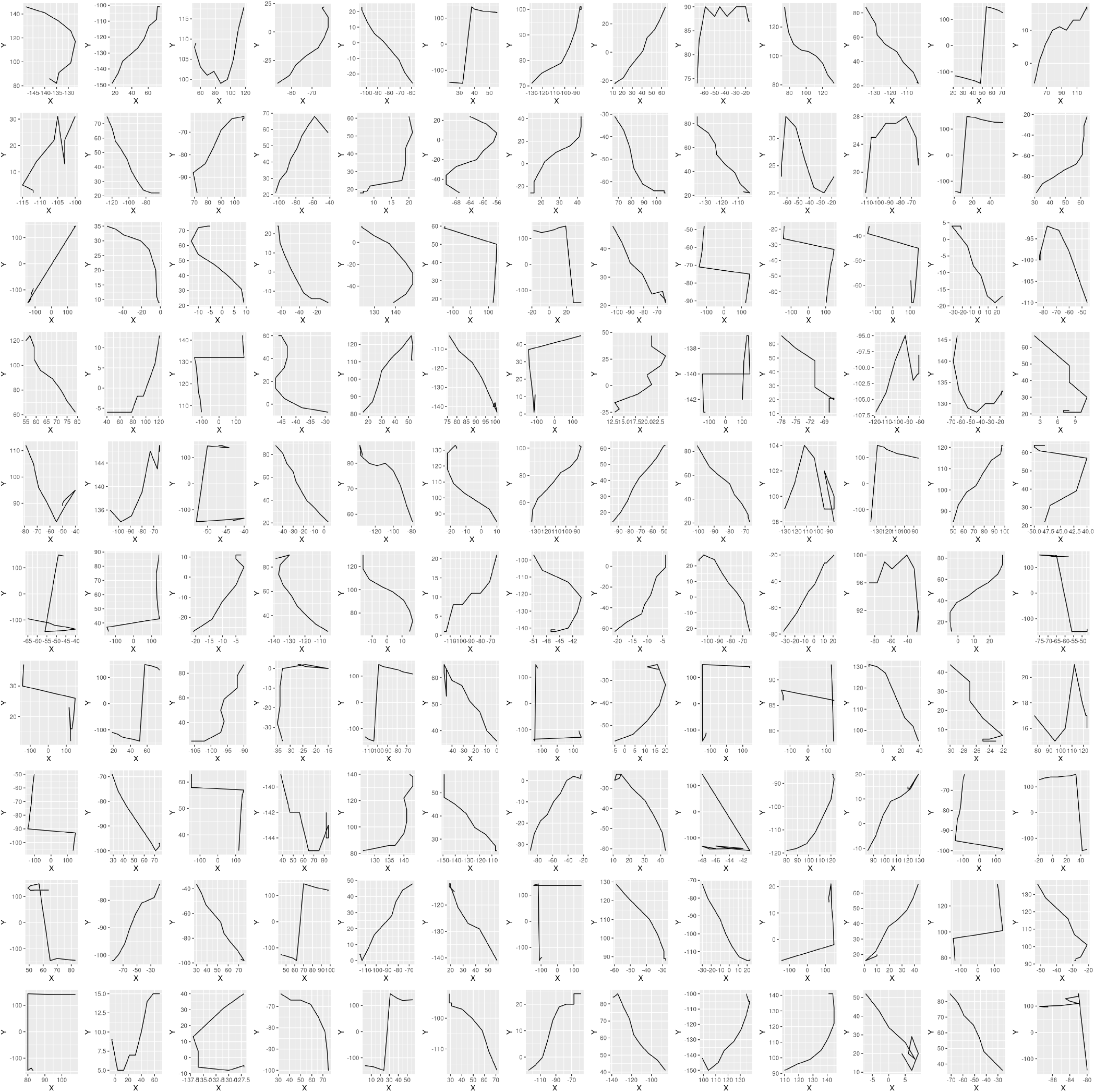
130 random 10-step segments from cluster C=2 (Table 1)

**Figure A.4:**
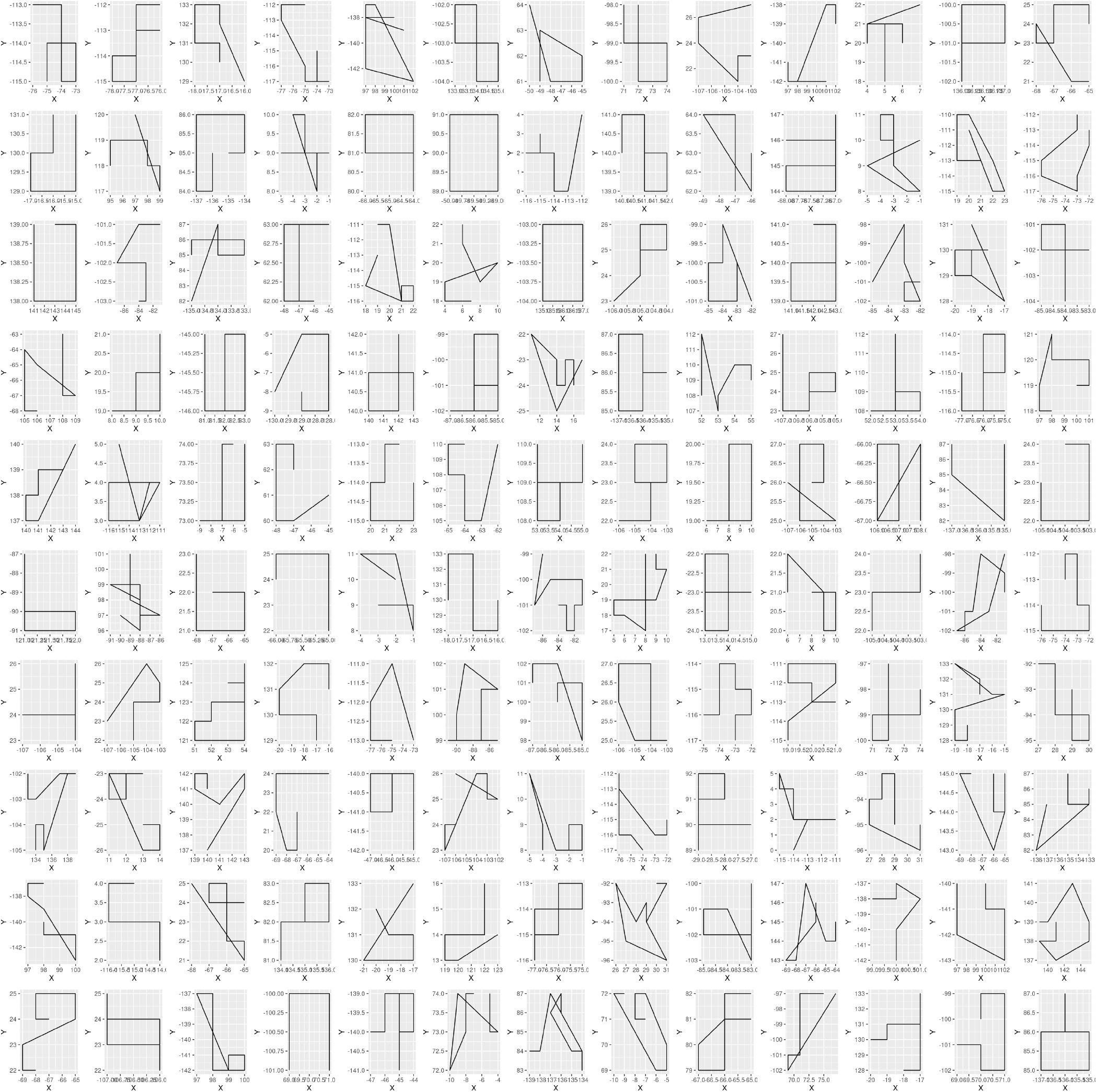
130 random 10-step segments from cluster C=3 (Table 1)

**Figure A.5:**
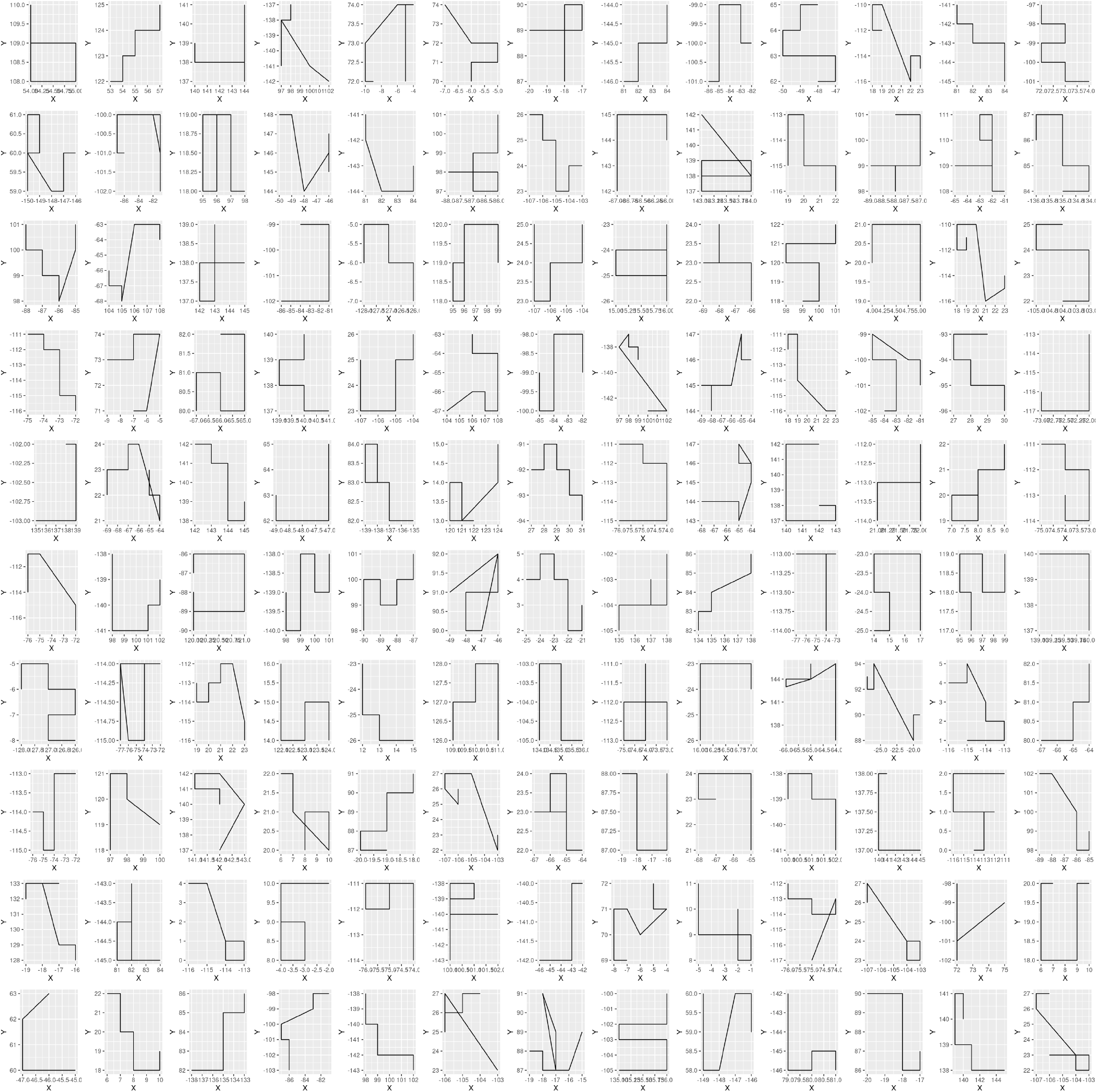
130 random 10-step segments from cluster C=4 (Table 1)

**Figure A.6:**
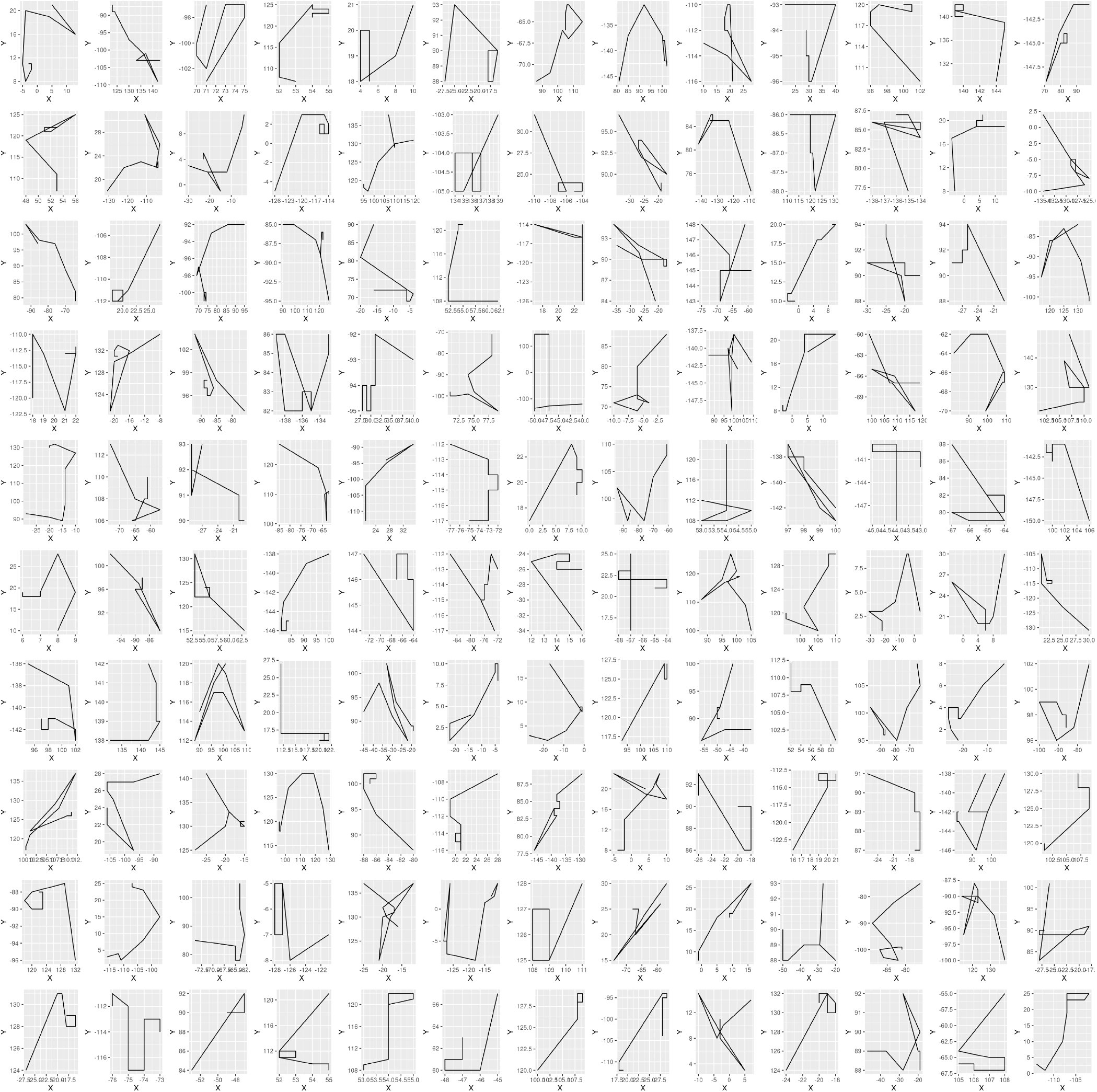
130 random 10-step segments from cluster C=5 (Table 1)

**Figure A.7:**
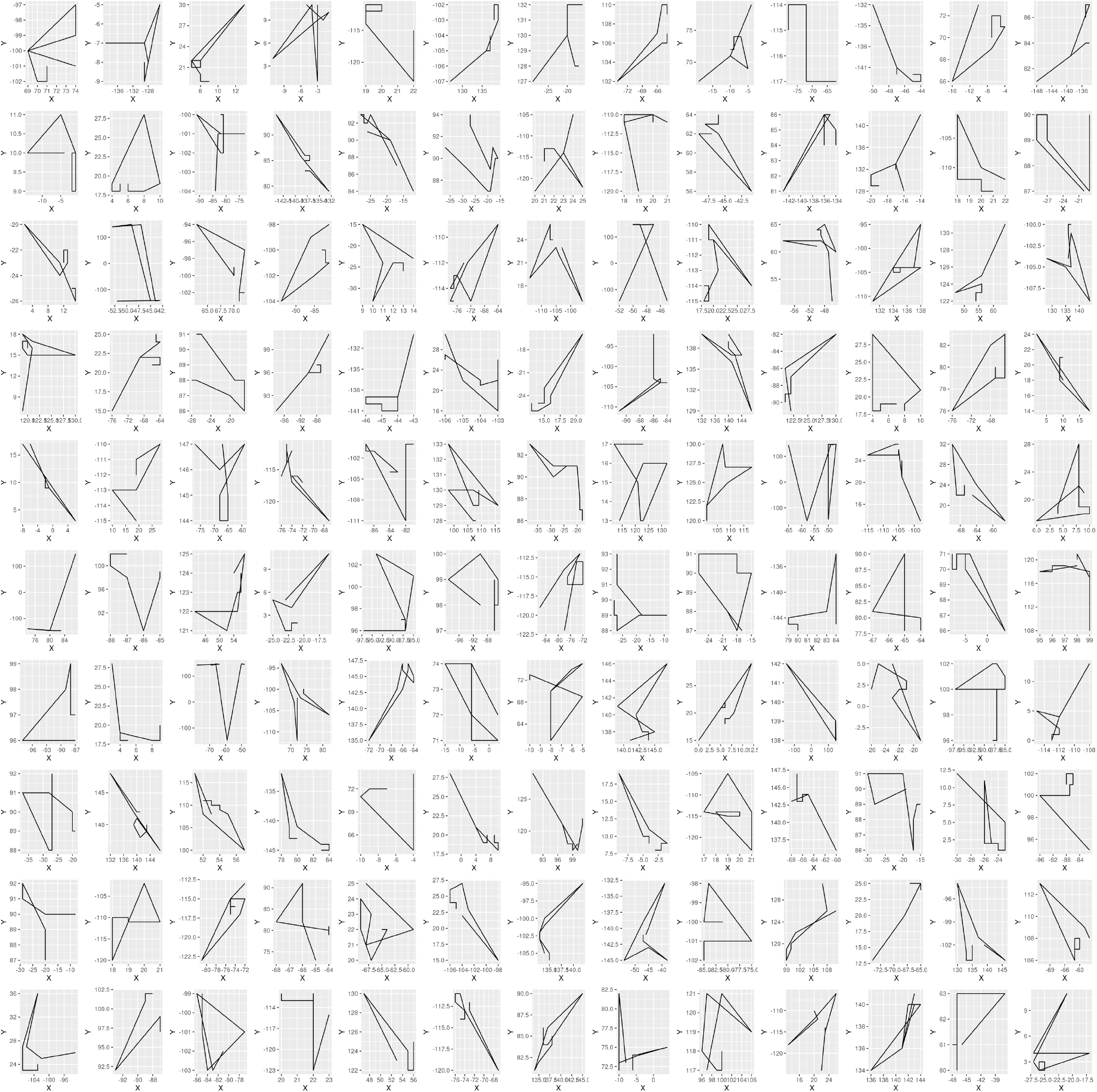
130 random 10-step segments from cluster C=6 (Table 1)

**Figure A.8:**
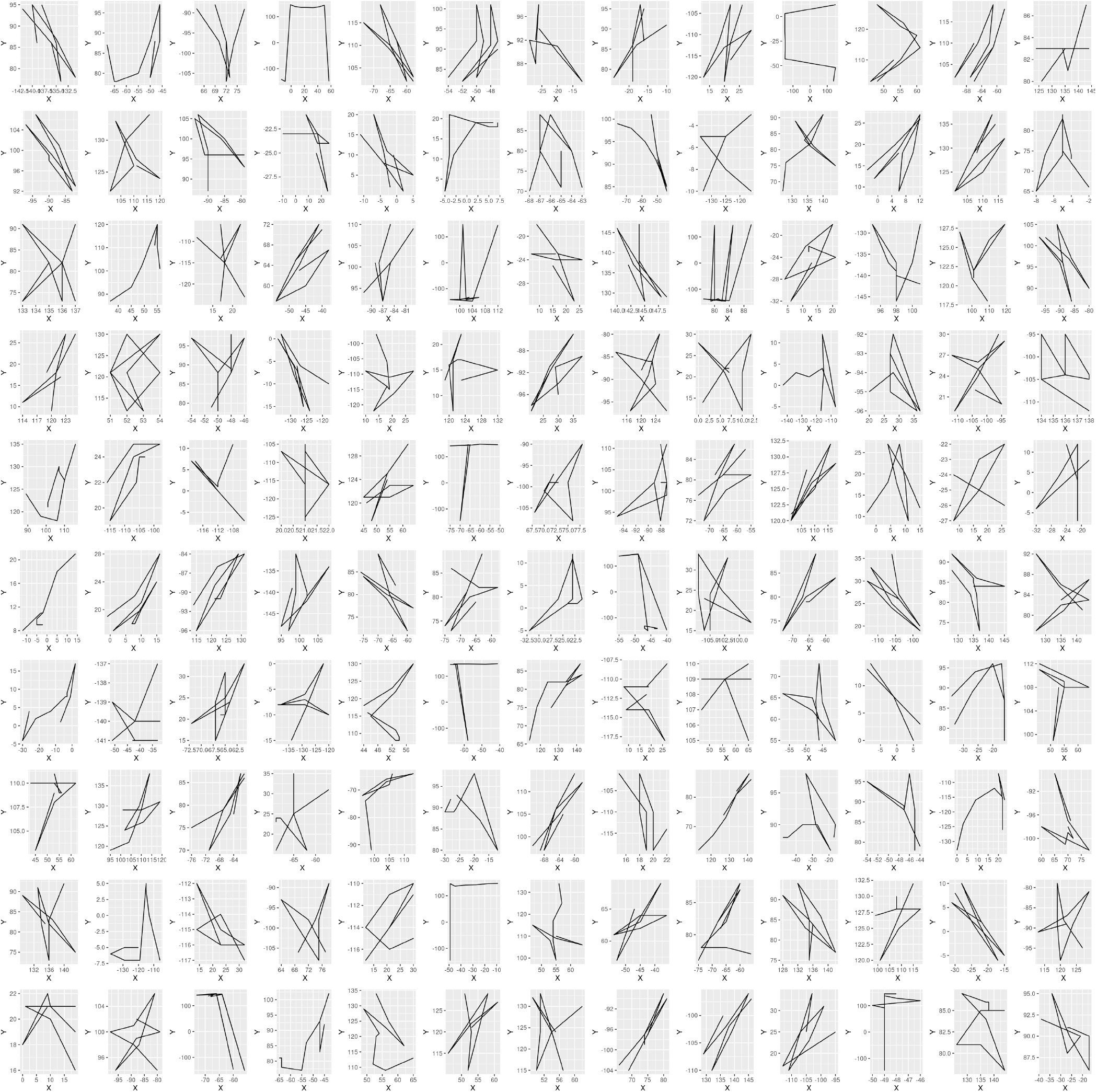
130 random 10-step segments from cluster C=7 (Table 1)

**Figure A.9:**
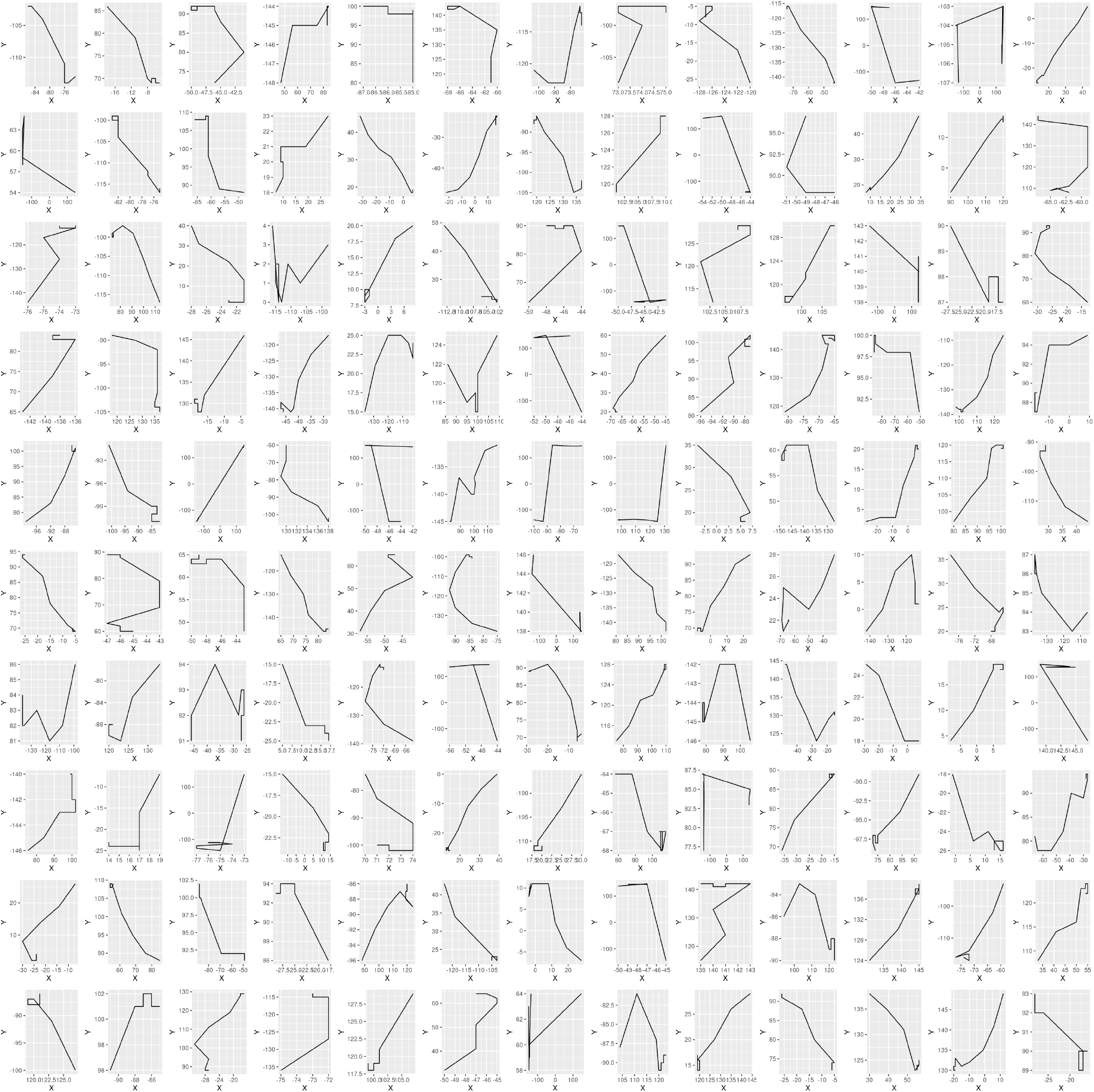
130 random 10-step segments from cluster C=8 (Table 1)

#### A.6 Plots of Adult Female Barn Owl StaMEs

Each plot in this subsection, particularly across different clusters, has had its axes for each panel automatically set by the plotting routine. Thus the size of segments is not comparable, only their relative shapes are informative.

**Figure A.10:**
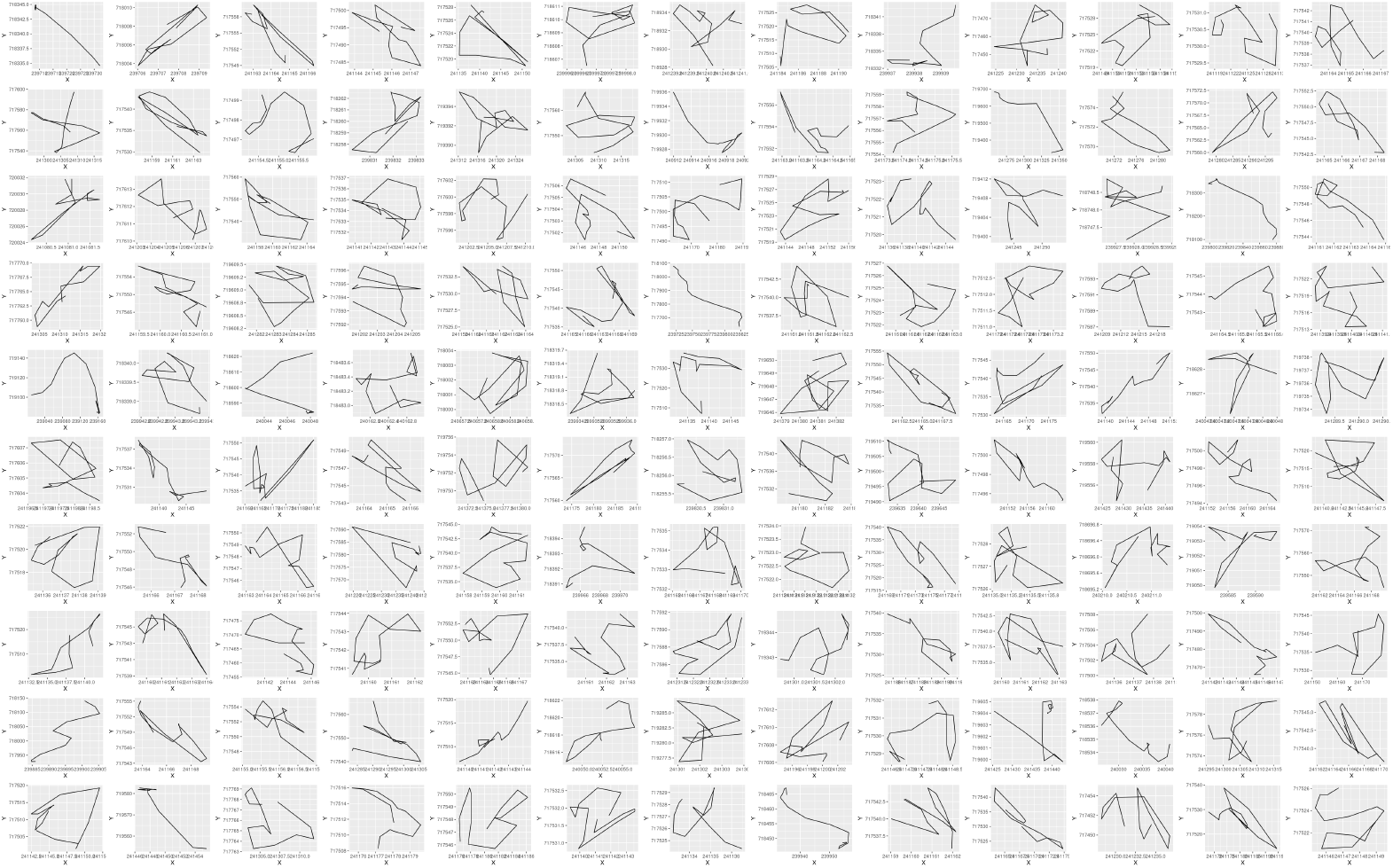
130 random adult female barn owl segments from cluster C=1 (Table 2)

**Figure A.11:**
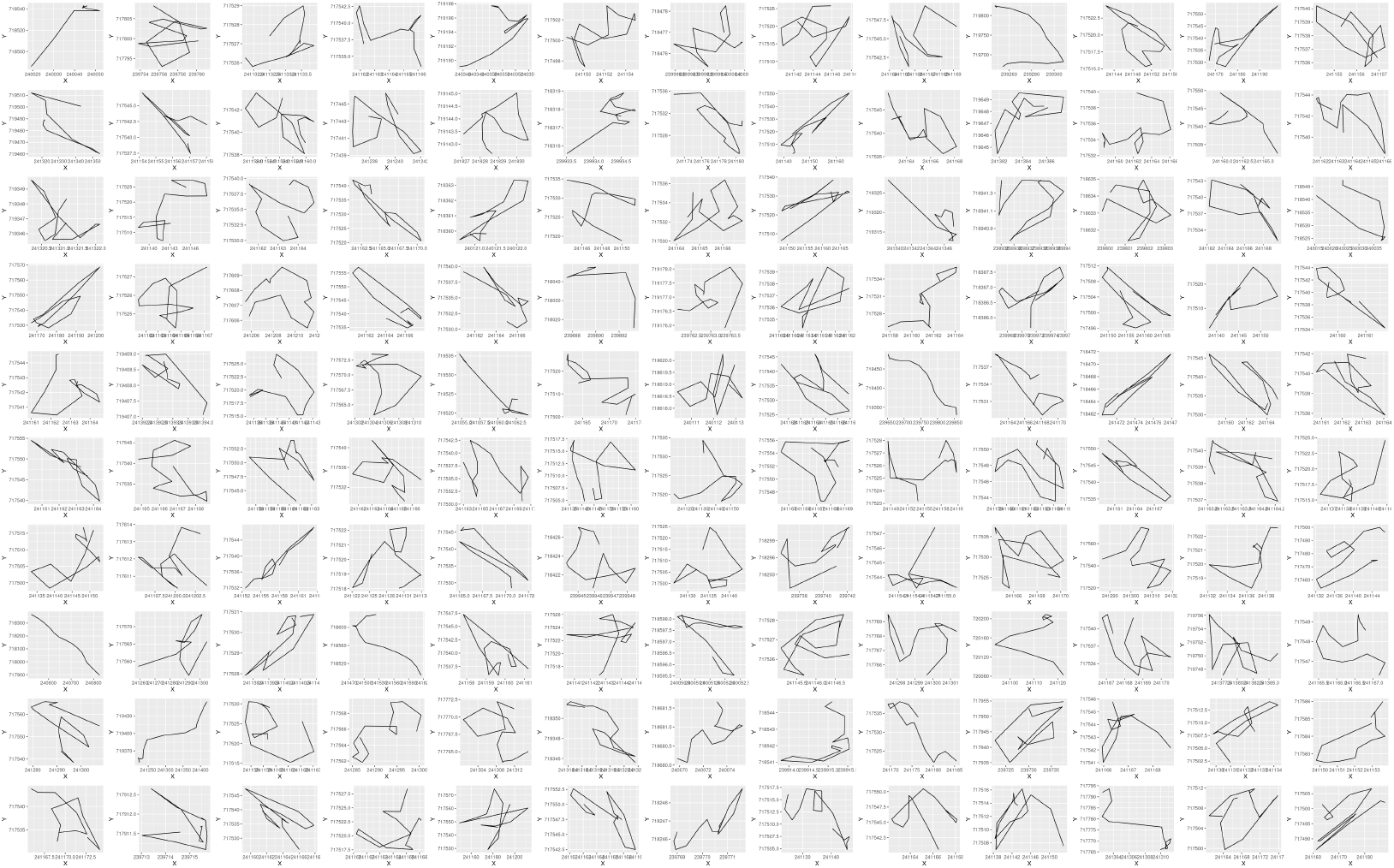
130 random adult female barn owl segments from cluster C=2 (Table 2)

**Figure A.12:**
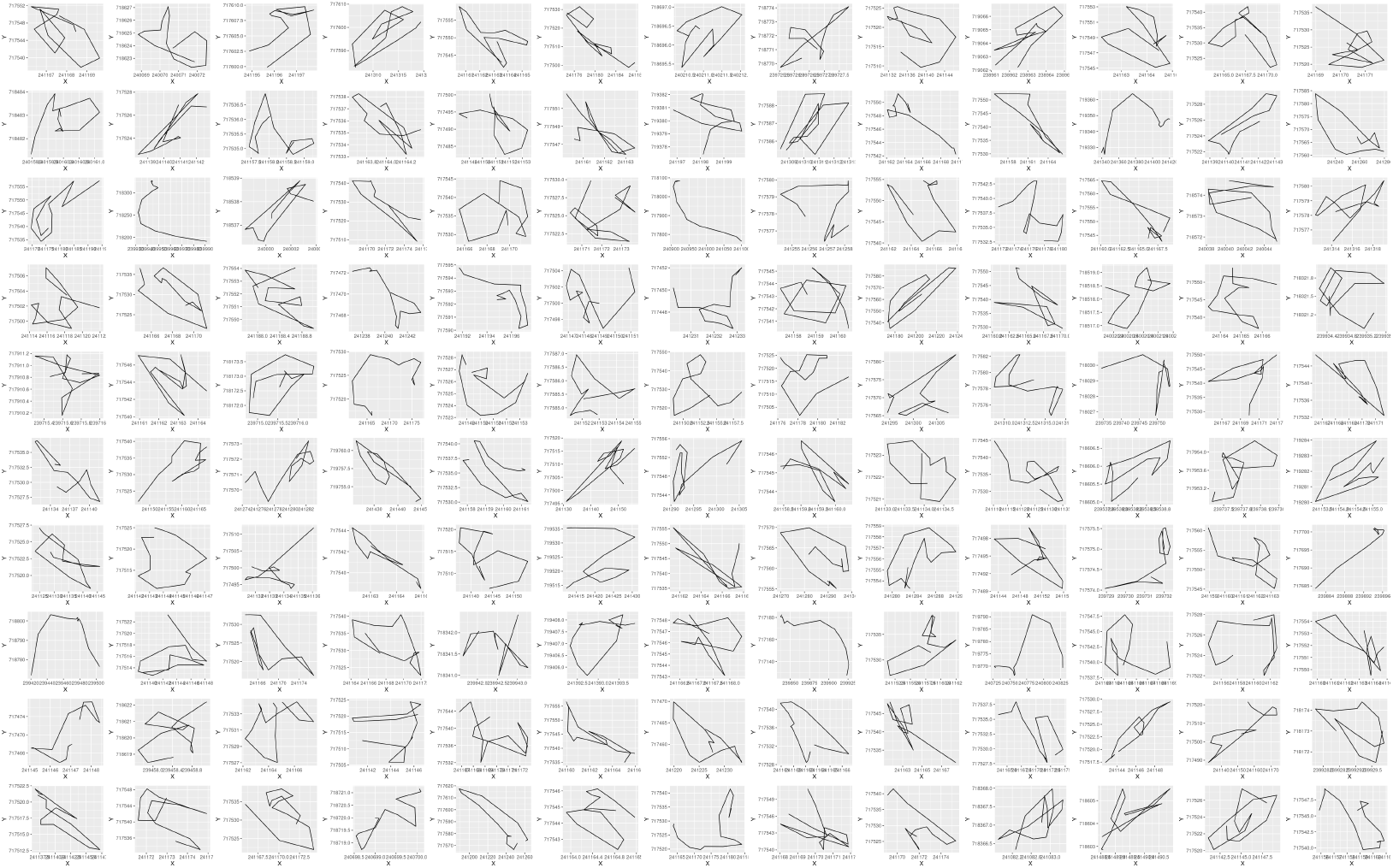
130 random adult female barn owl segments from cluster C=3 (Table 2)

**Figure A.13:**
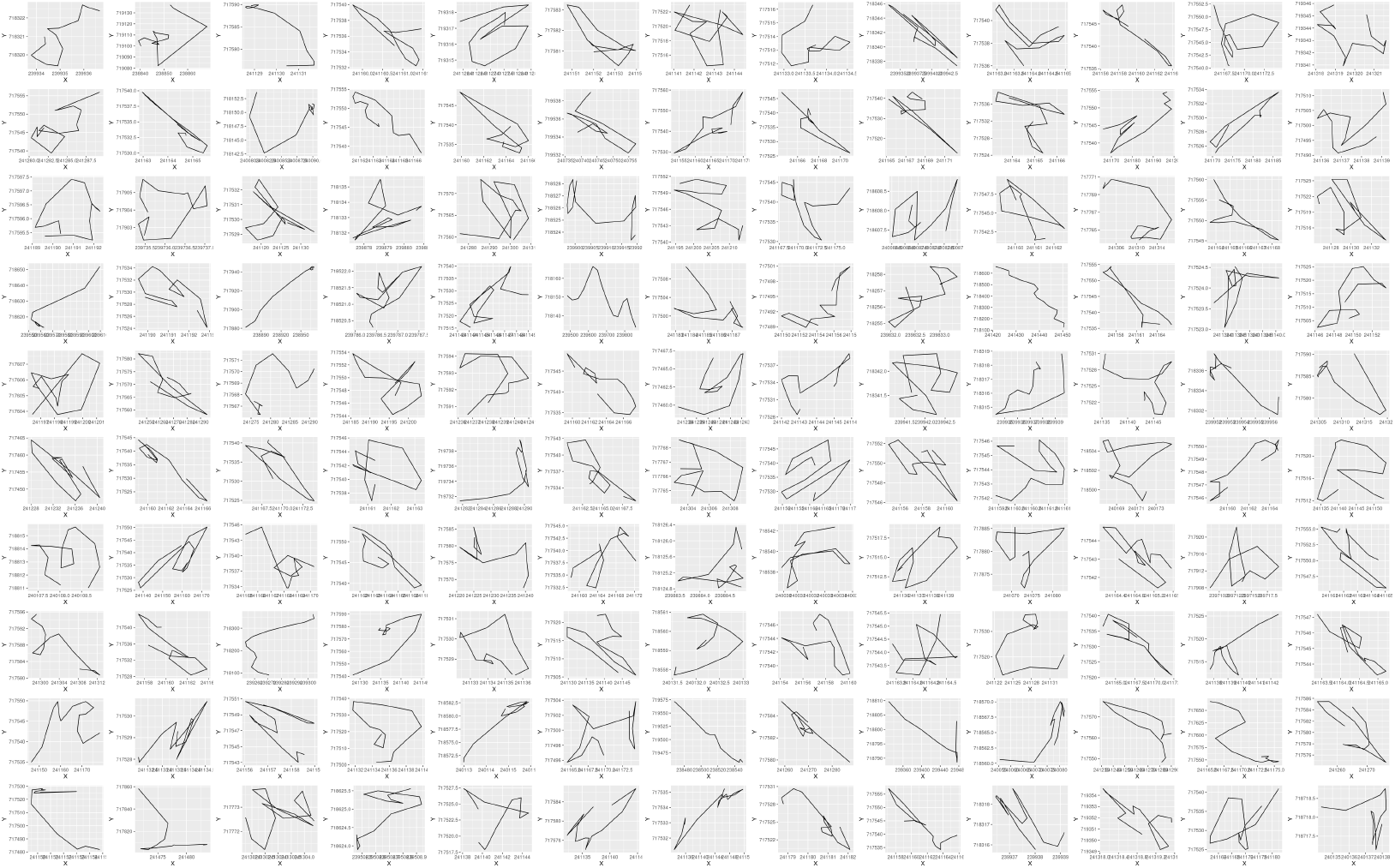
130 random adult female barn owl segments from cluster C=4 (Table 2)

**Figure A.14:**
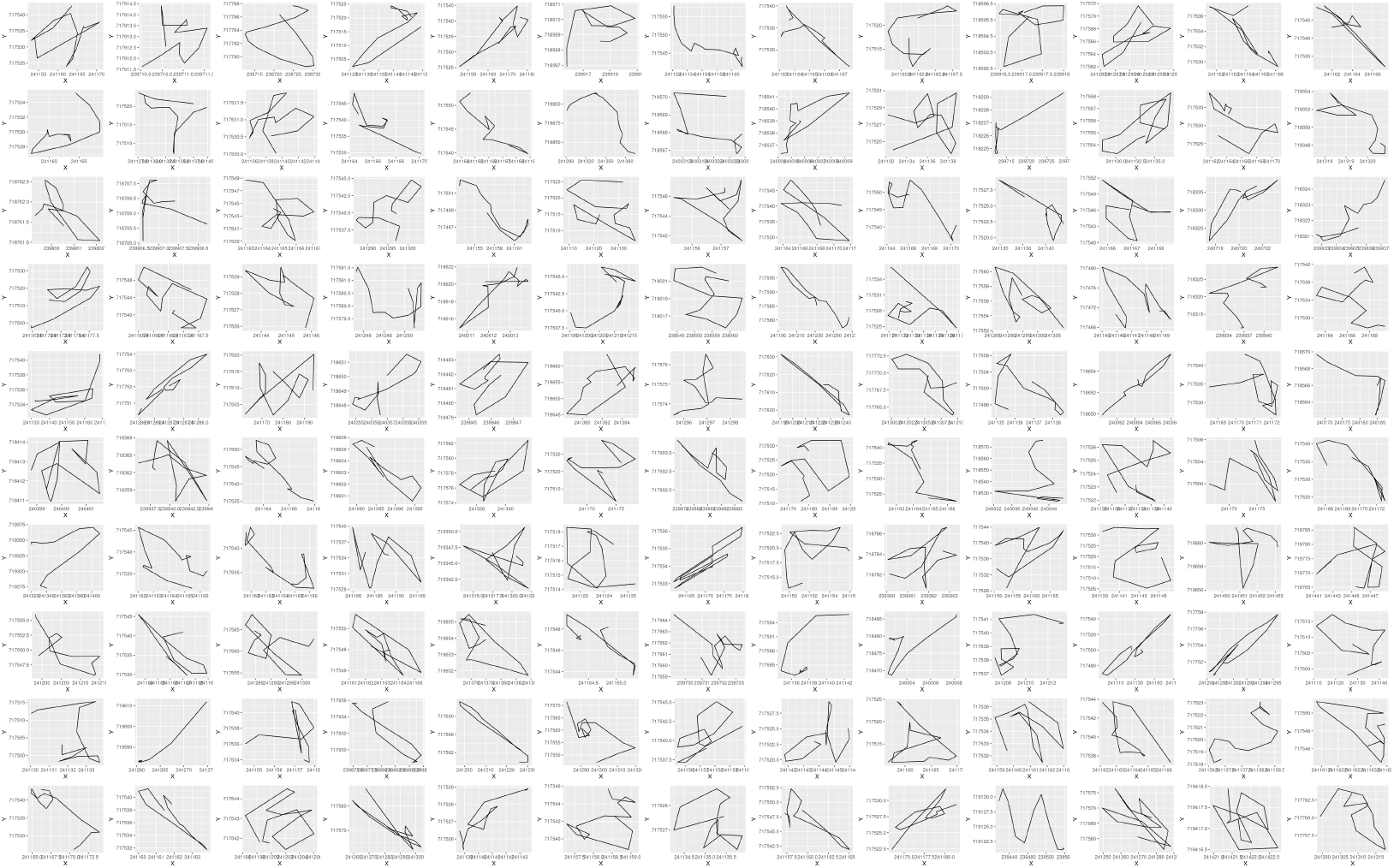
130 random adult female barn owl segments from cluster C=5 (Table 2)

**Figure A.15:**
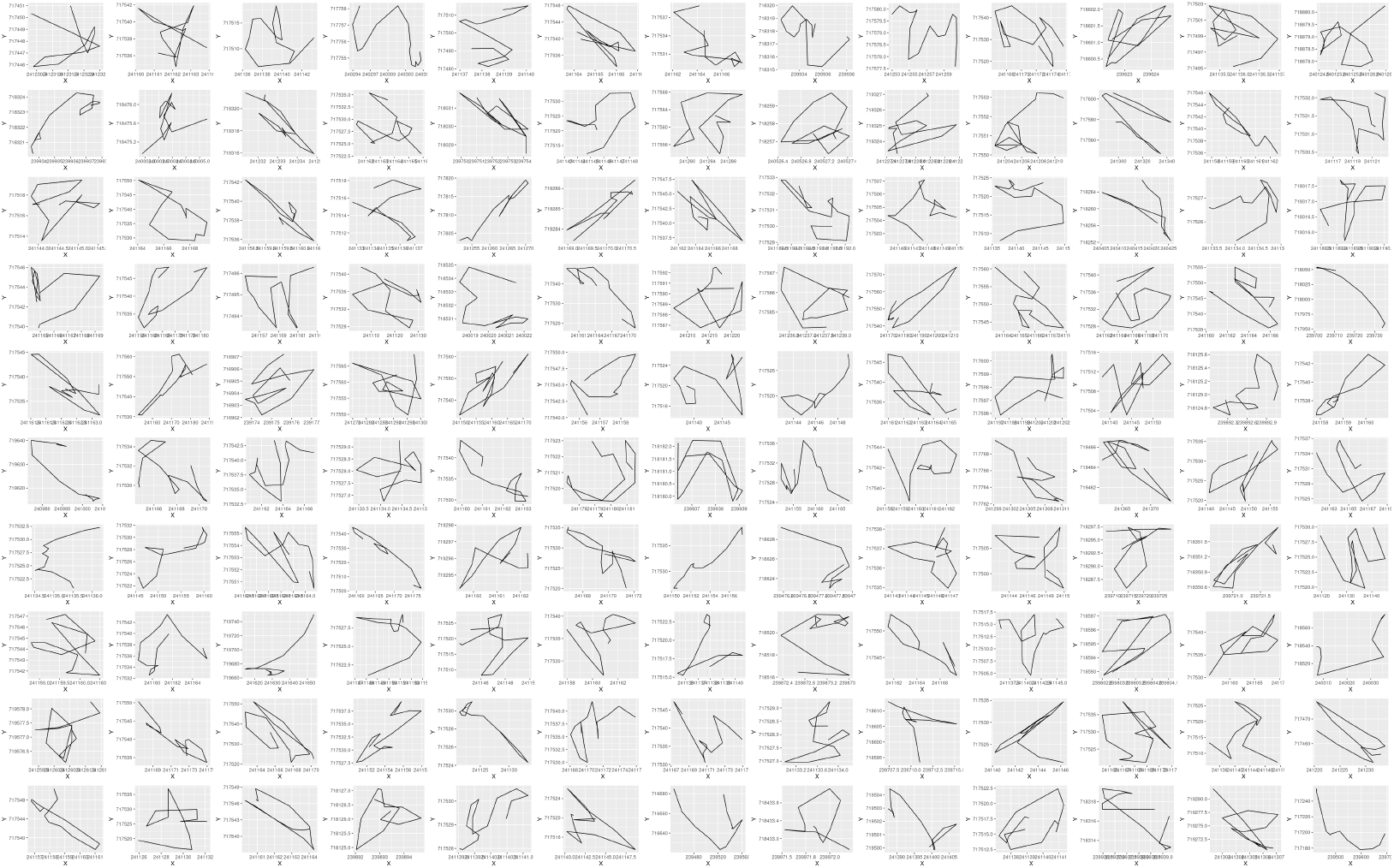
130 random adult female barn owl segments from cluster C=6 (Table 2)

**Figure A.16:**
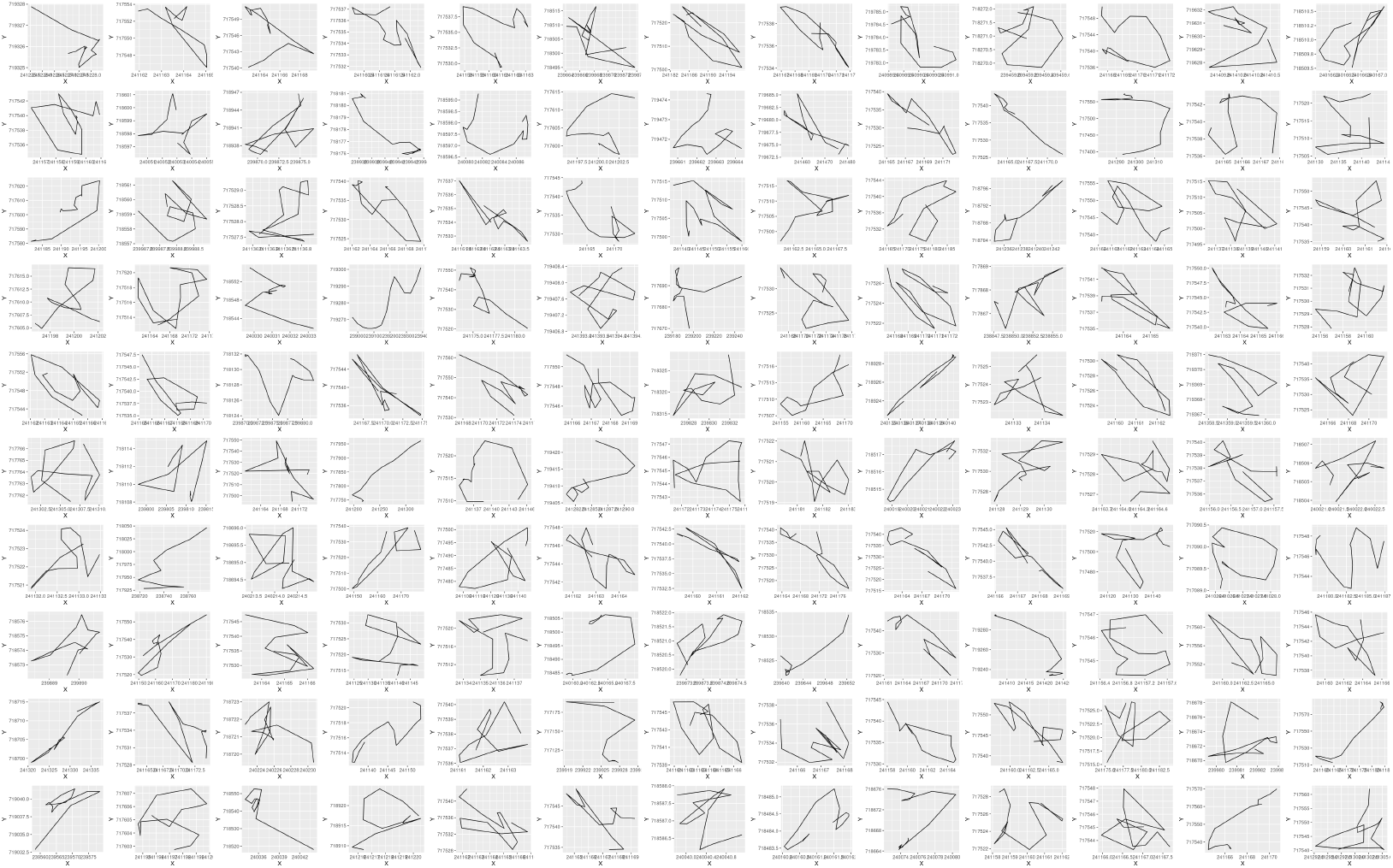
130 random adult female barn owl segments from cluster C=7 (Table 2)

**Figure A.17:**
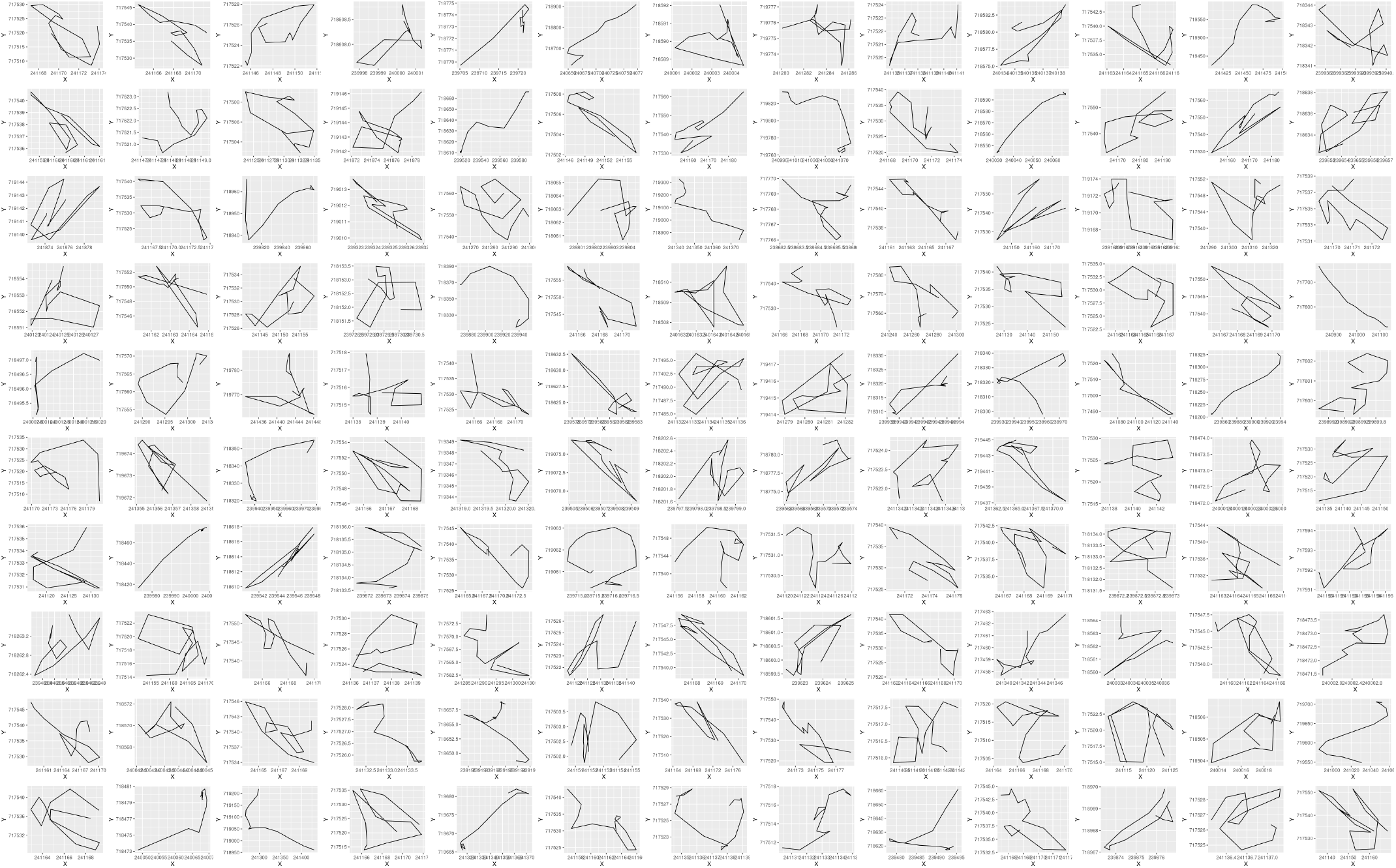
130 random adult female barn owl segments from cluster C=8 (Table 2)

#### A.7 Plots of Juvenile Male Barn Owl StaMEs

Each plot in this subsection, particularly across different clusters, has had its axes for each panel automatically set by the plotting routine. Thus the size of segments is not comparable, only their relative shapes is informative.

**Figure A.18:**
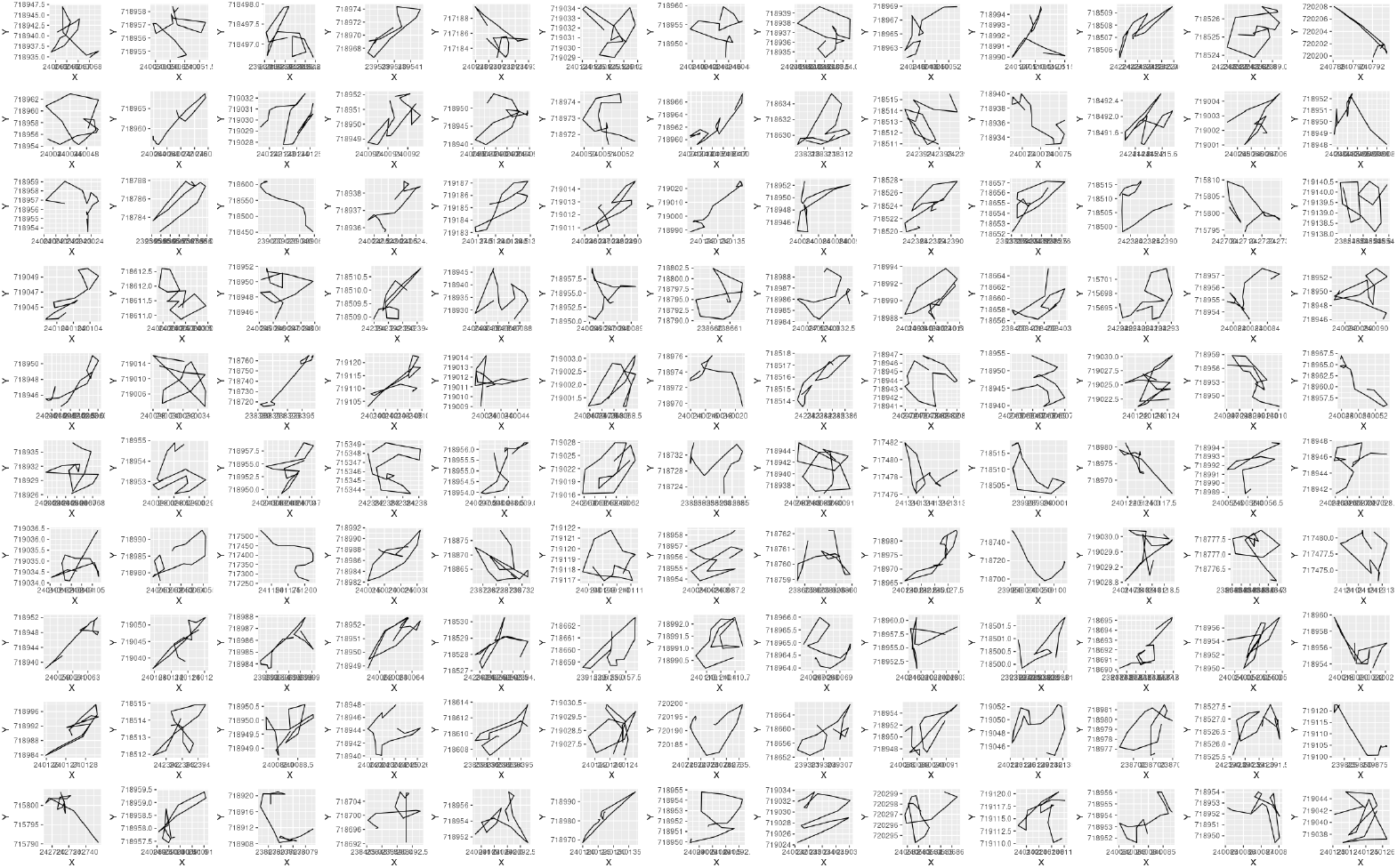
130 random adult female barn owl segments from cluster C=1 (Table 2)

**Figure A.19:**
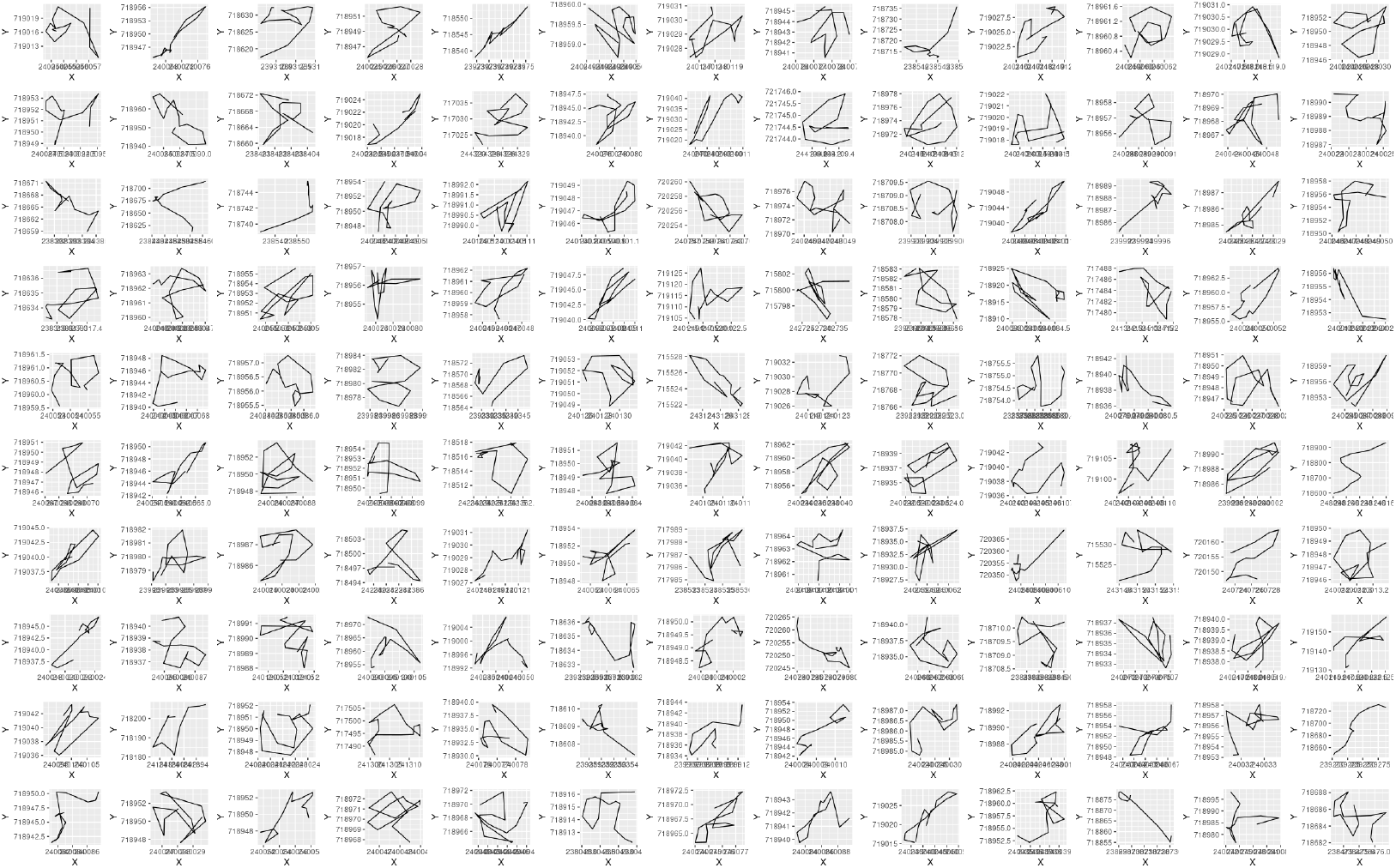
130 random juvenile barn owl segments from cluster C=2 (Table 2)

**Figure A.20:**
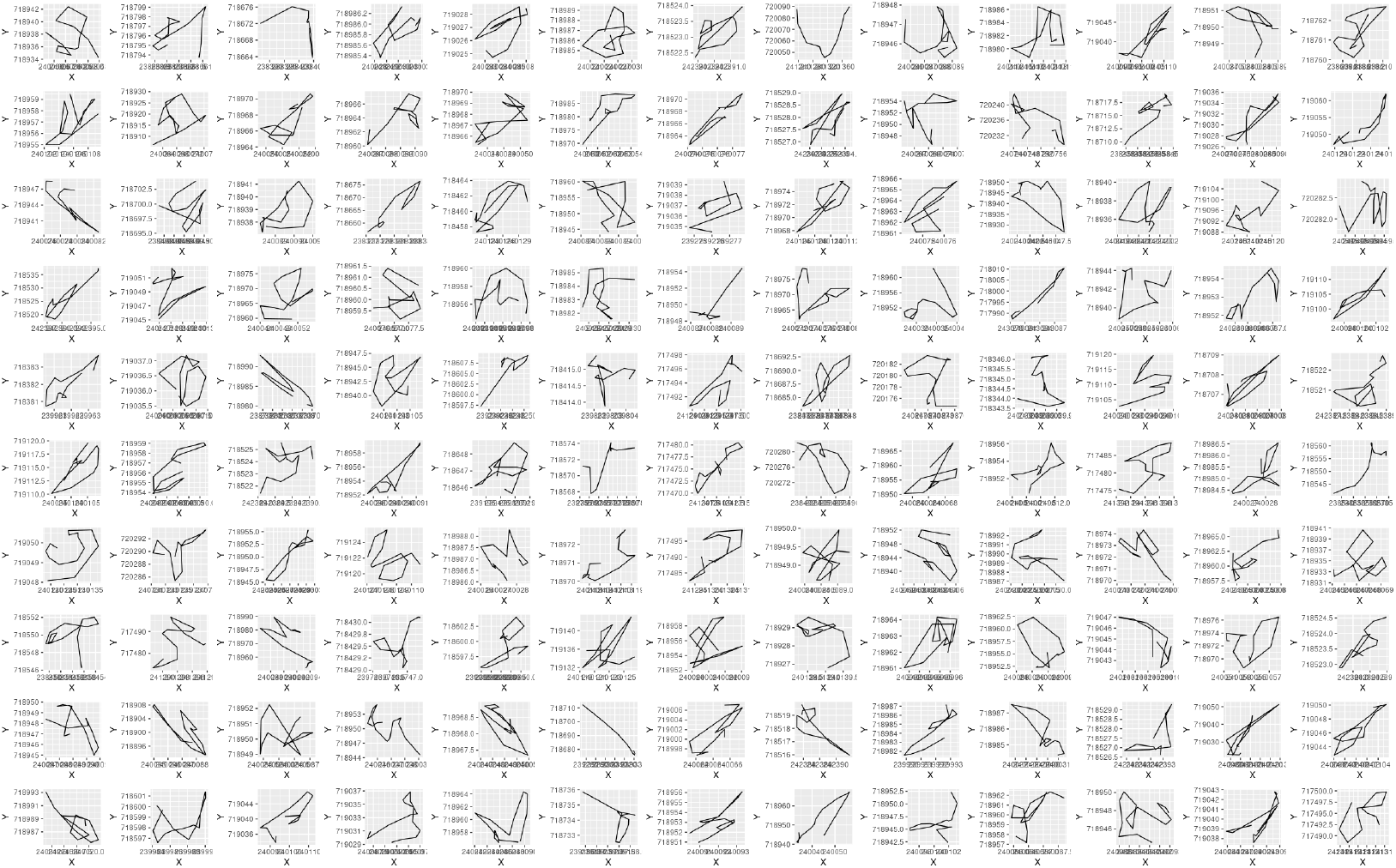
130 random juvenile barn owl segments from cluster C=3 (Table 2)

**Figure A.21:**
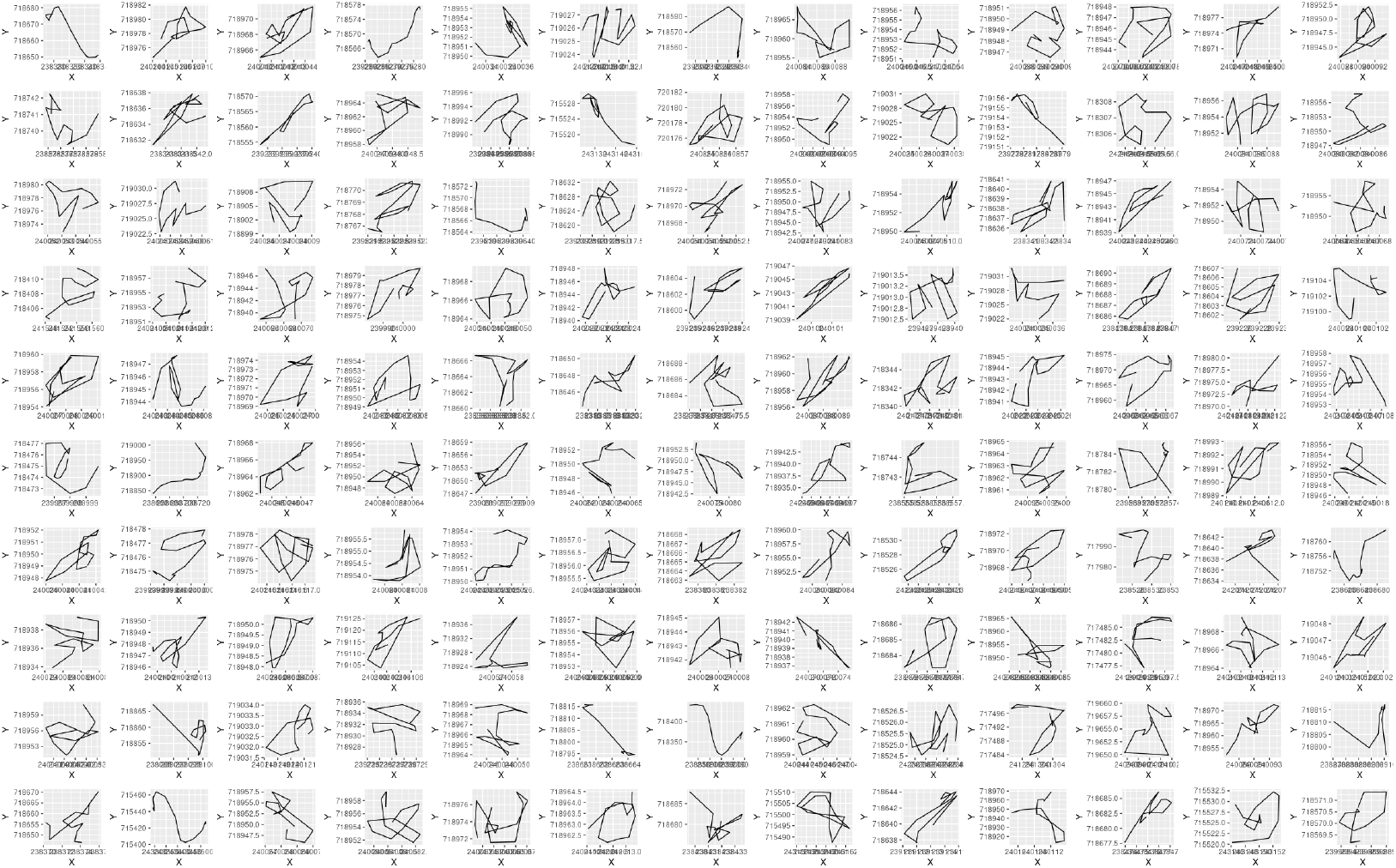
130 random juvenile barn owl segments from cluster C=4 (Table 2)

**Figure A.22:**
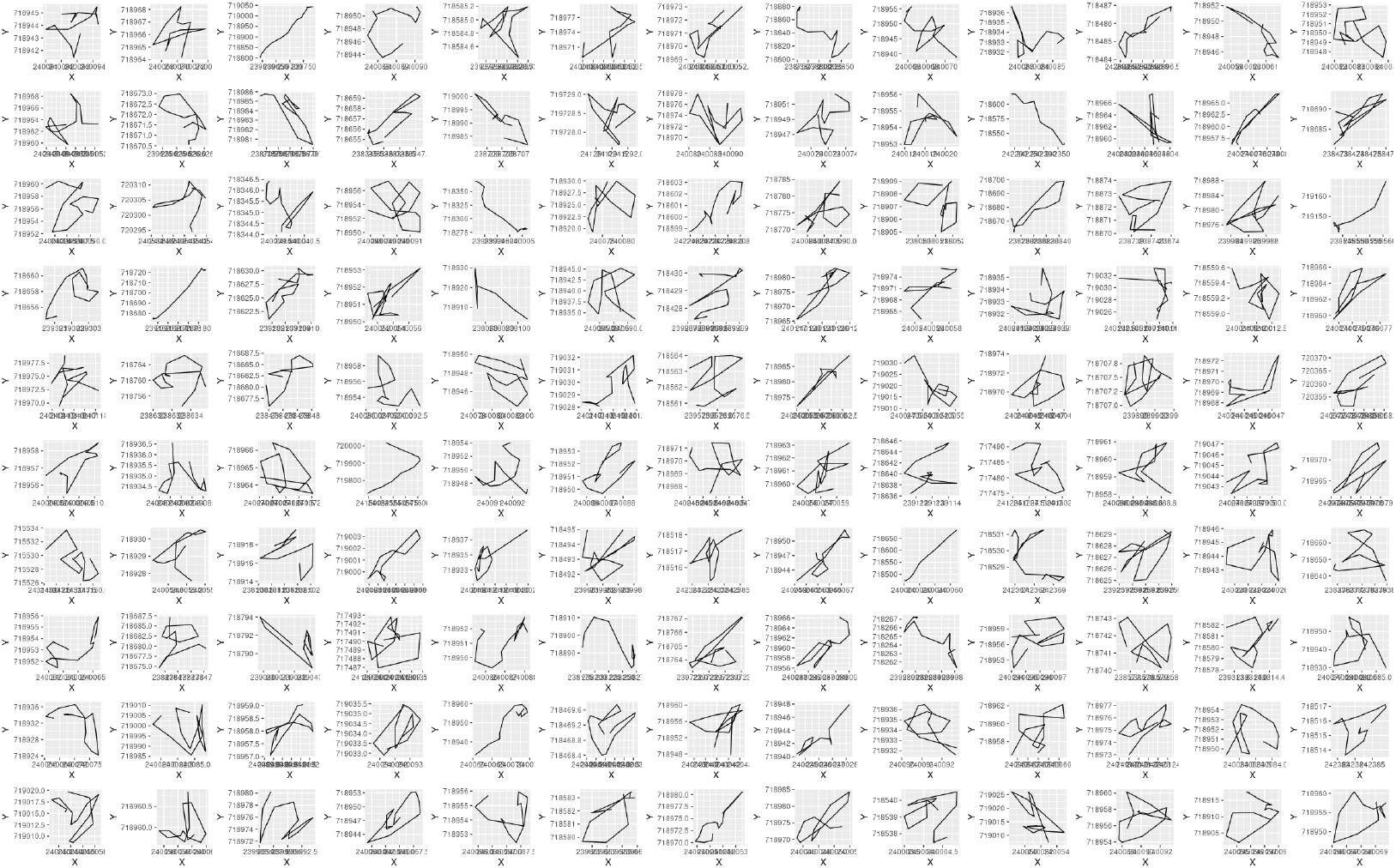
130 random juvenile barn owl segments from cluster C=5 (Table 2)

**Figure A.23:**
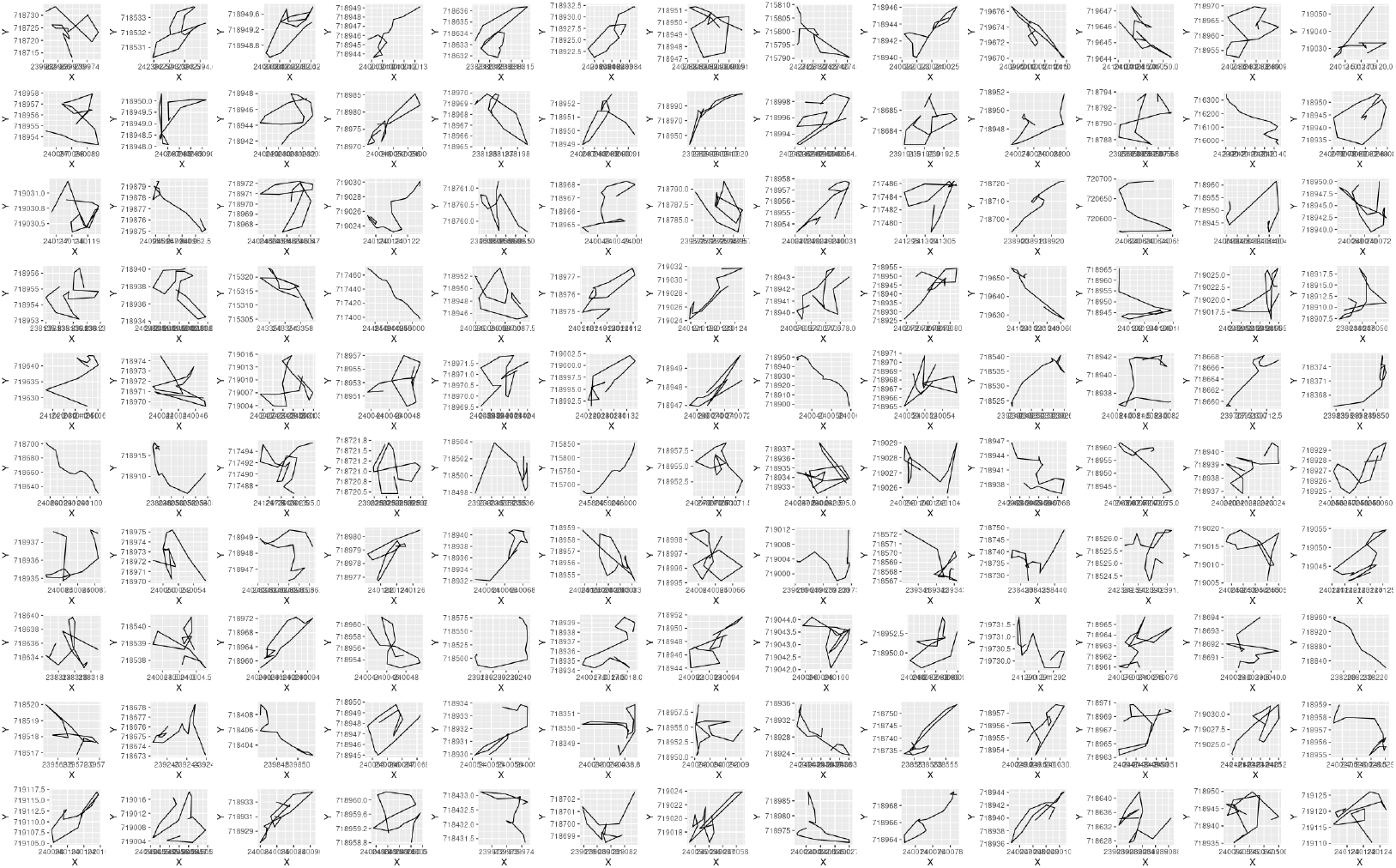
130 random juvenile barn owl segments from cluster C=6 (Table 2)

**Figure A.24:**
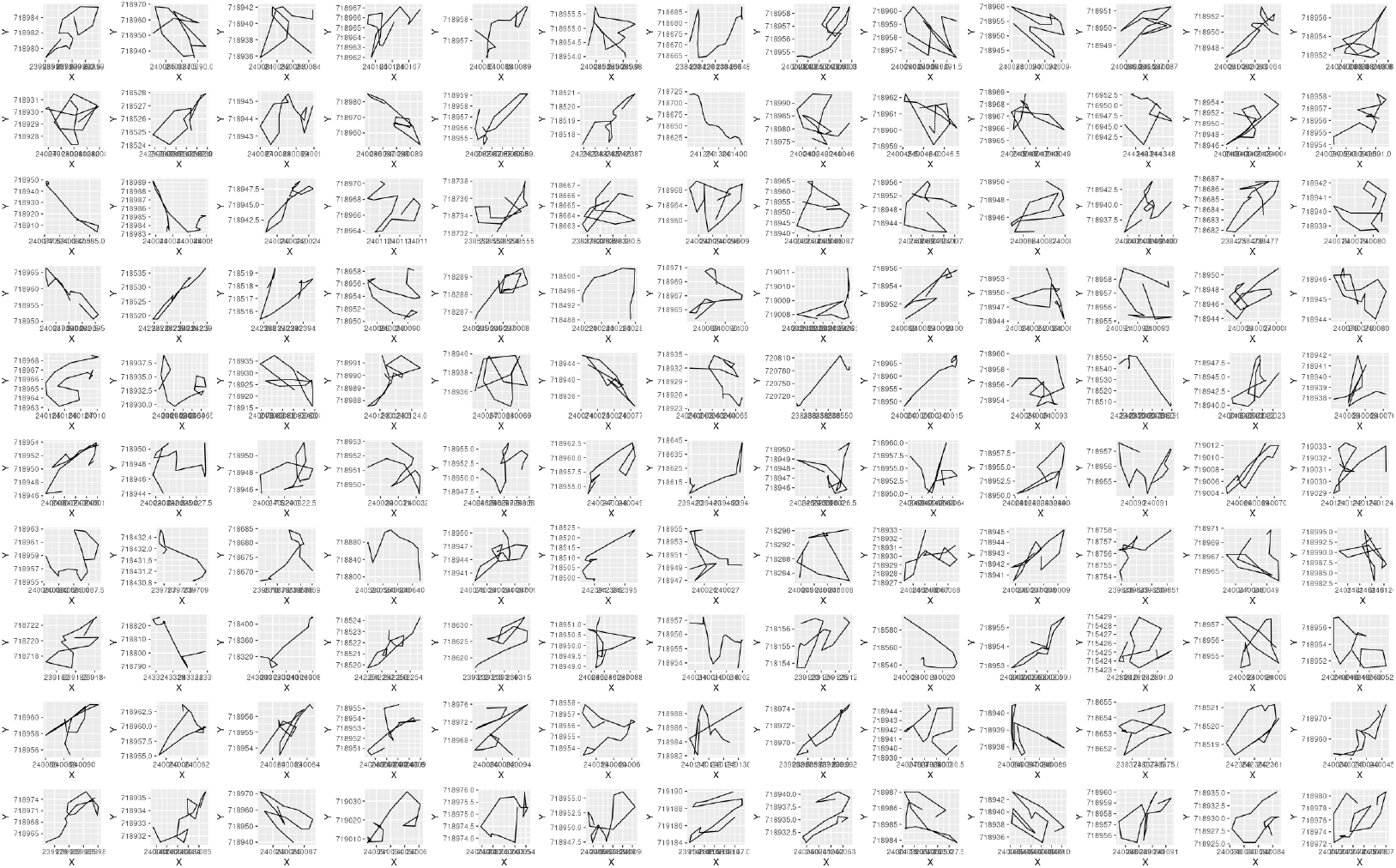
130 random juvenile barn owl segments from cluster C=7 (Table 2)

**Figure A.25:**
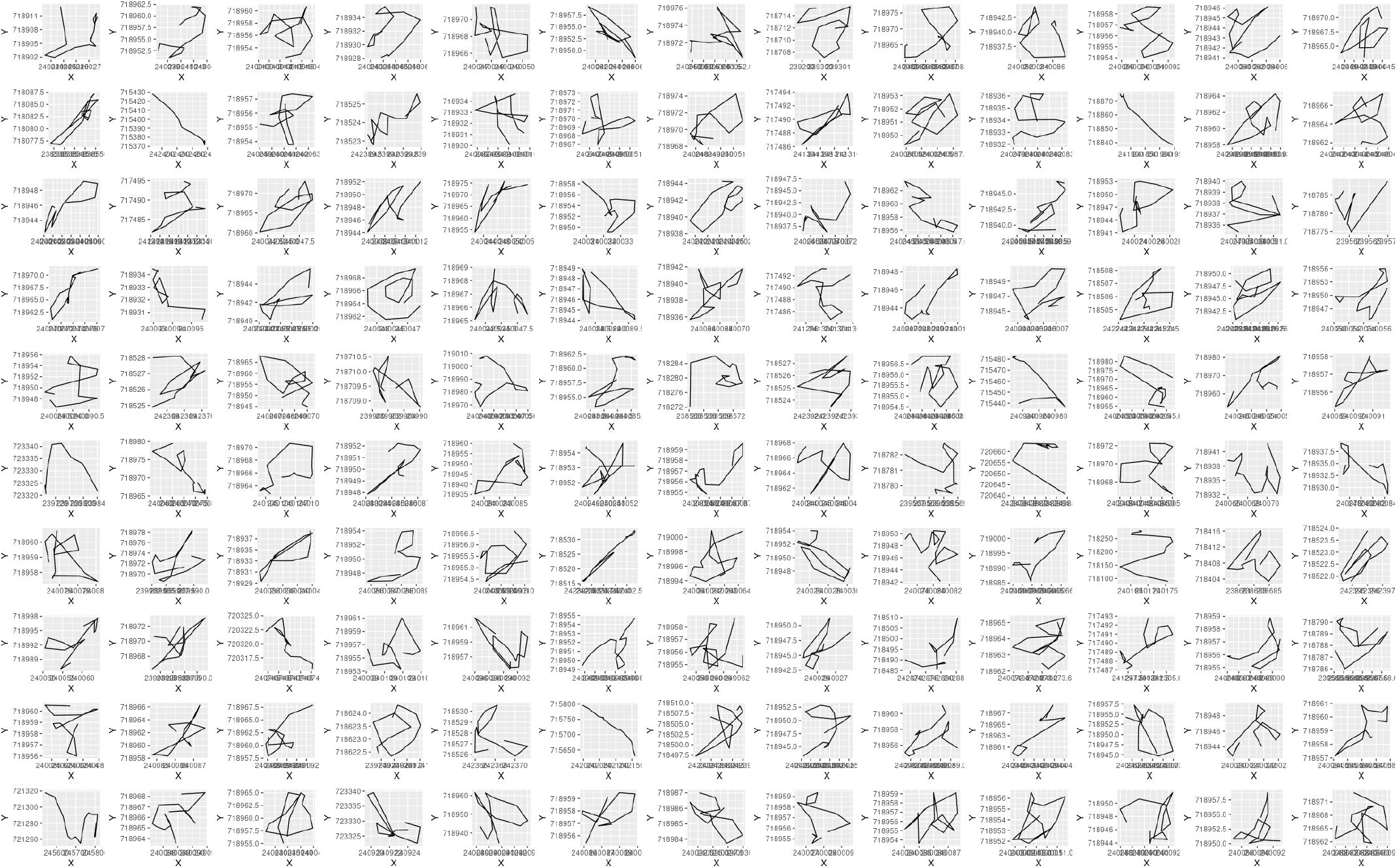
130 random juvenile barn owl segments from cluster C=8 (Table 2)

### B Numerical construction of a homogeneous movement map

A numerical representation of the mapping 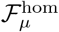 Eq 14, where we note that we have dropped the dependence on 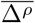 because of the lack of circular bias in our movement simulations and the index *μ* in the image space) can be constructed as follows:

1. For a set of kernels defined in terms a positive integer 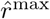 and an incremental angle Δ*ψ* (typically equal to *π/*12 or *π/*36) define the following set of kernels

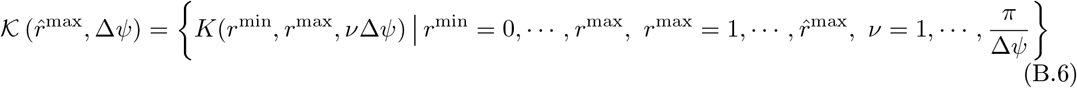
2. For each kernel in 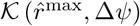 run a simulation of the model, moving the individual over the desired homogeneous landscape for a selected period of period of time. For example, if one wants to generate 100 segments, each 15 points long, then the simulation will proceed for 1500 time steps.
3. For each of the simulated trajectories of the kernels in 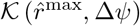, carry out a segmentation with segment size *μ* and generate the selected set of segment statistics, in our case the values *V*, SD^*V*^, |ΔΘ|, SD^|ΔΘ|^ for each segment. Compute the means of these statistics across the set of segments to obtain one candidate point of the mapping Eq 14.
4. In identifying the set of kernel (*r*^min^, *r*^max^, *ν*Δ*ψ*) that generates an image under the mapping 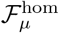 that is closest to the observed point 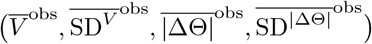 proceed as follows:
5. Discretize the image space of 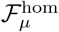 through a series of latin cube lattice computations of 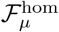 with arguments ranging over the cube with min and max bounding vertices 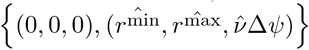 and intermediate cubes with min and max vertices 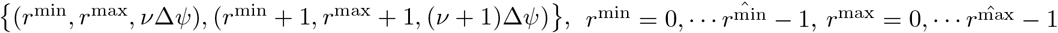 and 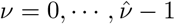.
6. Select the intermediate cube that contains the image 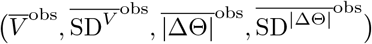: i.e., the cube for which, making the *K*_*α*_ arguments in 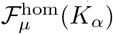 explicitly:

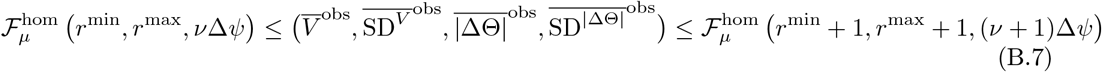
7. Either select the vertex in the cube defined by (*r*^min^, *r*^max^, *ν*Δ*ψ*), (*r*^min^ + 1, *r*^max^ + 1, (*ν* + 1)Δ*ψ*) whose image under 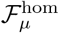 is closest to the observed point 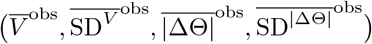 or use a finer resolution Latin Cube grid within this cube to locate a set of parameters the provides an image as close as desired to the observed point.

### C ANIMOVER_1: Access and Data

#### C.1 Output Data

At the end of the simulation the output data can either be automatically or manually saved (using a toggle switch on the console, see Fig 5, N in main text) as a csv file. The header to this file contains a list of all the parameter and switch settings that were used to generate the run. In addition, the data are listed in 8 different columns depicted in Fig. C.1.

**Figure C.1:**
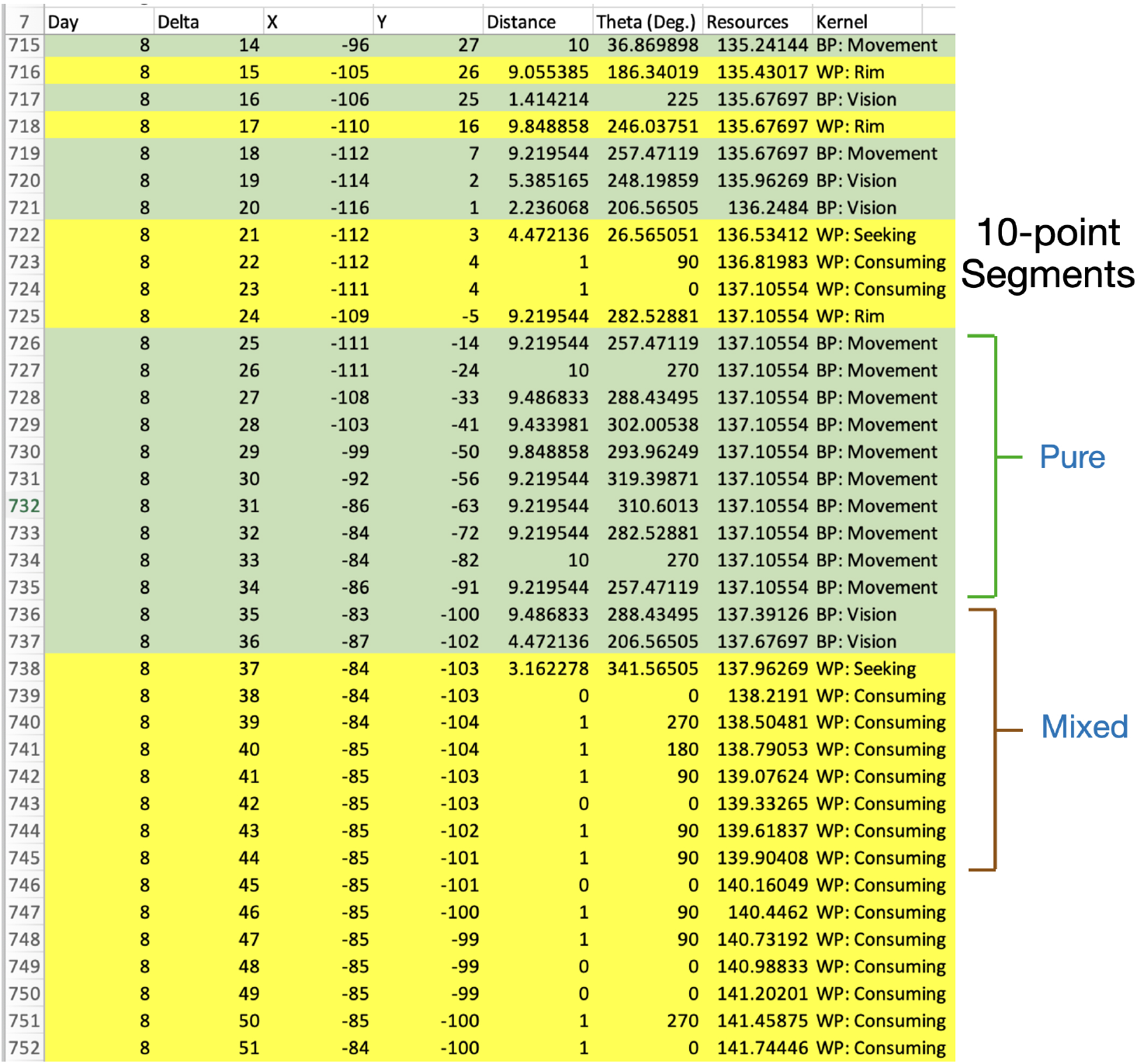
The 8 columns in the csv data output file (e.g., see the file `Two_Kernel_Movement.csv`, Supplementary Data) that can be saved at the end of the simulation as as follows: `Day`, `Delta` (within-day step), `X` (*x*-location), `Y` (*y*-location), `Distance` (distance moved), `Theta` (angle of heading in degrees), `Resources` (agent-state), `Kernel` (BP or WP). We note that one computes the turning angle Δ*θ* at time *t* by subtracting the angle of headings at time *t* from the angle of heading at time *t −* 1 taking into account that angles are specified modulo 360 degrees (e.g., if *θ*_*t−*1_ = 350 and *θ*_*t*_ = 10, then the turning angle is Δ*θ* = 10 *−* (350 *−* 360) = 20 degrees). Thus angles have heading range over [*−π, π*]. In addition, the third argument in our StaME is the average of the absolute values of the angles of heading rather than just the angles themselves. Also note that the BP movement mode (green entries) has two states: `Movement` (movement within the kernel rim which rim exists whenever 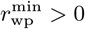) or `Vision` (movement within the kernel sector, possibly between the inner rim radius and the current location). The WP movement mode has three states: `Consuming` (when moving to a resource-rich location), `Seeking` (an interim step in looking for a resource rich location when the kernel contains insufficient resources), or `Rim` moving the maximum step length (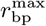 rather than 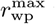 when the seeking state fails to provide a suitable resource location). The indicted 10-step segmentation of the data shows how both pure and mixed movement segments arise during segmentation.

#### C.2 Downloading and Running the App

ANIMOVER_1 and the most recent release of Numerus Studio can be downloaded without cost from the Numerus webpage https://www.numerusinc.com/studio/. Installers for Numerus Studio are provided for Mac and Windows platforms. Instructions for using Numerus Studio are contained in the RAMP Users Guide at https://wiki.numerusinc.com/index.php/Ramp_User_Guide

ANIMOVER_1 is deployed in the RAMP file `Ani1Cr3.nms`. After installing Numerus studio, open this file and launch it from the Studio launchpad. Documentation for this RAMP can be found at https://wiki.numerusinc.com/index.php/Animover_1.

### D ODD Protocol for ANIMOVER_1

The element numbering scheme used in the subsections below is that of the second ODD revision present in the Grimm et al. [63].

#### D.1 Purpose and Patterns

The reason for developing an animal movement simulator is articulated under goal c.) in the Introduction of the main text. More broadly, we construct a user friendly, highly flexible movement track simulator that can be used to test ideas and concepts. This includes testing hypotheses about underlying mechanisms that may lead to particular emergent patterns of movement behavior [111]. In addition, simulated data can be used, as we did to a limited extent in this paper and now do more extensively in our ongoing research, to evaluate methods for movement track analysis, and forecasting animal movement patterns in changing and novel environments. Although our simulator is built at two scales—1.) next-step decisions of where to move next using ideas from step-selection function analysis [54], 2.) time-already-spent-in-current-movement variables (e.g., mimicking satiation, increasing thirst, or hunger effects) and time-within-diel-cycle variables (e.g., when to head home)—patterns emerge at a third scale (different kinds of diel activity routines [42]), with higher scale seasonal patterns emergent as well.

#### D.2 Entities, state variables,and scales

The relevant entities and state variables are listed in Table 3. They are cells (as represent by their position and associated euclidean location in the cellular array 𝒜), the resource states *c*_*ab,t*_ of cells at time *t*, the individual agent’s location 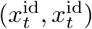, movement mode *α*_*t*_, angle of heading *θ*_*t*_, time in current movement mode 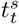, and internal state value *h*_*t*_ (e.g., level of resource satiation or its inverse, hunger).

#### D.3 Process overview and scheduling

The process overview and scheduling are illustrated in Fig 3, which depicts the computational flow sequence of the movement decision algorithm at the core of ANIMOVER_1. In short, after selecting parameter values and setting up the initial patch structure of the landscape (details next), the algorithm loops through a next-step process. This involves 1.) computing the location to which the individual moves from its current location based on the individual’s current movement mode and the summed resource state of the cells in its current neighborhood, 2.) computing the amount of resources the individual extracts from the location to which it moves, thereby updating its current state to include these new resources and the cost of movement, 3.) updating the resource state of the cell to which the agent moves and from which it extracts resources, 4.) updating the current movement mode based on the state of the neighborhood of the agent’s current location, 5.) implementing the STOP rule when either the end of the simulation has been reached (*t* = *T*) or the individuals internal state has hit 0 (*h*_*t*_ = 0; the individual is now dead).

##### D.3.1 Patch setup

The process for setting up the initial patch structure of the landscape is implemented by the runtime alterable module 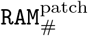, where the details of the different possible preselected versions # = 0 (default), # = 1 (selectable alternative), #2 (for use to supply customized code) are elaborated in Section 4.2 of the main text.

##### D.3.2 Resource extraction

The process for computing the amount of resource extracted from the current cell in which the agent is located and the new state of the agent as a function of the amount of resource it extracts is implemented by the runtime alterable module 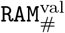, where the default version 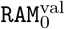 uses the resource density independent equations, Eq 6, and the alternative version 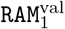 uses the resource density-dependent equations, Eq 7. It is also possible for the user to insert a customized set of equations by coding their own version 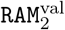 of this RAM.

##### D.3.3 Next step computation

The next-step computations are implemented by the within-patch and between-patch step selection procedures 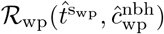 and 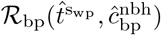 provided in Appendix A.3 above.

#### D.4 Design concepts

In the ODD protocol, this sections describes how the following 11 concepts were considered in the model.

##### D.4.1 Basic Principles

The basic principle behind our movement algorithm is that an individual chooses were to move next based on its current mode of movement (e.g., is it searching for resources or commuting to a new location), the statistics of the step-size and turning angles associated with its current movement mode, and the best location to land on its “next-step” given the state of the landscape within a radius of the largest value associated with the distribution of step sizes for its current movement mode.

##### D.4.2 Emergence

Emergence relates to patterns produced by a sequence of steps associated with the current movement mode, and by switching several times among movement modes to produce patterns at the scale of BAMs (e.g.: resting that emerges from a sequence of small, directionally random StaMEs; foraging that emerges from a mixture of medium and short StaMEs with relatively high turning angles; commuting that emerges from a sequence of large StaMEs with relatively low turning angles), DARS (e.g., commuting to a known distant location where feeding occurs and then returning versus interspersed searching and resting close to a home location), LiMPs (e.g., migration, ranging, or territoriality phases) and LiTs (a central place forager, versus a ranger, or an annual migrator; also see Fig 1 in main text).

##### D.4.3 Adaptation

Adaptation of movement is implicit in the model, first in the way movement kernels are impacted by the current landscape structure (see Fig 2C) and second in the fact that the state equations, depending on the particular 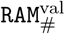 used (see above), contain feedback structures that result in density-dependent resource extraction. Additionally, we note that as the simulation progress, resource consumption by the agent during the early phases of the simulation reshapes the landscape and hence my lead to shift in the agent’s emergent movement patterns over time.

##### D.4.4 Objectives

From the agents point of view, the objective is to maximize resource intake at each within patch movement step. This is realized through rule ℛ_wp_.2 (Appendix A.3) where the maximum value 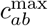 is computed and decisions where to move next are based upon the location of the cell that has this value.

##### D.4.5 Learning

An ability to adapt through learning is not included in ANIMOVER_1.

##### D.4.6 Prediction

As mentioned under element 1 (D.1 above), “forecasting animal movement patterns in changing and novel environments” is one of the purposes of the model. The ability of the model to reliably predict where an animal moves next, as well as forecast patterns of movement at larger scales requires that the parameters of the model first be fitted to the data using an estimation procedure suitable for the task in the context the type of model and data available [112]. This has not been done here since the third goal of this paper is both make available the ANIMOVER_1 simulation for the purpose elaborated in element 1 (D1) above and also, as articulated in the Introduction of the main text. Model parameter estimation is data specific and so will need to be undertaken in any studies that involve using ANIMOVER_1 to predict the movement of individuals by fitting kernel and state updating parameters to empirical data on both the movement and state of those individuals.

##### D.4.7 Sensing

An agent sensing the state of the environment is a very important component of ANIMOVER_1. This is done through the definition of movement-mode specific full and rim circle-sector kernels (Fig 2). Looking closely out the output in Fig C.1, in the last column (“Kernel”) 5 types of kernels are identified in the context of between-patch (BP) and within-patch (WP) movement. Specifically we see BP:Movement, BP:Vision, WP:Seeking, WP:Consuming, and WP:Rim. The BP:Movement and WP:Seeking are just the next-step between-patch and within-patch selection kernels. The BP:Vision kernel is invoked when an individual hits a landscape boundary during between-patch movement and thus needs to look all around (including behind itself) to make its next move (see ℛ_bp_.2 in Section A.3). The WP:seeking kernel is invoked with an individual within a patch does not find a cell to move to that has resources that exceeds its threshold requirements in which case it enlarges and fills out its current movement rim to a full movement sector (see ℛ_bp_.4*c* in Section A.3. If this kernel then fails to find a suitable cell for its next step, it moves at random to the best cell available in the kernel WP:Rim.

##### D.7.8 Interaction

In ANIMOVER_1 is a single-agent simulator and interactions are confined to a resource-consumption process involving the resources within the cell where the agent is currently located, according to Eqs 6 and 7. Clearly a multi-agent simulator can be developed where interactions among individuals with respect to both movement (either attraction or repulsion elements can be included) and resource exploitation (competition for resources among individuals in the same cell can be introduced into Eqs 6 and 7 using approaches such as those discussed in [53]

##### D.4.9 Stochasticity

Stochasticity enters into several different places in the simulation. First, the process of setting up patches in `RAM`^patch^ involves laying down a set of patch seeds and then randomly adding neighbors to these seeds with a certain probability, as articulated in Section 2.2 of the main text. Second, movement to a particular cell in the within and between patch movement procedures ℛ_wp_ and ℛ_bp_ involves a multinomial drawing in which cells higher resource cells falling within the movement kernel in operation at time are more likely to be selected than lower resource cells.

##### D.4.10 Collectives

Not applicable to ANIMOVER_1.

##### D.4.11 Observation

It is assume in ANIMOVER_1 that the simulate locations of the agent as it moves over the landscape are observed without error. If observation errors need to be introduced into this process, then noise can be added to the output in a form desired for the analysis hand.

#### D.5 Initialization

Initialization of the landscape patch structure is dealt with in Section 2.2 of the main text. The process for initializing other model parameters is discussed in Section 4.3, P6 and P7.

#### D.6 Input data

The process for initializing the landscape structure by reading in an input file is discussed in Section 4.3, as is the process for setting all the parameter values for the run.

#### D.7 Submodels

##### D.7.1 Landscape construction

This is handled by `RAM`^patch^, as discussed in D3.1 above.

##### D.7.2 Resource extraction

This is handled by `RAM`^val^, as discussed in D3.2 above.

##### D.7.3 Next step computation

This is handled by algorithms (step selection rules/procedures) 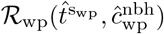 and 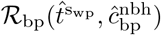 provided in Appendix A.3.

## Notes

https://www.numerusinc.com/studio/

https://wiki.numerusinc.com/index.php/Ramp_User_Guide

https://wiki.numerusinc.com/index.php/Animover_1

### Competing Interest Statement

The authors have declared no competing interest.

### Summary of Updates

This version has a revised abstract, a couple of pages of additional text, and additional column of results in Table 1, all added after receiving reviewers comments.

